# MLL1 complex is a critical regulator of fetal hemoglobin repression

**DOI:** 10.1101/2025.03.24.645036

**Authors:** Yufen Han, Bjorg Gudmundsdottir, Kristbjorn O. Gudmundsson, Kartik R. Roy, John Tisdale, Yang Du

## Abstract

Increasing fetal-type hemoglobin (HbF) expression in adult erythroid cells holds promise in the treatment of sickle cell disease (SCD) and β-thalassemia. We have identified MLL1 complex as a critical regulator of fetal and embryonic hemoglobin repression. Knockdowns of *MEN1* and *KMT2A,* encoding essential components of the complex, caused a significant downregulation of *BCL11A* expression and a substantial increase in γ- and ε-globin mRNA levels in HUDEP-2 cells. Significant binding of MEN1 and KMT2A were readily detected at the promoter and a critical enhancer of *BCL11A* in HUDEP-2 cells, suggesting that *BCL11A* is a direct transcriptional target of MLL1 complex. Consistent with these results, *MEN1* or *KMT2A* knockdown in normal human CD34^+^ hematopoietic stem and progenitor cells (HSPCs) induced to undergo erythroid differentiation also significantly decreased their *BCL11A* expression and increased their γ- and ε-globin expression and the production of F cells in the culture. Treatment of these cells with MENIN inhibitors yielded similar results and promoted erythroid differentiation with minimal effects on their growth. These findings underscore a critical role of MLL1 complex in regulating fetal and embryonic hemoglobin expression and suggest that MENIN inhibitors could offer a promising therapeutic approach for sickle cell disease and β-thalassemia.

## Introduction

β-globinopathies, including sickle cell disease (SCD) and β-thalassemia, are caused by β-globin gene mutations and are among the most prevalent monogenic disorders globally. SCD has been successfully treated with non-myeloablative hematopoietic stem cell (HSC) transplantation^1,2^. Gene therapy and gene editing approaches are also potentially curative for β-globinopathies ^3^. However, due to high cost and procedural complexity for these treatments, pharmacological approaches, particularly re-activating fetal-type hemoglobin (HbF), still offer an attractive and promising treatment option for β-globinopathies^4^. HbF’s short-term presence after birth is responsible for the absence of SCD disease symptoms early in life, and hydroxyurea, an FDA-approved drug for treating SCD, reduces symptoms later in life through induction of HbF expression^5–8^. To better understand the molecular mechanisms controlling the repression of HbF in adult cells, a number of genes encoding transcription factors, chromatin-modifying enzymes, and translational regulators have been identified to play significant roles in this process^9–25^. Targeting these genes in erythroid cells *in vitro* or in animal models leads to increased embryonic and fetal hemoglobin expression. Among these regulators, *BCL11A* has been considered as a major controller of HbF repression based on genome-wide association studies in human^26^ and also studies identifying its expression as a crucial target of several known regulators of HbF repression^11,13,23,24^. Developing approaches to downregulate *BCL11A* expression in erythroid cells holds great promise for treating SCD and β-thalassemia. It is possible that the transcription of *BCL11A* may be critically regulated by epigenetic mechanisms and could therefore be targeted by epigenetic modulation; however, such mechanisms remain unknown.

The MLL1 complex is a multiprotein histone methyltransferase complex capable of catalyzing mono-, di-, and tri-methylation of lysine 4 on histone 3 (H3K4) and thought to play an important role in activating transcription. The central components of the complex, KMT2A-N and KMT2A-C (also known as MLL1-N and MLL1-C), are encoded by the *KMT2A* (also known as *MLL1*, *MLL*, *HRX*, *ALL1*) gene, homologous to *Drosophila melanogaster Trithorax*. The full length KMT2A protein is proteolytically processed by Taspase-1 into KMT2A-N and KMT2A-C that normally associate with one another in the MLL1 complex ^27,28^. Additional key components of this complex include MEN1 (also known as MENIN) ^29^, PSIP1 (also known as LEDGF) ^30^, ASH2L^31^, WDR5^32,33^, RBBP5^34,35^ and DPY30^36,37^. Both MEN1 and PSIP1 are essential for the MLL1 complex to bind to chromatin^30^. Gene targeting studies of *Kmt2a* and *Men1* in mice have indicated an important role for the MLL1 complex in normal fetal and adult hematopoiesis^38,39^. *KMT2A* also is frequently involved in chromosomal translocations in acute myeloid leukemia (AML) and acute lymphoblastic leukemia (ALL), leading to the production of fusion proteins with the N-terminal part of KMT2A fused with C-terminus of over 90 different fusion partners^40^. The oncogenic activity of KMT2A fusion proteins also require their interaction with MEN1, which prompted the development of MENIN inhibitors capable of blocking such interaction for the treatment of leukemia. These inhibitors have demonstrated significant anti-leukemia activity with only mild side effects in clinical testing with Revumenib recently approved by FDA.

In this study, we identify the MLL1 complex as an important regulator of fetal hemoglobin repression through its direct regulation of *BCL11A* transcription. Our studies also identify MENIN inhibitors as promising drug candidates for treating SCD and β-thalassemia.

## Results

### *MEN1* and *KMT2A* are critical regulators of fetal and embryonic hemoglobin silencing in HUDEP-2 cells

We have previously identified *Pogz* as a critical regulator of fetal hemoglobin repression^13^. Pogz knockout fetal liver cells express significantly higher levels of *Kmt2a* than wild-type fetal liver cells^13^, suggesting that *Kmt2a* may also play a role in regulating fetal hemoglobin repression. To test this possibility, we transduced human erythroblast HUDEP-2 cells^41^ with 2 independent lentiviral shRNAs targeting *KMT2A* and a non-targeting control shRNA and examined the effects of *KMT2A* downregulation on the expression of γ- and ε-globin. Interestingly, *KMT2A* knockdown induced an up to over 15-fold increase in both γ- and ε-globin mRNA levels without significantly affecting *β-globin* expression in these cells (Figure 1A and 1B). As KMT2A is known to interact and function together with MEN1 in the MLL1 complex, we next decreased *MEN1* expression using lentiviral shRNAs in HUDEP-2 cells. Similar increases in γ- and ε-globin mRNA levels were also observed upon *MEN1* knockdown (Figure 1C and 1D), indicating that MLL1 complex is likely involved in the regulation of fetal hemoglobin repression.

**Figure 1.**
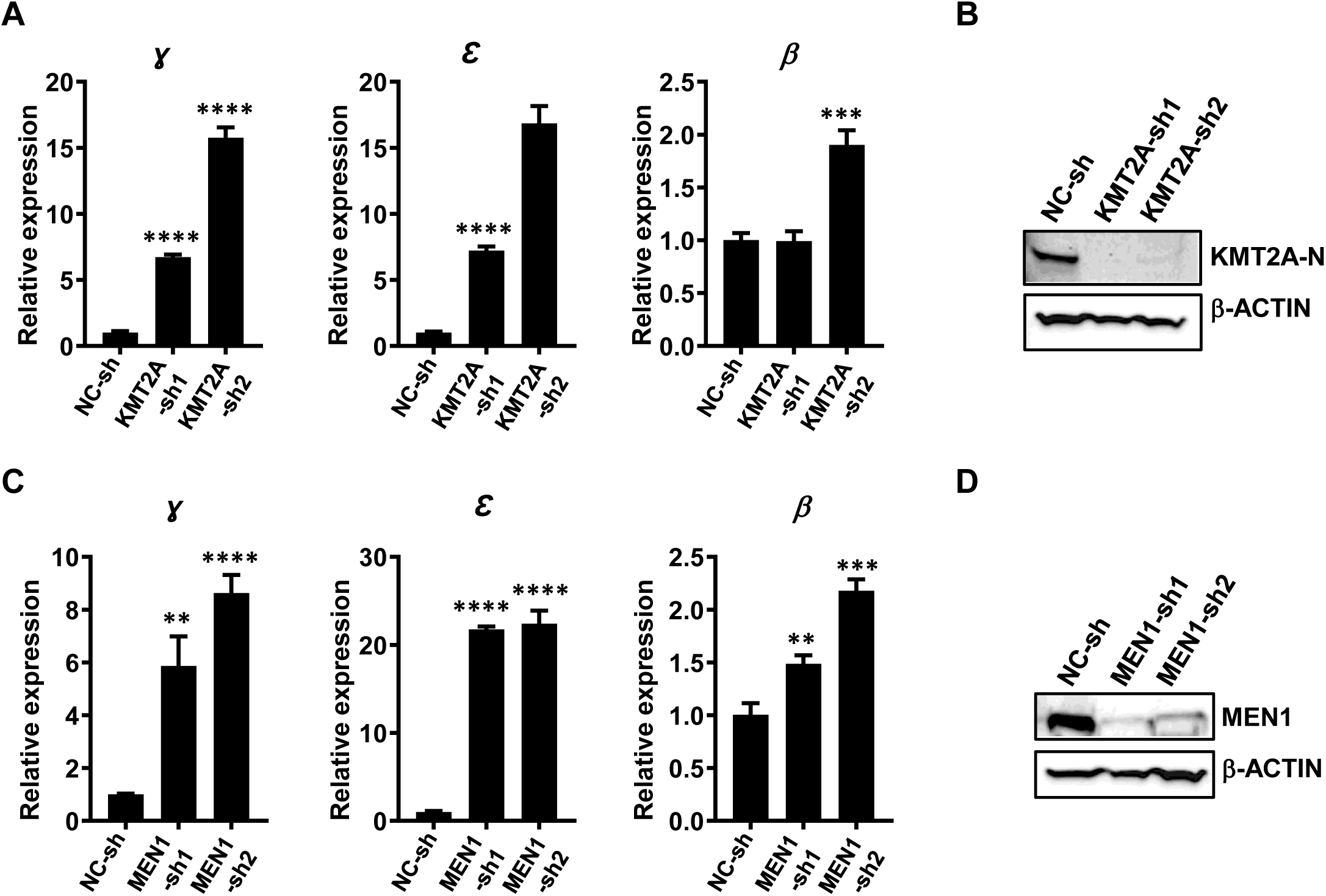
*KMT2A* and *MEN1* knockdown increased fetal and embryonic hemoglobin expression in HUDEP-2 cells. (A) Real-time RT-PCR analysis of γ−, ε−, and β-globin mRNA levels in HUDEP-2 cells at 96 hours after infection with lentiviral shRNAs targeting KMT2A (KMT2A-sh1 and -sh2) or non-targeting control lentiviral shRNA (NC-sh). Relative expression levels were calculated by normalizing to *RPL4* mRNA levels in the same sample and also to cells infected with NC-sh virus. Data are represented as mean +/- SD (n=3). (B) Representative Western blotting analysis of protein levels of KMT2A and control β-ACTIN in HUDEP-2 cells at 72 hours after infection with the indicated lentiviral shRNAs. (C) Real-time RT-PCR analysis of γ−, ε−, and β-globin mRNA levels in HUDEP-2 cells at 96 hours after infection with lentiviral shRNAs targeting MEN1 (MEN1-sh1 and -sh2) or non-targeting control lentiviral shRNA (NC-sh). Relative expression levels were calculated by normalizing to *RPL4* mRNA levels in the same sample and also to cells infected with NC-sh virus. Data are represented as mean +/- SD (n=3). (D) Representative Western blotting analysis of protein levels of MEN1 and control β-ACTIN in HUDEP-2 cells at 72 hours after infection with the indicated lentiviral shRNAs. **, *p* < 0.01; ***, *p* < 0.001; ****, *p* < 0.0001 (two-tailed Student’s *t* test).

### *MEN1* and *KMT2A* are critical regulators of *BCL11A* expression

To investigate the mechanisms underlying the regulation of fetal hemoglobin expression by KMT2A and MEN1, we carried out RNA-seq studies to identify gene expression changes at 72 hours after *MEN1* knockdown by lentiviral shRNA in HUDEP-2 cells. We found that 3063 genes were differentially expressed between the *MEN1* knockdown and control populations with 1417 genes downregulated in the knockdown cells and 1646 genes upregulated using 2-fold change in gene expression as cutoff (Figure 2A and Table S1). To identify pathways regulated by *MEN1*, we performed gene set enrichment analysis (GSEA) of the differentially expressed genes after *MEN1* knockdown. Analysis using ontology gene sets showed that *MEN1* knockdown significantly downregulated genes involved in a number of processes including ribonucleoprotein biogenesis and tRNA metabolism (Supplementary Figure 1). Importantly, consistent with the increased in γ- and ε-globin mRNA levels upon *MEN1* knockdown, *BCL11A* known as a major regulator of fetal hemoglobin repression was significantly downregulated in *MEN1* knockdown cells (Figure 2B). This result was confirmed by real-time RT-PCR in HUDEP-2 cells transduced with both shRNAs targeting MEN1 (Figure 2C, left panel). BCL11A protein levels also decreased significantly after *MEN1* knockdown (Figure 2C, right panel). Similar results were also observed after *KMT2A* knockdown in HUDEP-2 cells (Figure 2D), suggesting that *BCL11A* expression in erythroblast cells is in part controlled by the MLL1 complex and the significant decrease in BCL11A levels is likely responsible for the increases in γ- and ε-globin mRNA levels. Moreover, a significant reduction of *BCL11A* mRNA levels could be detected in HUDEP-2 cells as early as 36 hours after transduction with lentiviral shRNAs targeting *MEN1* (Supplementary Figure 2), further suggesting that *BCL11A* could be a direct transcriptional target of MLL1 complex. To test this notion, we performed chromatin immunoprecipitation (ChIP) analysis in HUDEP-2 cells at 72 hours after transduction by the non-targeting control or a *MEN1*-specific lentiviral shRNA using an antibody specific for MEN1 and control IgG. Significant bindings of MEN1 were readily detected at promoter region and a critical enhancer of *BCL11A*, but not at a control region located at 16.8 kb upstream of the transcriptional start site (Figure 2E). As expected, a significant reduction in MEN1-binding was detected at the 2 positive MEN1-binding regions in *MEN1* knockdown cells. Similar results were also observed when a KMT2A-speific antibody was used for ChIP analysis in control and *KMT2A* knockdown HUDEP-2 cells (Figure 2F). These results suggest that *BCL11A* is a direct transcriptional target of the MLL1 complex.

**Figure 2.**
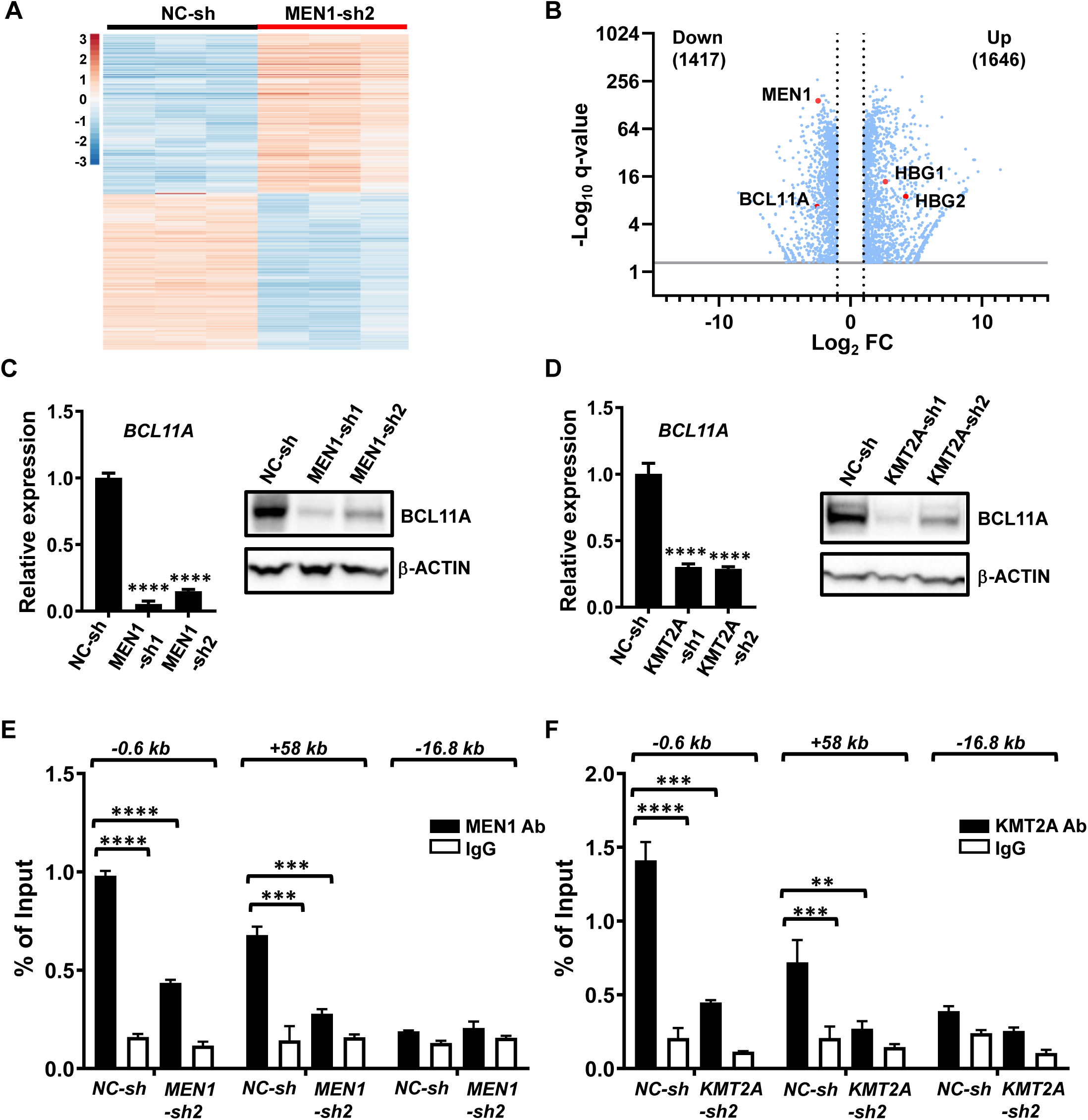
*BCL11A* is a direct transcriptional target of MLL1 complex. (A) Heatmap showing differential gene expression at 72 hours after *MEN1* knockdown in HUDEP-2 cells (B) Volcano plot showing differential gene expression upon *MEN1* knockdown in HUDEP-2 cells. (C) Left panel, real-time RT-PCR analysis of *BCL11A* mRNA levels in HUDEP-2 cells at 72 hours after infection with lentiviral shRNAs targeting *MEN1* (MEN1-sh1 and -sh2) or non-targeting control lentiviral shRNA (NC-sh). Relative expression levels were calculated by normalizing to *RPL4* mRNA levels in the same sample and also to cells infected with NC-sh virus. Data are represented as mean +/- SD (n=3). Right panel, representative Western blotting analysis of protein levels of BCL11A and control β-ACTIN in HUDEP-2 cells at 72 hours after infection with the indicated lentiviral shRNAs. (D) Left panel, real-time RT-PCR analysis of *BCL11A* mRNA levels in HUDEP-2 cells at 72 hours after infection with lentiviral shRNAs targeting *KMT2A* (KMT2A-sh1 and -sh2) or non-targeting control lentiviral shRNA (NC-sh). Data are represented as mean +/- SD (n=3). Right panel, representative Western blotting analysis of protein levels of BCL11A and control β-ACTIN in HUDEP-2 cells at 72 hours after infection with the indicated lentiviral shRNAs. (E) Quantitative ChIP analysis of indicated regions of *BCL11A* promoter (-0.6 kb), enhancer (+58 kb), and control (-16.8 kb) region in HUDEP-2 cells at 72 hours after transduction with lentiviral shRNA NC-sh or MEN1-sh2 using a MEN1-specific antibody. Data are represented as mean +/- SD (n=3). (F) Similar ChIP analysis as in (E) using HUDEP-2 cells at 72 hours after transduction with lentiviral shRNA NC-sh or KMT2A-sh2 and a KMT2A-N-specific antibody. **, *p* < 0.01; ***, *p* < 0.001; ****, *p* < 0.0001 (two-tailed Student’s *t* test).

### *MEN1* and *KMT2A* are critical regulators of fetal and embryonic hemoglobin silencing in normal human erythroid cells

To start assessing the role of MEN1 and KMT2A in regulating fetal and embryonic hemoglobin expression in normal human erythroid cells, we examined their expression during erythroid differentiation of adult human CD34^+^ hematopoietic stem and progenitor cells (HSPCs) in a three-phase *in vitro* erythroid culture system as previously described^42,43^. Consistent with the regulation of *BCL11A* transcription by the MLL1 complex, we found that MEN1 and KMT2A proteins are expressed at significant levels during early phase of differentiation and that their expression declines gradually during terminal differentiation (Supplementary Figure 3). To investigate whether *MEN1* regulates fetal and embryonic hemoglobin silencing in normal human cells, we induced erythroid differentiation of CD34^+^ HSPCs after transducing them with *MEN1*-specific and control lentiviral shRNAs at day 2 of the expansion phase and assessed fetal and embryonic hemoglobin expression at 11 days after differentiation induction. As a positive control, we also knocked down *BCL11A* using a previously published lentiviral shRNA^10^. Real-time RT-PCR analysis showed that both *MEN1* shRNAs significantly increased γ and ε-globin transcript levels compared to control shRNA without causing overt changes to mRNA levels of β-globin (Figure 3A). Overall, *MEN1* knockdowns increased the proportion of γ-globin transcripts in total β-like globin transcripts to over 35%, similar to the increase induced by *BCL11A* knockdown (Figure 3B). Consistent with the increases in the mRNA levels, γ-globin protein levels also significantly increased in *MEN1* and *BCL11A* knockdown cells over control cells at day 14 of differentiation (Figure 3C). After quantification by HPLC, the γ-globin protein levels reached over 22% of total β-like globin levels in cells transduced by MEN1-sh2 (Figure 3D). There was also a significant increase in the percentage of F cells produced in the knockdown cultures (Figure 3E). In supporting the role of MLL1 complex in activating *BCL11A*, both mRNA and protein levels of BCL11A were significantly reduced in the *MEN1* knockdown cells (Figure 3F). We also followed the effects of *MEN1* knockdown on the progression of erythroid differentiation by monitoring the expression of CD71 and CD235a of the knockdown cells in comparison to control cells over the differentiation process. No significant effects by the knockdown on erythroid differentiation were observed (Figure 3G). Terminal differentiation was not blocked as many orthochromatic erythroblasts and reticulocytes can be observed in the day 14 cultures of *MEN1* knockdown cells (Supplementary Figure 4). A significant reduction in the number of cells generated in the knockdown cultures compared to the control culture also was observed (Figure 3H). Since there was no consistent increase in apoptosis detected in the knockdown cultures over the control culture (Supplementary Figure 5), this suggests that *MEN1* may be important for the proliferation of differentiating erythroid progenitors. Taken together, these results suggest that *MEN1* is critical for the repression of fetal and embryonic hemoglobin and the growth of the erythroid progenitors but may not be essential for erythroid differentiation.

**Figure 3.**
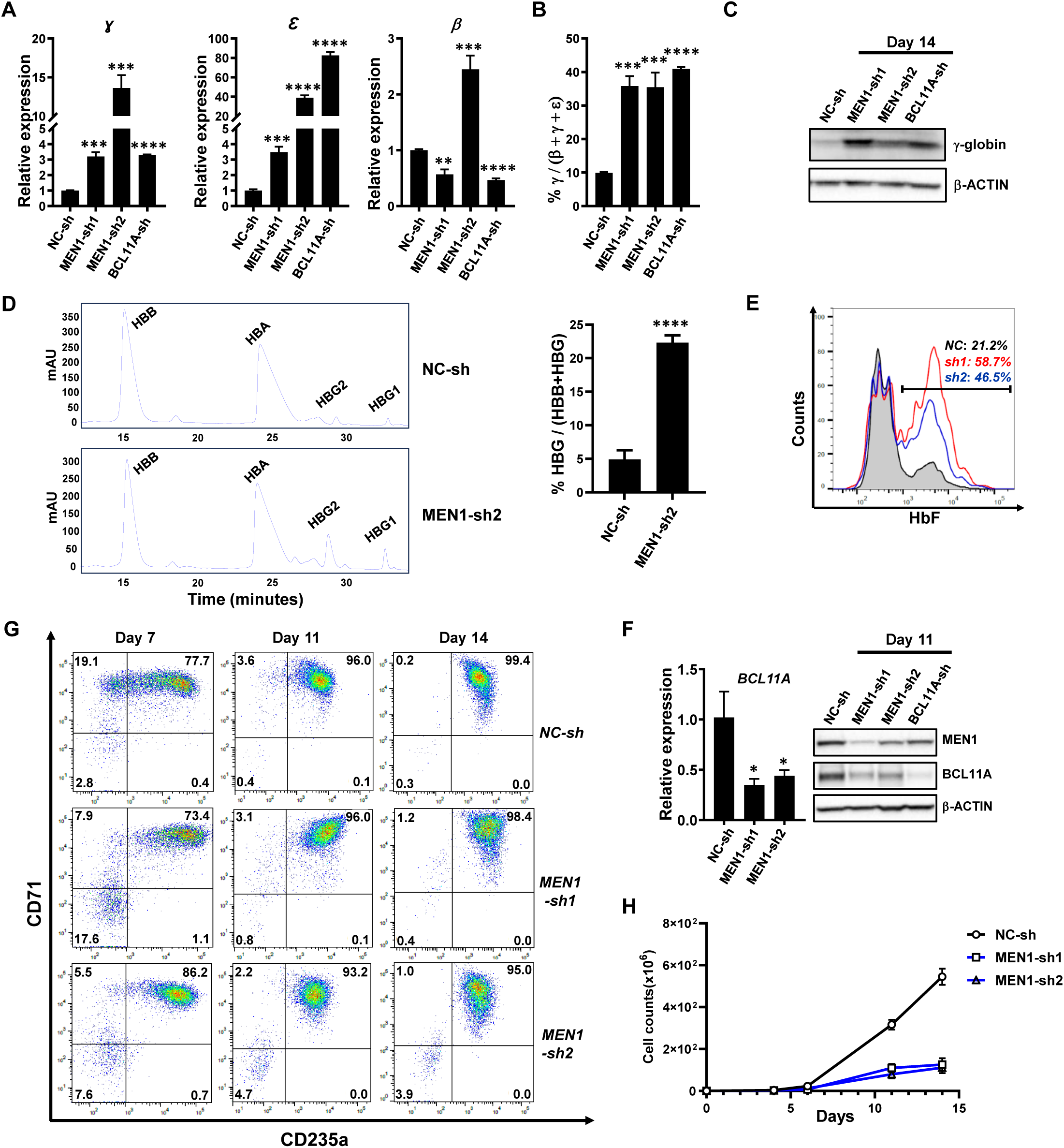
*MEN1* knockdown significantly increased fetal and embryonic hemoglobin expression in erythroid cells differentiated from normal human CD34^+^ HSPCs. Human CD34^+^ HSPCs transduced by lentiviral shRNAs targeting *MEN1* (MEN1-sh1 and -sh2), *BCL11A* (BCL11A-sh), or non-targeting control lentiviral shRNA (NC-sh) were induced to undergo erythroid differentiation. (A) γ-, ε-, and β-globin mRNA levels in cells at day 11 after differentiation induction were analyzed by real-time RT-PCR. Relative expression levels were calculated by normalizing to *GAPDH* mRNA levels in the same sample and also to cells infected with NC-sh virus. Data are represented as Mean +/- SD (n=3). (B) Bar graph showing γ-globin mRNA levels as a percentage of total β-like globin (β+g+e) mRNA levels in transduced cells based on results in A. (C) Representative Western blotting analysis of protein levels of γ-globin and control β-ACTIN in cells transduced by indicated lentiviral shRNAs at day 14 after induction of differentiation. (D) Left panels, representative chromatograms from HPLC analysis of hemoglobin levels in cells transduced by NC-sh or MEN1-sh2 at day 14 of differentiation. Right panel, Mean and SD of percentages of γ-globin (HBG1+HBG2) levels of total β-like globin (HBB+HBG1+HBG2) levels assessed by HPLC analysis (n=3). (E) Representative flow cytometry analysis of F cells in indicated control and MEN1 knockdown cultures at day 14 of differentiation. (F) Left panel, *BCL11A* mRNA levels in cells at day 11 after differentiation induction analyzed by real-time RT-PCR. Data are represented as Mean +/- SD (n=3). Right panel, Representative Western blotting analysis of protein levels of MEN1, BCL11A and control GAPDH in the same cells. (G) Representative flow cytometry analysis of CD71 and CD235a expression on cells generated from indicated transduced CD34^+^ HSPCs at day 7, 11, and 14 of erythroid differentiation. (H) Growth curves of differentiation cultures of indicated transduced CD34^+^ HSPCs (n=3). *, *p* < 0.05; **, *p* < 0.01; ***, *p* < 0.001; ****, *p* < 0.0001 (two-tailed Student’s *t* test).

Similarly, we also examined the role of *KMT2A* in fetal and embryonic hemoglobin silencing by knocking down its expression in CD34^+^ HSPCs. Analogous to the effects of *MEN1* knockdown, the knockdown of *KMT2A* induced significant increases in γ and ε-globin mRNA levels, γ-globin protein expression, and the percentage of F cells (Figure 4A-4E). Additionally, *BCL11A* expression was downregulated (Figure 4F). We also observed a delay in maturation at day 7 of differentiation but no block in terminal differentiation (Figure 4G and Supplementary Figure 4). As with *MEN1* knockdown, erythroid cell growth was reduced following *KMT2A* knockdown (Figure 4H and Supplementary Figure 5). These findings support that MLL1 complex is an important regulator of fetal and embryonic hemoglobin silencing and growth of erythroid precursor cells.

**Figure 4.**
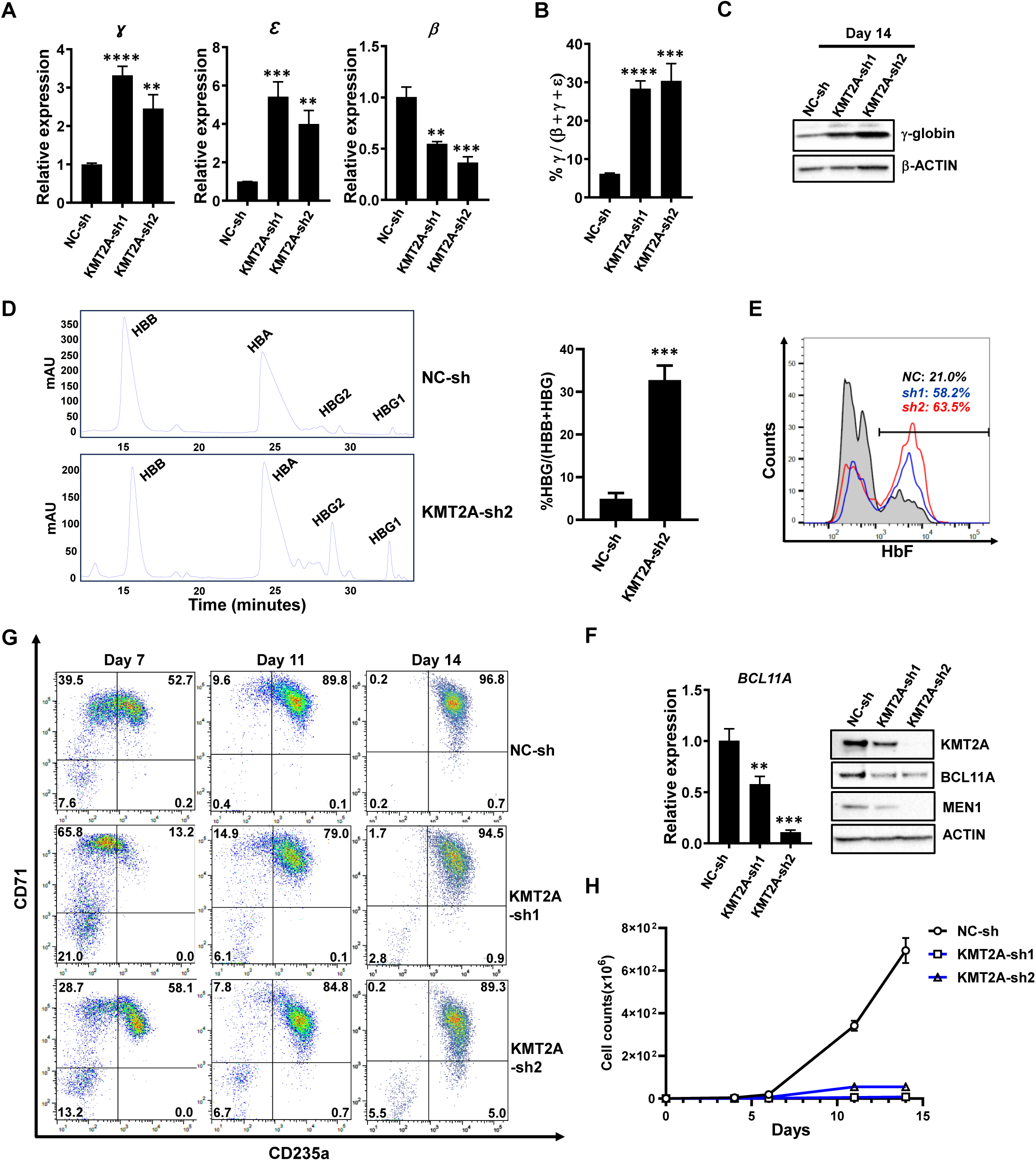
*KMT2A* knockdown significantly increased fetal and embryonic hemoglobin expression in erythroid cells differentiated from normal human CD34^+^ HSPCs. Human CD34^+^ HSPCs transduced by lentiviral shRNAs targeting *KMT2A* (KMT2A-sh1 and -sh2) or non-targeting control lentiviral shRNA (NC-sh) were induced to undergo erythroid differentiation. (A) γ-, ε-, and β-globin mRNA levels in cells at day 11 after differentiation induction were analyzed by real-time RT-PCR. Relative expression levels were calculated by normalizing to GAPDH mRNA levels in the same sample and also to cells infected with NC-sh virus. Data are represented as mean +/- SD (n=3). (B) Bar graph showing γ-globin mRNA levels as a percentage of total β-like globin (β+g+e) mRNA levels in transduced cells based on results in A. (C) Representative Western blotting analysis of protein levels of γ-globin and control β-ACTIN in cells transduced by indicated lentiviral shRNAs at day 14 after induction of differentiation. (D) Left panels, representative chromatograms from HPLC analysis of hemoglobin levels in cells transduced by NC-sh or KMT2A-sh2 at day 14 of differentiation. Right panel, Mean and SD of percentages of γ-globin (HBG1+HBG2) levels of total β-like globin (HBB+HBG1+HBG2) levels assessed by HPLC analysis (n=3). (E) Representative flow cytometry analysis of F cells in indicated control and *KMT2A* knockdown cultures at day 14 of differentiation. (F) Left panel, *BCL11A* mRNA levels in cells at day 11 after differentiation induction analyzed by real-time RT-PCR. Data are represented as mean +/- SD (n=3). Right panel, representative Western blotting analysis of protein levels of KMT2A, BCL11A, MEN1 and control β-ACTIN in the same cells. (G) Representative flow cytometry analysis of CD71 and CD235a expression on cells generated from indicated transduced CD34^+^ HSPCs at day 7, 11, and 14 of erythroid differentiation. (H) Growth curves of differentiation cultures of indicated transduced CD34^+^ HSPCs (n=3). **, *p* < 0.01; ***, *p* < 0.001; ****, *p* < 0.0001 (two-tailed Student’s *t* test).

MENIN inhibitors have been designed to disrupt the function of MLL1 complex by blocking the interaction between MEN1 and KMT2A for the treatment of AML and mild side effects have been reported for these inhibitors in clinical trials ^44–46^. To start testing the possibility that they also could be used for the treatment of SCD and β-thalassemia, we examined their capability to induce fetal hemoglobin expression in CD34^+^ HSPCs. Starting from day 2 of the expansion phase, treatment with MENIN inhibitor MI-3454 at 2 different concentrations (0.2 and 0.5 µM) induced a dose dependent upregulation of γ and ε-globin mRNA levels with the highest proportion of γ-globin transcripts in total β-like globin transcripts reaching close to 30% at day 11 of differentiation (Figure 5A and 5B). MI-3454 treatment also induced a significant increase in γ-globin protein levels detected by Western blotting analysis (Figure 5C). HPLC quantification further show an increase of up to 3-fold in the proportion of γ-globin protein levels in total β-like globin levels induced by MI-3454 (Figure 5D). F-cells numbers were also significantly increased by MI-3454 (Figure 5E). As expected, these increases were accompanied by a downregulation of *BCL11A* expression levels (Figure 5F). Consistent with studies in leukemia cells, treatment with MI-3454 also caused a downregulation of MEN1 protein levels (Figure 5F). Treatment with MI-3454 also promoted differentiation of the erythroid progenitor cells as significantly more cells became positive for both CD71 and CD235a expression at day 7 of differentiation and more cells displaying reduced levels of CD71 at day 14 of differentiation compared to control cells (Figure 5G). In addition, only a slight reduction in cell growth was observed at late stage of differentiation for cells treated with higher MI-3454 dosage (Figure 5H), suggesting minimal toxicity of the treatment to the cells. Similar increases in the proportion of γ-globin protein levels also were detected by Western blot and HPLC with two additional MENIN inhibitors Ziftomenib (KO-539) and Revumenib (SNDX-5613), which have been tested in clinical trials for the treatment of AML (Figure 5I). These results support that MENIN inhibitors are effective in increasing HbF expression in human adult erythroid cells.

**Figure 5.**
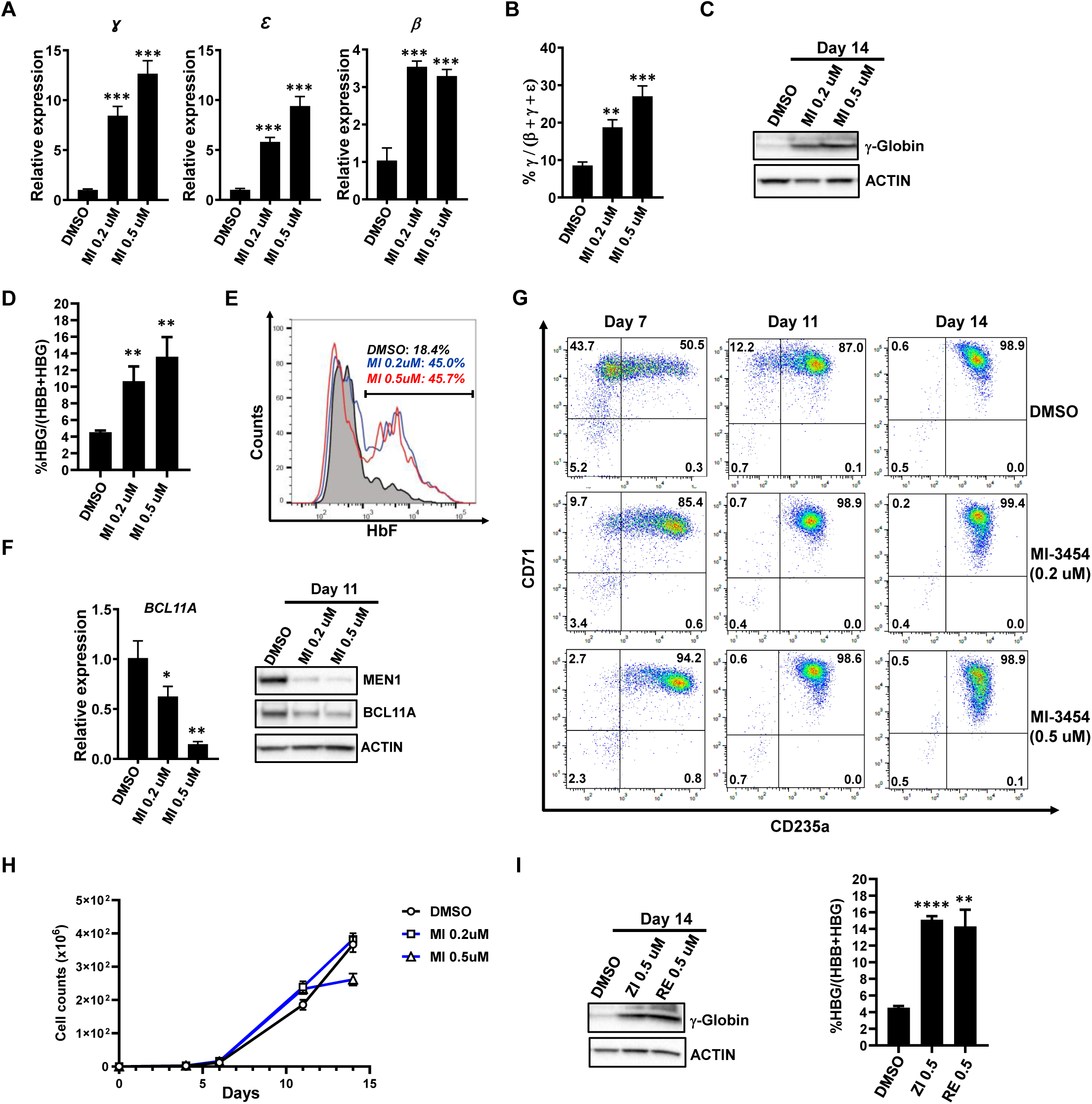
MENIN inhibitors significantly increased fetal and embryonic hemoglobin expression in erythroid cells differentiated from normal human CD34^+^ HSPCs. Human CD34^+^ HSPCs were treated with MI-3454 during induced erythroid differentiation. (A) γ-, ε-, and β-globin mRNA levels in cells at day 11 after differentiation induction were analyzed by rea-time RT-PCR. Relative expression levels were calculated by normalizing to *GAPDH* mRNA levels in the same sample and also to cells treated with DMSO. Data are represented as mean +/- SD (n=3). (B) Bar graph showing γ-globin mRNA levels as a percentage of total β-like globin (β+g+e) mRNA levels in MI-3454-treated and control cells based on results in A. (C) Representative Western blotting analysis of protein levels of γ- globin and control β-ACTIN in MI-3454-treated and control cells at day 14 after induction of differentiation. (D) Mean and SD of percentages of γ-globin (HBG1+HBG2) levels of total β-like globin (HBB+HBG1+HBG2) levels assessed by HPLC analysis (n=3). (E) Representative flow cytometry analysis of F cells in MI-3454-treated and control cultures at day 14 of differentiation. (F) Left panel, *BCL11A* mRNA levels in cells at day 11 after differentiation induction analyzed by real-time RT-PCR. Data are represented as mean +/- SD (n=3). Right panel, representative Western blotting analysis of protein levels of MEN1, BCL11A and control GAPDH in MI-3454-treated and control cells at day 11 after induction of differentiation. (G) Representative flow cytometry analysis of CD71 and CD235a expression on MI-3454-treated and control cells at day 7, 11, and 14 of erythroid differentiation. (H) Growth curves of differentiation cultures of MI-3454-treated and control CD34^+^ HSPCs (n=3). (I) Left panel, representative Western blotting analysis of protein levels of γ-globin and control β-ACTIN in progenies of CD34^+^ HSPCs treated with Ziftomenib (ZI), Revumenib (RE), or control DMSO at day 14 after induction of differentiation. Right panel, Mean and SD of percentages of γ-globin (HBG1+HBG2) levels of total β-like globin (HBB+HBG1+HBG2) levels in same cells assessed by HPLC analysis (n=3). *, *p* < 0.05; **, *p* < 0.01; ***, *p* < 0.001; ****, *p* < 0.0001 (two-tailed Student’s *t* test).

## Discussion

*BCL11A* is a major regulator of fetal hemoglobin repression and represents an attractive target for treating SCD and β-thalassemia; however, the mechanisms responsible for its transcriptional activation in erythroid cells remains unclear. Although several transcription factors including KLF1, POGZ, GATA1, and ATF4 have been identified to activate its transcription^11,13,24,47^, the epigenetic mechanism involved remains unknown. Our finding of the involvement of MLL1 complex in regulating *BCL11A* transcription suggests its possible recruitment by some of these transcription factors through physical interaction. This finding also raises the interesting possibility that MENIN inhibitors could be effective in increasing HbF levels in adult erythroid cells by reducing *BCL11A* expression. Our results further show that treatment with MENIN inhibitors at the tested concentrations was capable of increasing the proportion of γ-globin levels by up to 3-fold in erythroid cells generated from normal CD34^+^ HSPCs. These inhibitors, already in human clinical trials and one already FDA-approved, could be repurposed for testing in the hemoglobinopathies either alone or in combination with hydroxyurea for inducing HbF. Although MEN1 has been identified as a tumor suppressor gene in the endocrine system^48,49^, binding of MENIN inhibitors may only partially inhibit the function of the protein as these small molecules were designed to specifically bind to the MEN1 domain mediating MEN1/KMT2A interaction and MEN1 is known to interact with a number of proteins through other domains^50^. This notion is also supported by our results showing that the effects of MI-3454 treatment on erythroid cells appear to be different from that of MEN1 or KMT2A knockdown. Instead of causing no effect or a delay in early maturation while significantly reducing cell numbers, MI-3454 promoted erythroid differentiation and had minimal negative effects on cell growth. Therefore, it is possible that long-term treatment with MENIN inhibitors may not induce development of endocrine tumors. Long-term studies in animal models will be required to clarify this issue. In addition, the differentiation-promoting effect of MENIN inhibitors could also be beneficial for β-thalassemia patients as ineffective erythropoiesis has been documented in this disease^51^. Taken together, our study indicates that MENIN inhibitors could be effective for treating SCD and β-thalassemia by significantly increasing fetal hemoglobin expression.

## Acknowledgments

This work was supported by Defense Health Agency (DHA) Grant PED-86-11875 (Y.D.) and Uniformed Services University of the Health Sciences (USUHS) Grant RAMP310534 (Y.D.). This study also was supported by the Division of Intramural Research, NHLBI and federal funds from the Frederick National Laboratory for Cancer Research, NIH (HHSN261200800001E).

The views presented in this manuscript are those of the authors; no endorsement by USUHS or the Department of Defense has been given or should be inferred. The content of this publication also does not necessarily reflect the views or policies of the Department of Health and Human Services, nor does the mention of trade names, commercial products, or organizations imply endorsements by the US Government.

## Author Contributions

Y.H. performed most of the experiments and data analysis; B.G. performed experiments and data analysis and contributed to writing of the manuscript; K.O.G. performed data analysis and contributed to writing of the manuscript; K.R.R. performed experiments; J.T. contributed vital reagents and to writing of the manuscript; Y.D. designed and performed research, analyzed data, and wrote the manuscript.

## Declaration of Interests

Y.D. is listed as inventor on a PCT application No. PCT/US2022/014109, entitled: “Reactivation of Embryonic and fetal Hemoglobin,” applied for by the Henry M. Jackson Foundation of Military Medicine, Inc. The other authors declare no conflict of interests.

## Methods

### Cell culture

HUDEP-2 (RCB4557) cells were provided by the RIKEN BRC through the National Bio-Resource Project of the MEXT/AMED, Japan, and maintained in StemSpan SFEM supplemented with 3 units/ml EPO, 50ng/ml SCF, 10^-6^ M dexamethasone, and 1 µg/ml Doxycycline. Purified peripheral blood CD34+ cells were purchased from StemCell Technologies or obtained from the National Heart, Lung, and Blood Institute (NHLBI) and the National Institute of Diabetes, Digestive, and Kidney diseases (NIDDK) under studies 02-H-0160 and 08-H-0156. Cells are first expanded in StemSpan-XF media supplemented with CC100 (StemCell Technologies) for 4 days at a density of 10^5^ cells/ml. Erythroid differentiation is subsequently induced in 3 phases using a base culture medium of Iscove’s Modified Dulbecco’s Medium with addition of 1% pen/strep, 2% human peripheral blood plasma (Innovative Research), 3% human AB serum (Sigma Aldrich), 200 ug/mL Holo human transferrin (Sigma Aldrich), 3 IU/ml heparin (Sigma Aldrich), and 10 ug/mL human insulin (Sigma Aldrich). In phase I (day 0 to day 6), CD34^+^ cells after expansion are cultured at a concentration of 10^5^/ml in the presence of 10 ng/mL stem cell factor, 1 ng/mL IL-3, and 3 IU/mL erythropoietin (Epogen or Procrit). In phase II (day 7 to day 11), only stem cell factor and erythropoietin are added to the base medium, and cells are seeded at 2 x10^5^/ml concentration. In phase III (day 12 to day 21, the cell concentration was adjusted to 10^6^/mL at the beginning), the medium for this phase was the base medium plus 3IU/ml erythropoietin, and the concentration of transferrin was adjusted to 1 mg/ml. All cell cultures were incubated in 5% CO2 at 37°C.

### Drug treatment

MENIN inhibitors MI-3454, Ziftomenib (KO-539) and Revumenib (SNDX-5613) were purchased from MedChemExpress (Monmouth Junction, NJ) and dissolved in DMSO. Drug treatments for CD34^+^ HSPCs were started at day 2 of the expansion phase and were replenished every 2 days afterwards by replacing 75% of old medium with fresh medium with new drugs.

### Lentiviral production, infection, and analysis

pLKO.1 lentiviral constructs containing shRNAs for NC-sh, KMT2A-sh1, KMT2A-sh2, MEN1-sh1, and MEN1-sh2 were purchased from Sigma Aldrich (NC-sh, SHC002; KMT2A-sh1, TRCN0000005954; MEN1-sh1, TRCN0000040140; MEN1-sh2, TRCN0000040142; St. Louis, MO). KMT2A-sh2 was cloned into pLKO.1 and its targeting sequence is 5′ TGC CAA GCA CTG TCG AAA TTA 3′. Infectious lentivirus was produced and tittered as described previously ^52^. Lentiviral transduction of cells was carried out by spinoculation in which a mixture of lentivirus and target cells was spun at 2000 x g for 90 minutes at 37°C. Transduced cells were selected with puromycin (2 μg/ml) at 24 hours after infection for a duration of 48 hours.

### RNA extraction, real-time RT-PCR, and RNA-sequencing

For real-time RT-PCR, total RNA was extracted from cells using the NucleoSpin RNA Plus kit (Takara Bio). Oligo-dT-primed cDNA samples were prepared using the Superscript IV First-Strand Synthesis System (Invitrogen, Carlsbad, CA), and real-time PCR analysis was performed in triplicates using SYBR green detection reagents (Applied Biosystem) on a QuantStudio 6 Flex (Applied Biosystems). Relative changes in expression were calculated according to the ΔΔCt method. The cycling conditions were 50°C for 2 minutes followed by 95°C for 2 minutes, and then 40 cycles of 95°C for 15 seconds and 60°C for 1 minute. The gene-specific primer sequences used are listed below.

For RNA-seq, total RNA was isolated from HUDEP-2 cells at 72 hours after transduction with MEN1-sh2, or control (NC-sh) pLKO.1 lentiviral shRNA and selection with 2 ug/ml of puromycin. RNA quality was checked with the Agilent Bioanalyzer 2100 (Agilent) and sequencing libraries were generated using the NEBNext Ultra II Directional Library Prep Kit (NEB) after PolyA selection. Samples were sequenced using NovaSeq (Illumina) with ∼40 million 150bp, paired-end reads.

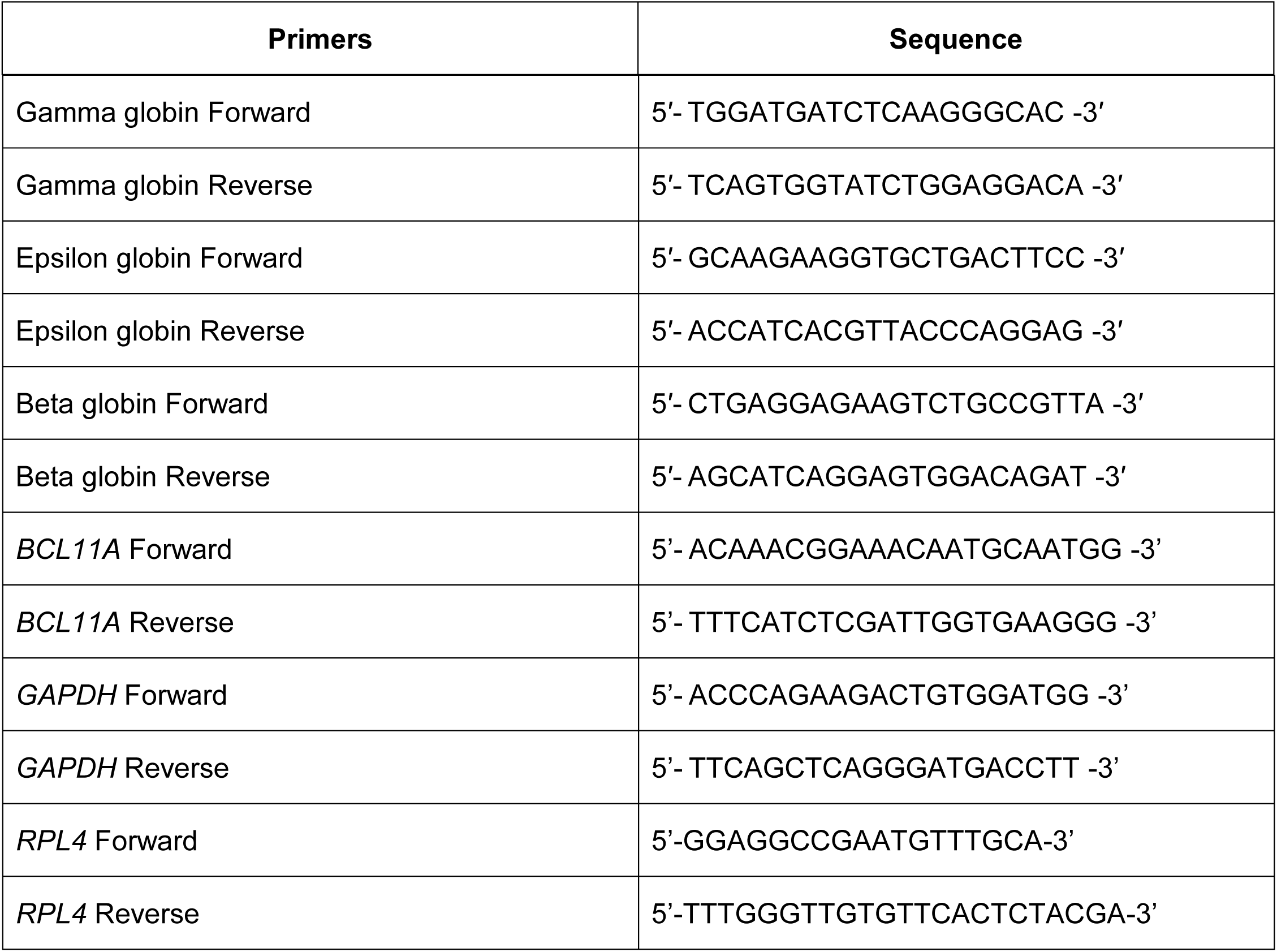

### Western blotting analysis

For western blotting analysis, the cells were washed twice with cold PBS and whole cell lysates prepared by direct lysis of cell pellets in heated 2 x SDS sample buffer. Cell lysates were resolved on tris-glycine gels (Life Technologies) before transferring onto nitrocellulose membranes (Bio-Rad). For protein detection following antibodies were used: anti-KMT2A (A300-086A, Bethyl Laboratories), anti-MEN1 (A300-115A, Bethyl Laboratories), anti-BCL11A (Ab19487, Abcam), anti-Hemoglobin γ (SC-21756, Santa Cruz Biotechnology), and anti-β-Actin (MAB1501R, Millipore). Secondary antibodies used include goat anti-rabbit (SC-2004, Santa Cruz Biotechnology). Protein bands were visualized by incubation with SuperSignal West Femto chemiluminescent substrate (Pierce).

### Chromatin Immunoprecipitation

ChIP analyses were performed using ChIP-IT Express kit (Active Motif). Immunoprecipitations were performed using anti-MLL1 (Bethyl laboratories) and control rabbit IgG (Cell Signaling). Chromatin DNA was purified using iPure kit v2 (Diagenode) and quantified by real-time PCR. The following *BCL11A*-specific primers were used:

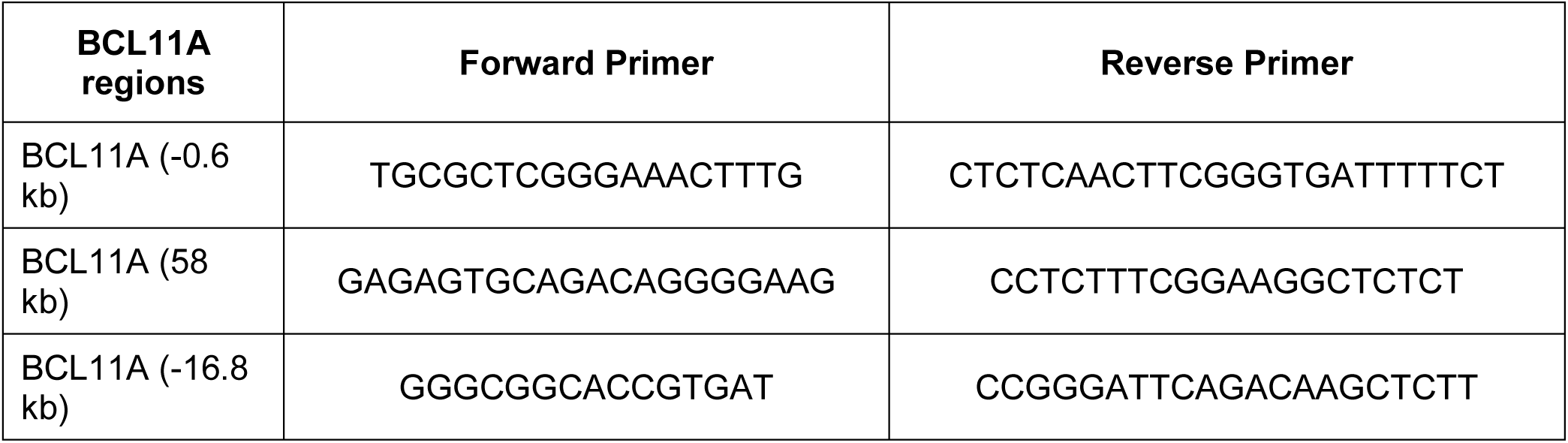

### Flow cytometry

Flow cytometry analysis of differentiating human CD34^+^ cells was performed using BD FACSymphony A5 flow cytometer. Cells were blocked by incubation with anti-FcγR-II/III and subsequently stained with antibodies against CD71 (Biolegend) and CD235a (Biolegend). Dead cells were excluded by staining with Sytox Blue (Invitrogen).

### F cells analysis

Cells were fixed with 0.05% glutaraldehyde (diluted freshly in DPBS solution) for 10 minutes at room temperature. Permeabilization off cells was performed with 0.1% Triton X-100 in DPBS at room temperature for 10 minutes. Permeabilized cells were then stained with PE-conjugated anti-HbF antibody (560041, BD Pharmingen) for 15 minutes at room temperature and analyzed using BD FACSymphony A5.

### Reversed-phase high-performance liquid chromatography (HPLC)

To determine the quantity of hemoglobin proteins (HBA, HBB, HBG1/2) in erythroblast generated in the 3-phase culture system, cell lysates were prepared and analyzed by HPLC using Agilent 1100 HPLC series (Agilent Technologies, Santa Clara, CA). Briefly, 2-3 x 10^6^ cells were washed 3 times with PBS, lysed in 50 ml HPLC grade water and centrifuged for 15 minutes at 13000 g. The supernatant was transferred to a new tube, 5 µl TCEP (100mM) added, and the solution incubated for 5 minutes at room temperature. Finally, 45 µl of 0.2% TFA /64% acetonitrile was added and the solution vortexed 3x for 30 seconds before analysis by HPLC.

### Data analysis

For RNA-seq analysis, FastQC (version v0.11.8) was applied to check the quality of raw reads. Trimmomatic (version v0.38) was applied to cut adaptors and trim low-quality bases with default setting. STAR Aligner version 2.7.1a was used to align the reads. Picard tools (version 2.20.4) was applied to mark duplicates of mapping. The StringTie version 2.0.4 was used to assemble the RNA-Seq alignments into potential transcripts. DESeq2 (v1.14.1) was used to identify differentially expressed genes (DEGs) between pairwise conditions from raw gene read counts. Significant DEGs were identified as those with an FDR q-value < 0.05 and an absolute fold change ≥ 2 (i.e. |log2 (fold-change)| ≥ 1) across samples. Gene set enrichment analysis (GSEA) was performed using the GSEA v4.3.3 application (www.gsea-msigdb.org/gsea/downloads.jsp). The RNA-seq data reported in this paper have been deposited at NCBI Gene Expression Ominibus with the accession code GSE 292144.

### Quantification and statistical analysis

Data are represented as mean +/- SD. Sample sizes were determined by previous experiences. No samples were excluded from analyses. All data except RNA-seq results were analyzed by two-tailed Student’s *t-test* using GraphPad Prism 10 software.

