## Supplementary Figures for "MLL1 complex is a critical regulator of fetal hemoglobin repression"

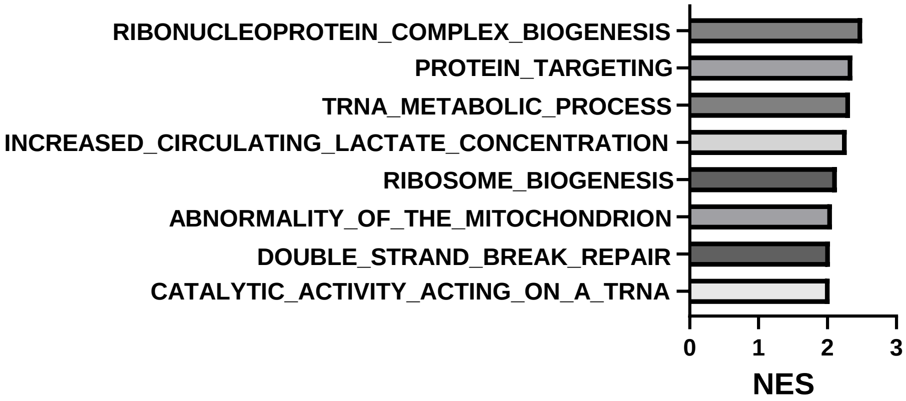


**Figure S1.** Top GO gene sets with positive enrichment from GSEA analysis of differentially expressed genes in *MEN1* knockdown HUDEP-2 cells.

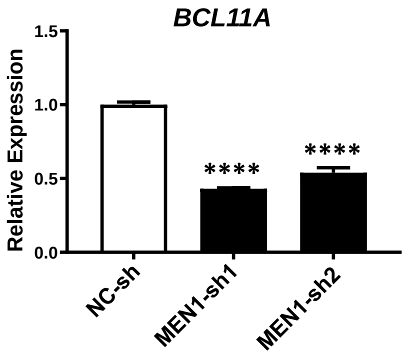


**Figure S2. *MEN1* knockdown induced a rapid reduction in *BCL11A* mRNA levels.** Real time RT-PCR analyses of *BCL11A* mRNA levels at 36 hours after infection with the indicated lentiviral shRNAs in HUDEP-2 cells. Relative expression levels were calculated by normalizing to *β-ACTIN* mRNA levels in the same sample and also in cells infected by NC-sh virus. The mean and SD of each relative expression level are shown.

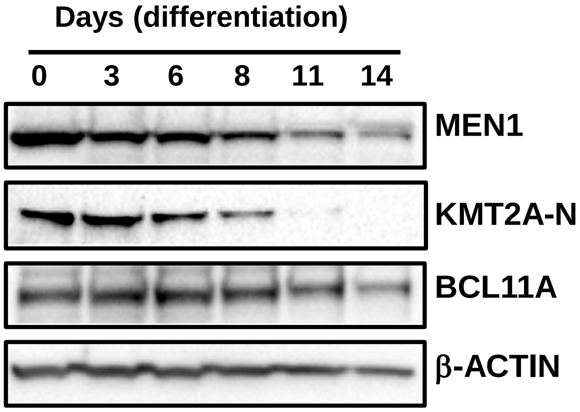


**Figure S3. Expression patterns of MEN1 and MLL1 during erythroid differentiation.** Western blotting analysis of MEN1, KMT2A-N, BCL11A, and β-ACTIN protein levels in human $CD34^+$ HSPCs at indicated time points during induced erythroid differentiation.

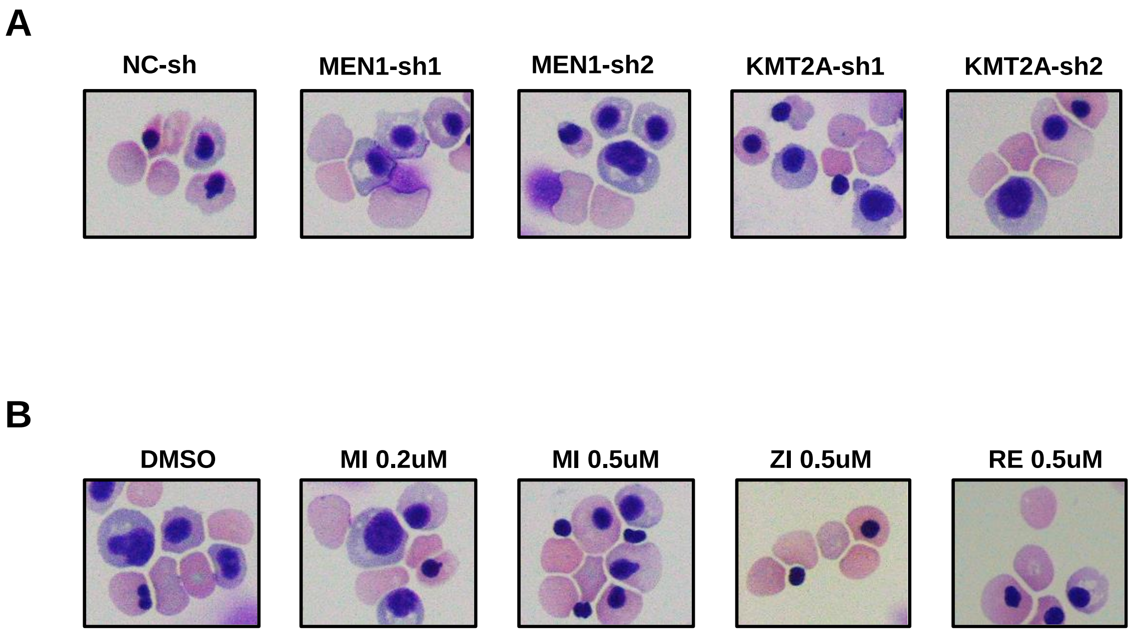


**Figure S4**. **Inhibition of MLL1 complex does not block terminal erythroid differentiation of normal human CD34$^+$ HSPCs.** A. Representative images of May-Grunwald Giemsa–stained erythroid cells generated by CD34$^+$ HSPCs after transduction by indicated lentiviral shRNAs and subsequent induction of differentiation for 14 days. B. Representative images of May-Grunwald Giemsa–stained erythroid cells generated by CD34$^+$ HSPCs after treatment with indicated MENIN inhibitors and induction of differentiation for 14 days.

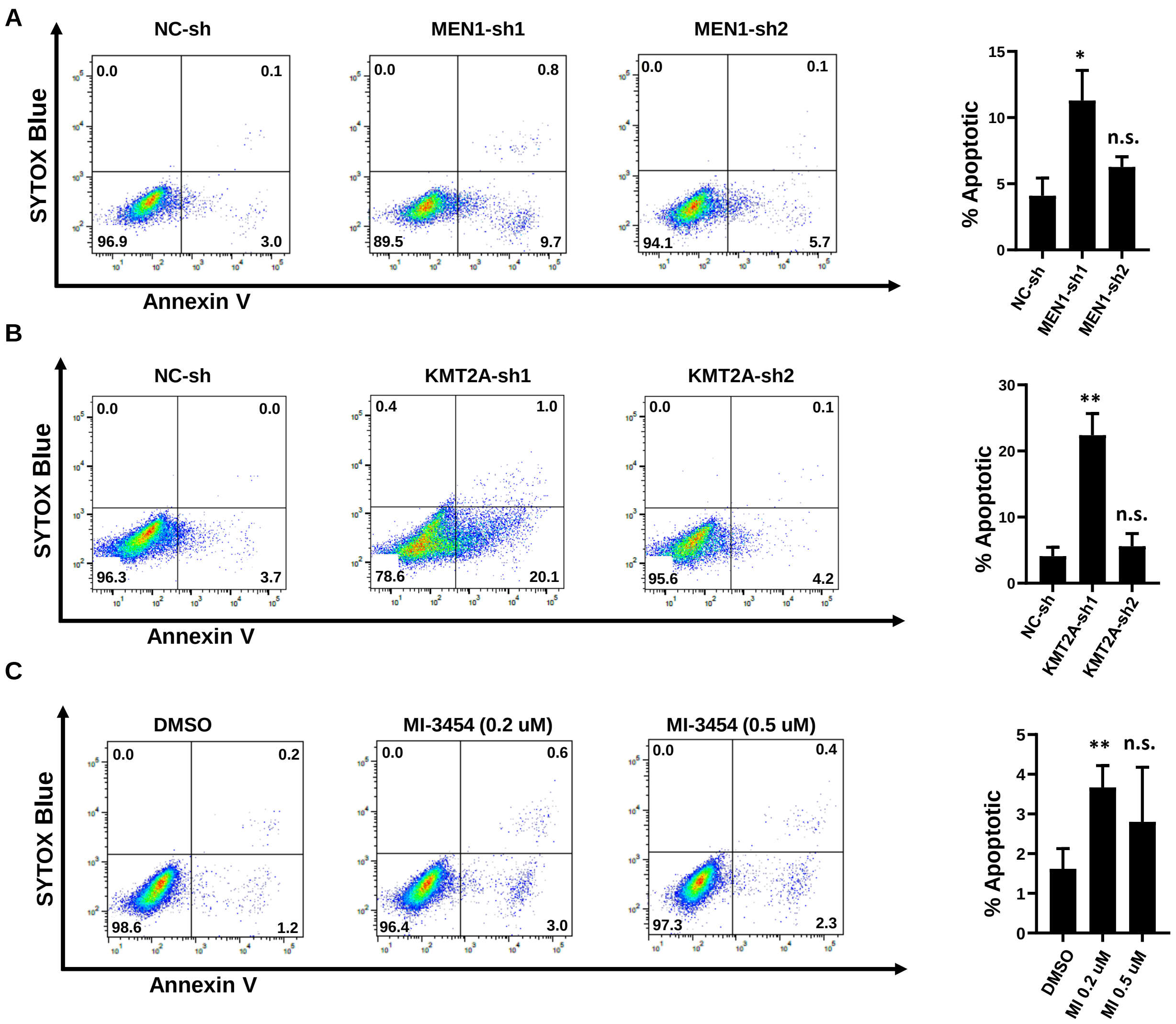


**Figure S5. Effects of MLL1 complex inhibition on apoptosis of erythroid cells generated from normal human CD34$^+$ HSPCs**. (A) Left panel, apoptotic cells detected by Annexin V staining in erythroid progenitor cells generated from CD34$^+$ HSPCs transduced by indicated *MEN1*-specific or control lentiviral shRNAs at day 4 after differentiation induction. Right panel, Mean and SD of percentages of apoptotic cells analyzed in the left panel. (B) Left panel, apoptotic cells detected by Annexin V staining in erythroid progenitor cells generated from CD34$^+$ HSPCs transduced by indicated *KMT2A*-specific or control lentiviral shRNAs at day 4 after differentiation induction. Right panel, Mean and SD of percentages of apoptotic cells analyzed in the left panel. (C) Left panel, apoptotic cells detected by Annexin V staining in erythroid progenitor cells generated from CD34$^+$ HSPCs treated with MENIN inhibitor MI-3454 at indicated concentrations at day 4 after differentiation induction. Right panel, Mean and SD of percentages of apoptotic cells analyzed in the left panel.
