## Supplementary Table 1 for "MLL1 complex is a critical regulator of fetal hemoglobin repression"

**Differentially expressed genes induced by *MEN1* knockdown in HUDEP-2 cells**

| EnsemblID | GeneSymbol | log2FoldChange | padj |
|---|---|---|---|
| ENSG00000010278 | CD9 | 11.40213162 | 3.64E-20 |
| ENSG00000140279 | DUOX2 | 9.709477882 | 3.71E-19 |
| ENSG00000066629 | EML1 | 9.453781846 | 9.75E-27 |
| ENSG00000196132 | MYT1 | 9.356172619 | 1.02E-26 |
| ENSG00000226741 | LINC02554 | 8.946016083 | 3.52E-16 |
| ENSG00000288783 | gene_sourcehavana_ | 8.853212011 | 1.66E-11 |
| ENSG00000140274 | DUOXA2 | 8.818196929 | 4.68E-11 |
| ENSG00000140522 | RLBP1 | 8.768984555 | 5.03E-12 |
| ENSG00000198910 | L1CAM | 8.750725512 | 4.10E-36 |
| ENSG00000073737 | DHRS9 | 8.655595135 | 7.76E-11 |
| ENSG00000163814 | CDCP1 | 8.615989481 | 9.77E-15 |
| ENSG00000135074 | ADAM19 | 8.584717868 | 3.69E-10 |
| ENSG00000142615 | CELA2A | 8.562687506 | 1.07E-10 |
| ENSG00000145198 | VWA5B2 | 8.554697043 | 1.21E-10 |
| ENSG00000204175 | GPRIN2 | 8.448561426 | 4.02E-10 |
| ENSG00000116701 | NCF2 | 8.353897583 | 4.75E-10 |
| ENSG00000165124 | SVEP1 | 8.353700782 | 5.69E-10 |
| ENSG00000005001 | PRSS22 | 8.318069662 | 1.73E-09 |
| ENSG00000148795 | CYP17A1 | 8.262537503 | 5.28E-14 |
| ENSG00000122711 | SPINK4 | 8.000606445 | 4.70E-18 |
| ENSG00000167612 | ANKRD33 | 7.611463643 | 1.43E-07 |
| ENSG00000135472 | FAIM2 | 7.607682769 | 3.02E-07 |
| ENSG00000244588 | RAD21L1 | 7.56301205 | 1.56E-07 |
| ENSG00000189120 | SP6 | 7.502855543 | 1.24E-20 |
| ENSG00000073150 | PANX2 | 7.266836885 | 7.23E-07 |
| ENSG00000181523 | SGSH | 7.259222031 | 5.51E-15 |
| ENSG00000163395 | IGFN1 | 7.082956253 | 7.01E-06 |
| ENSG00000175311 | ANKS4B | 7.067554643 | 3.47E-06 |
| ENSG00000026751 | SLAMF7 | 7.00617529 | 4.43E-06 |
| ENSG00000164251 | F2RL1 | 6.999841234 | 2.36E-06 |
| ENSG00000130545 | CRB3 | 6.972929899 | 4.39E-07 |
| ENSG00000167580 | AQP2 | 6.963983832 | 2.59E-90 |
| ENSG00000135346 | CGA | 6.949403239 | 5.02E-07 |
| ENSG00000068615 | REEP1 | 6.917341816 | 8.00E-06 |
| ENSG00000198691 | ABCA4 | 6.815996028 | 1.02E-05 |
| ENSG00000079393 | DUSP13 | 6.799710717 | 1.56E-06 |
| ENSG00000291052 | ABCC13 | 6.779664409 | 1.33E-05 |
| ENSG00000141639 | MAPK4 | 6.771191137 | 3.71E-05 |
| ENSG00000160145 | KALRN | 6.742144461 | 7.32E-12 |
| ENSG00000131409 | LRRC4B | 6.69996735 | 2.76E-18 |
| ENSG00000261514 | LINC01976 | 6.611656593 | 3.73E-05 |
| ENSG00000127129 | EDN2 | 6.574669816 | 0.00015 |
| ENSG00000167914 | GSDMA | 6.568505736 | 3.77E-05 |
| ENSG00000248323 | LUCAT1 | 6.565029762 | 0.0001 |
| ENSG00000162654 | GBP4 | 6.552298003 | 0.000185 |
| ENSG00000081041 | CXCL2 | 6.537776895 | 9.10E-05 |
| ENSG00000278965 | gene_sourcehavana | 6.528848392 | 8.05E-05 |

| | | | |
|---|---|---|---|
| ENSG00000219438 | TAFA5 | 6.507318711 | 5.95E-05 |
| ENSG00000095739 | BAMBI | 6.48256978 | 9.31E-05 |
| ENSG00000288718 | gene_sourcehavana | 6.456618174 | 8.30E-05 |
| ENSG00000263818 | RDM1P5 | 6.405287637 | 9.59E-05 |
| ENSG00000187944 | C2orf66 | 6.354622144 | 0.000139 |
| ENSG00000144130 | NT5DC4 | 6.338781398 | 0.00021 |
| ENSG00000181045 | SLC26A11 | 6.330209323 | 0.000184 |
| ENSG00000171303 | KCNK3 | 6.289801012 | 0.00021 |
| ENSG00000257948 | gene_sourcehavana | 6.280207201 | 0.000209 |
| ENSG00000180537 | RNF182 | 6.237359889 | 0.000345 |
| ENSG00000166268 | MYRFL | 6.217769617 | 0.000321 |
| ENSG00000152315 | KCNK13 | 6.183293003 | 3.44E-18 |
| ENSG00000154451 | GBP5 | 6.181575921 | 0.001176 |
| ENSG00000233143 | DIRC3-AS1 | 6.143214899 | 0.00046 |
| ENSG00000137491 | SLCO2B1 | 6.140748971 | 4.03E-24 |
| ENSG00000289497 | gene_sourcehavana_ | 6.129585405 | 0.000481 |
| ENSG00000112139 | MDGA1 | 6.102943872 | 5.66E-07 |
| ENSG00000111732 | AICDA | 6.094353671 | 8.32E-05 |
| ENSG00000182170 | MRGPRG | 6.066904544 | 0.000692 |
| ENSG00000169439 | SDC2 | 5.955891203 | 0.001264 |
| ENSG00000111907 | TPD52L1 | 5.942404672 | 1.84E-31 |
| ENSG00000279365 | gene_sourceensembl | 5.924063505 | 0.00015 |
| ENSG00000223601 | EBLN1 | 5.919618144 | 0.001471 |
| ENSG00000112902 | SEMA5A | 5.917968472 | 0.001815 |
| ENSG00000150201 | FXYD4 | 5.899989203 | 4.85E-16 |
| ENSG00000164530 | PI16 | 5.899634605 | 0.001266 |
| ENSG00000259477 | gene_sourcehavana | 5.876715949 | 0.001623 |
| ENSG00000276070 | CCL4L2 | 5.872037819 | 4.93E-28 |
| ENSG00000103196 | CRISPLD2 | 5.871361891 | 1.76E-14 |
| ENSG00000226252 | gene_sourcehavana | 5.84318994 | 0.000182 |
| ENSG00000120875 | DUSP4 | 5.839535728 | 3.32E-70 |
| ENSG00000168081 | PNOC | 5.833672611 | 0.004419 |
| ENSG00000251230 | MIR3945HG | 5.824552489 | 5.09E-06 |
| ENSG00000205786 | LINC01531 | 5.819489276 | 0.002608 |
| ENSG00000229981 | LINC01435 | 5.816681731 | 0.002551 |
| ENSG00000187957 | DNER | 5.816647963 | 0.002076 |
| ENSG00000166866 | MYO1A | 5.793558883 | 5.62E-41 |
| ENSG00000291233 | gene_sourceensembl | 5.788504995 | 7.98E-06 |
| ENSG00000112182 | BACH2 | 5.777422392 | 0.001914 |
| ENSG00000115525 | ST3GAL5 | 5.761556948 | 0.000399 |
| ENSG00000182261 | NLRP10 | 5.757363093 | 0.00334 |
| ENSG00000007174 | DNAH9 | 5.74991217 | 0.003505 |
| ENSG00000086717 | PPEF1 | 5.746093953 | 1.44E-30 |
| ENSG00000091656 | ZFHX4 | 5.731571159 | 0.002708 |
| ENSG00000268324 | LRRC2-AS1 | 5.725779129 | 0.002616 |
| ENSG00000101265 | RASSF2 | 5.717195444 | 0.002842 |
| ENSG00000027644 | INSRR | 5.70889175 | 0.000303 |
| ENSG00000138316 | ADAMTS14 | 5.698614291 | 0.00484 |
| ENSG00000120913 | PDLIM2 | 5.659370821 | 0.004096 |

| | | | |
|---|---|---|---|
| ENSG00000187908 | DMBT1 | 5.649869223 | 0.0044 |
| ENSG00000233750 | CICP27 | 5.643661879 | 0.00346 |
| ENSG00000204577 | LILRB3 | 5.643641635 | 0.003515 |
| ENSG00000262454 | MIR193BHG | 5.636411406 | 0.00055 |
| ENSG00000175535 | PNLIP | 5.61227017 | 0.005835 |
| ENSG00000284624 | gene_sourcehavana | 5.607141921 | 0.006571 |
| ENSG00000176641 | RNF152 | 5.585614267 | 0.006702 |
| ENSG00000101680 | LAMA1 | 5.583459236 | 0.001661 |
| ENSG00000127324 | TSPAN8 | 5.567023401 | 0.000797 |
| ENSG00000184344 | GDF3 | 5.559163314 | 0.004662 |
| ENSG00000285179 | gene_sourcehavana | 5.556884337 | 0.005291 |
| ENSG00000104368 | PLAT | 5.533548923 | 0.007642 |
| ENSG00000147041 | SYTL5 | 5.533378458 | 0.010704 |
| ENSG00000112541 | PDE10A | 5.514296298 | 0.005671 |
| ENSG00000071909 | MYO3B | 5.509063991 | 0.001025 |
| ENSG00000272398 | CD24 | 5.505657995 | 1.99E-08 |
| ENSG00000280241 | gene_sourcehavana | 5.490750559 | 0.006357 |
| ENSG00000112796 | ENPP5 | 5.489624625 | 6.69E-13 |
| ENSG00000135269 | TES | 5.485216579 | 0.002105 |
| ENSG00000289085 | gene_sourcehavana_ | 5.479714883 | 6.64E-05 |
| ENSG00000072657 | TRHDE | 5.452488916 | 0.016362 |
| ENSG00000071246 | VASH1 | 5.448222197 | 4.05E-141 |
| ENSG00000126266 | FFAR1 | 5.439657772 | 0.001166 |
| ENSG00000125618 | PAX8 | 5.439249933 | 0.00777 |
| ENSG00000290888 | gene_sourcehavana | 5.436499863 | 0.009775 |
| ENSG00000075275 | CELSR1 | 5.428440008 | 1.82E-164 |
| ENSG00000103310 | ZP2 | 5.427743797 | 0.00992 |
| ENSG00000104093 | DMXL2 | 5.423581794 | 5.06E-07 |
| ENSG00000183696 | UPP1 | 5.413964676 | 0.001538 |
| ENSG00000170074 | FAM153A | 5.404731042 | 0.012099 |
| ENSG00000074410 | CA12 | 5.380280918 | 0.000137 |
| ENSG00000287374 | gene_sourcehavana_ | 5.369244522 | 0.009635 |
| ENSG00000239887 | C1orf226 | 5.362403717 | 0.011341 |
| ENSG00000289423 | gene_sourcehavana_ | 5.355514148 | 0.009087 |
| ENSG00000289621 | gene_sourcehavana_ | 5.341429402 | 0.010898 |
| ENSG00000163435 | ELF3 | 5.331064401 | 2.21E-06 |
| ENSG00000288738 | gene_sourcehavana_ | 5.329301592 | 1.04E-06 |
| ENSG00000081923 | ATP8B1 | 5.328819767 | 1.67E-70 |
| ENSG00000228252 | COL6A4P2 | 5.321706242 | 0.012409 |
| ENSG00000158270 | COLEC12 | 5.311644799 | 0.011585 |
| ENSG00000261888 | gene_sourcehavana | 5.306977558 | 0.011971 |
| ENSG00000163803 | PLB1 | 5.286217448 | 0.015504 |
| ENSG00000197506 | SLC28A3 | 5.273555732 | 4.38E-10 |
| ENSG00000198142 | SOWAHC | 5.267246842 | 7.48E-29 |
| ENSG00000057704 | TMCC3 | 5.263458739 | 0.001926 |
| ENSG00000198756 | COLGALT2 | 5.262924301 | 0.012604 |
| ENSG00000284722 | gene_sourcehavana | 5.257657941 | 0.000135 |
| ENSG00000107742 | SPOCK2 | 5.249591815 | 5.77E-08 |
| ENSG00000188906 | LRRK2 | 5.239070892 | 0.000127 |

| | | | |
|---|---|---|---|
| ENSG00000235750 | KIAA0040 | 5.236099933 | 0.022383 |
| ENSG00000168939 | SPRY3 | 5.235245845 | 0.028086 |
| ENSG00000130518 | IQCN | 5.229851407 | 3.10E-10 |
| ENSG00000109101 | FOXN1 | 5.209531155 | 0.015815 |
| ENSG00000269952 | gene_sourcehavana | 5.207974171 | 0.019779 |
| ENSG00000164398 | ACSL6 | 5.207042026 | 3.07E-27 |
| ENSG00000012223 | LTF | 5.198051596 | 1.10E-05 |
| ENSG00000163825 | RTP3 | 5.192057016 | 0.000279 |
| ENSG00000103316 | CRYM | 5.188139035 | 9.01E-49 |
| ENSG00000253686 | LINC01484 | 5.186747324 | 0.032404 |
| ENSG00000108849 | PPY | 5.186224623 | 0.032104 |
| ENSG00000198515 | CNGA1 | 5.176892901 | 0.016453 |
| ENSG00000289286 | gene_sourcehavana_ | 5.153989499 | 0.040783 |
| ENSG00000262580 | gene_sourcehavana | 5.144271922 | 0.025118 |
| ENSG00000274719 | gene_sourcehavana | 5.142149265 | 4.79E-06 |
| ENSG00000184985 | SORCS2 | 5.138456398 | 3.27E-06 |
| ENSG00000278498 | gene_sourceensembl | 5.130652433 | 0.019841 |
| ENSG00000277363 | SRCIN1 | 5.125501456 | 0.024434 |
| ENSG00000290021 | gene_sourcehavana_ | 5.111660638 | 9.65E-08 |
| ENSG00000255057 | gene_sourcehavana | 5.108788696 | 0.023891 |
| ENSG00000167618 | LAIR2 | 5.081023738 | 0.039954 |
| ENSG00000213981 | gene_sourcehavana | 5.075747898 | 0.022716 |
| ENSG00000135144 | DTX1 | 5.072449483 | 0.027779 |
| ENSG00000178199 | ZC3H12D | 5.072449483 | 0.027779 |
| ENSG00000275756 | gene_sourcehavana | 5.064192076 | 0.026524 |
| ENSG00000177993 | ZNRF3-AS1 | 5.057461489 | 0.007851 |
| ENSG00000176753 | PAK6-AS1 | 5.041488196 | 0.02669 |
| ENSG00000161798 | AQP5 | 5.040717818 | 0.030773 |
| ENSG00000101470 | TNNC2 | 5.03184598 | 0.024846 |
| ENSG00000227920 | gene_sourcehavana | 5.02355665 | 0.007864 |
| ENSG00000103089 | FA2H | 4.998131218 | 1.69E-05 |
| ENSG00000139151 | PLCZ1 | 4.996083933 | 0.006435 |
| ENSG00000217646 | H2BC16P | 4.993116702 | 0.03818 |
| ENSG00000254862 | LGR4-AS1 | 4.993116702 | 0.03818 |
| ENSG00000078114 | NEBL | 4.992660665 | 0.028996 |
| ENSG00000105427 | CNFN | 4.992660665 | 0.028996 |
| ENSG00000115896 | PLCL1 | 4.989468238 | 3.89E-14 |
| ENSG00000289419 | gene_sourcehavana_ | 4.987735388 | 0.006975 |
| ENSG00000065809 | FAM107B | 4.987044951 | 3.13E-13 |
| ENSG00000143365 | RORC | 4.981493702 | 0.03481 |
| ENSG00000082781 | ITGB5 | 4.981165871 | 2.22E-35 |
| ENSG00000175879 | HOXD8 | 4.962541141 | 0.03026 |
| ENSG00000187867 | PALM3 | 4.957412245 | 0.000831 |
| ENSG00000212296 | SNORD72 | 4.950850682 | 0.038287 |
| ENSG00000145416 | MARCHF1 | 4.943907779 | 7.07E-12 |
| ENSG00000221520 | MIR1285-1 | 4.9258101 | 0.032901 |
| ENSG00000132932 | ATP8A2 | 4.916474441 | 2.66E-10 |
| ENSG00000234869 | LINC02925 | 4.913349022 | 0.038239 |
| ENSG00000264672 | SEPTIN4-AS1 | 4.912265053 | 0.019073 |

| | | | |
|---|---|---|---|
| ENSG00000204815 | ODAD4 | 4.90272711 | 5.63E-74 |
| ENSG00000142627 | EPHA2 | 4.891220732 | 2.03E-05 |
| ENSG00000171136 | RLN3 | 4.889053059 | 0.008379 |
| ENSG00000124827 | GCM2 | 4.882801673 | 1.46E-09 |
| ENSG00000285010 | OOSP4A | 4.877806226 | 0.044611 |
| ENSG00000127528 | KLF2 | 4.87538447 | 6.57E-14 |
| ENSG00000275234 | gene_sourcehavana | 4.873587403 | 0.002886 |
| ENSG00000251076 | gene_sourcehavana | 4.854196365 | 3.97E-36 |
| ENSG00000167306 | MYO5B | 4.836847052 | 3.21E-94 |
| ENSG00000286431 | gene_sourcehavana_ | 4.833310148 | 0.001743 |
| ENSG00000226289 | CDCA4P3 | 4.831366398 | 0.042787 |
| ENSG00000264931 | MIR3138 | 4.822332455 | 0.049146 |
| ENSG00000270806 | C17orf50 | 4.816390918 | 0.020102 |
| ENSG00000283646 | LINC02009 | 4.794488526 | 5.22E-38 |
| ENSG00000167244 | IGF2 | 4.791112561 | 0.013112 |
| ENSG00000064886 | CHI3L2 | 4.762979118 | 0.013944 |
| ENSG00000168032 | ENTPD3 | 4.750972155 | 0.012912 |
| ENSG00000101892 | ATP1B4 | 4.746411158 | 1.59E-89 |
| ENSG00000109610 | SOD3 | 4.725171751 | 0.013832 |
| ENSG00000128918 | ALDH1A2 | 4.715570191 | 2.32E-48 |
| ENSG00000166049 | PASD1 | 4.712495653 | 5.94E-11 |
| ENSG00000107719 | PALD1 | 4.68927868 | 1.68E-25 |
| ENSG00000166863 | TAC3 | 4.683124923 | 9.75E-50 |
| ENSG00000166689 | PLEKHA7 | 4.675002354 | 6.16E-22 |
| ENSG00000105550 | FGF21 | 4.669788511 | 0.014213 |
| ENSG00000131620 | ANO1 | 4.666167242 | 0.000323 |
| ENSG00000101445 | PPP1R16B | 4.66398733 | 0.01987 |
| ENSG00000144645 | OSBPL10 | 4.663417826 | 8.92E-13 |
| ENSG00000171729 | TMEM51 | 4.649875237 | 0.020086 |
| ENSG00000175793 | SFN | 4.641709398 | 3.54E-09 |
| ENSG00000171346 | KRT15 | 4.617902762 | 2.33E-10 |
| ENSG00000113916 | BCL6 | 4.616584592 | 4.78E-74 |
| ENSG00000122176 | FMOD | 4.615630832 | 8.39E-41 |
| ENSG00000162687 | KCNT2 | 4.608960834 | 0.001799 |
| ENSG00000173482 | PTPRM | 4.598353262 | 3.51E-09 |
| ENSG00000271737 | gene_sourcehavana | 4.592732012 | 0.019342 |
| ENSG00000125735 | TNFSF14 | 4.592452371 | 3.46E-59 |
| ENSG00000177675 | CD163L1 | 4.588846762 | 0.031464 |
| ENSG00000130762 | ARHGEF16 | 4.584147943 | 5.31E-14 |
| ENSG00000141431 | ASXL3 | 4.56487974 | 5.84E-24 |
| ENSG00000248559 | gene_sourcehavana | 4.549339121 | 0.024173 |
| ENSG00000173083 | HPSE | 4.539669871 | 2.13E-50 |
| ENSG00000224186 | PITX1-AS1 | 4.5177153 | 7.77E-111 |
| ENSG00000133321 | PLAAT4 | 4.494871222 | 0.032652 |
| ENSG00000239801 | DENND6A-AS1 | 4.454119221 | 0.034503 |
| ENSG00000205038 | PKHD1L1 | 4.445569985 | 2.24E-17 |
| ENSG00000226510 | UPK1A-AS1 | 4.44552782 | 0.00483 |
| ENSG00000216901 | ZNF603P | 4.442983521 | 0.027211 |
| ENSG00000279108 | gene_sourcehavana | 4.432377528 | 0.007044 |

| | | | |
|---|---|---|---|
| ENSG00000144366 | GULP1 | 4.431685185 | 8.44E-09 |
| ENSG00000164185 | ZNF474 | 4.425343405 | 0.004569 |
| ENSG00000104892 | KLC3 | 4.422998089 | 1.83E-05 |
| ENSG00000184698 | OR51M1 | 4.421096731 | 0.030593 |
| ENSG00000237596 | PDE7B-AS1 | 4.41698214 | 3.75E-05 |
| ENSG00000175093 | SPSB4 | 4.407608853 | 0.031304 |
| ENSG00000254815 | LMNTD2-AS1 | 4.38526868 | 0.035919 |
| ENSG00000117114 | ADGRL2 | 4.382388692 | 0.049809 |
| ENSG00000158258 | CLSTN2 | 4.362678032 | 2.58E-07 |
| ENSG00000023445 | BIRC3 | 4.358627077 | 5.83E-15 |
| ENSG00000287860 | gene_sourcehavana_ | 4.357741693 | 2.31E-13 |
| ENSG00000086967 | MYBPC2 | 4.352956468 | 0.043657 |
| ENSG00000260641 | TSPAN5-DT | 4.351097987 | 1.45E-05 |
| ENSG00000115423 | DNAH6 | 4.347689663 | 0.049202 |
| ENSG00000107295 | SH3GL2 | 4.343175954 | 0.034725 |
| ENSG00000125730 | C3 | 4.340602643 | 2.30E-20 |
| ENSG00000079102 | RUNX1T1 | 4.33719163 | 4.78E-33 |
| ENSG00000204010 | IFIT1B | 4.333040426 | 0.002642 |
| ENSG00000079101 | CLUL1 | 4.328631197 | 1.29E-07 |
| ENSG00000227331 | RPL7AP22 | 4.323671808 | 0.007833 |
| ENSG00000074219 | TEAD2 | 4.314884036 | 0.040645 |
| ENSG00000288754 | gene_sourcehavana_ | 4.309613274 | 0.034153 |
| ENSG00000197653 | DNAH10 | 4.300046282 | 0.045041 |
| ENSG00000196188 | CTSE | 4.289446652 | 1.48E-69 |
| ENSG00000236095 | gene_sourcehavana | 4.276731606 | 0.047267 |
| ENSG00000270095 | gene_sourcehavana | 4.270786946 | 0.045917 |
| ENSG00000006747 | SCIN | 4.250996682 | 0.034493 |
| ENSG00000260919 | gene_sourcehavana | 4.230619253 | 0.009776 |
| ENSG00000140600 | SH3GL3 | 4.224063064 | 1.08E-10 |
| ENSG00000239998 | LILRA2 | 4.213027136 | 3.73E-06 |
| ENSG00000196565 | HBG2 | 4.212996333 | 1.20E-09 |
| ENSG00000104921 | FCER2 | 4.203692667 | 0.004337 |
| ENSG00000089472 | HEPH | 4.190682432 | 8.59E-37 |
| ENSG00000278952 | gene_sourcehavana | 4.177834562 | 0.002977 |
| ENSG00000108852 | MPP2 | 4.175831226 | 6.29E-51 |
| ENSG00000269246 | gene_sourcehavana | 4.170747232 | 0.000361 |
| ENSG00000130513 | GDF15 | 4.156588005 | 3.29E-31 |
| ENSG00000285870 | gene_sourcehavana | 4.148504011 | 0.016089 |
| ENSG00000171561 | OR2AT4 | 4.145994393 | 7.61E-09 |
| ENSG00000290003 | gene_sourcehavana_ | 4.116618877 | 0.013208 |
| ENSG00000060566 | CREB3L3 | 4.111743309 | 1.08E-34 |
| ENSG00000233435 | AGGF1P2 | 4.110582826 | 0.013208 |
| ENSG00000250359 | PTP4A1P4 | 4.088899896 | 7.69E-12 |
| ENSG00000166592 | RRAD | 4.086290643 | 8.82E-13 |
| ENSG00000204060 | FOXO6 | 4.085400858 | 3.20E-08 |
| ENSG00000136052 | SLC41A2 | 4.085397688 | 0.000199 |
| ENSG00000113083 | LOX | 4.084495868 | 1.47E-73 |
| ENSG00000133687 | TMTC1 | 4.082032954 | 7.40E-09 |
| ENSG00000131831 | RAI2 | 4.081791007 | 0.004706 |

| | | | |
|---|---|---|---|
| ENSG00000260629 | BGLT3 | 4.04749518 | 4.47E-05 |
| ENSG00000007384 | RHBDF1 | 4.040219597 | 7.62E-08 |
| ENSG00000120057 | SFRP5 | 4.03212499 | 1.88E-61 |
| ENSG00000012124 | CD22 | 4.023615674 | 4.82E-21 |
| ENSG00000101298 | SNPH | 4.022534878 | 1.15E-68 |
| ENSG00000188385 | JAKMIP3 | 4.02184762 | 0.000186 |
| ENSG00000164199 | ADGRV1 | 4.010508379 | 0.006276 |
| ENSG00000259655 | gene_sourcehavana | 3.992818387 | 0.026757 |
| ENSG00000198794 | SCAMP5 | 3.984038248 | 0.000978 |
| ENSG00000162825 | NBPF20 | 3.973458854 | 2.82E-16 |
| ENSG00000142408 | CACNG8 | 3.950248894 | 2.32E-06 |
| ENSG00000223865 | HLA-DPB1 | 3.941962447 | 2.02E-06 |
| ENSG00000164128 | NPY1R | 3.93962256 | 7.46E-289 |
| ENSG00000290808 | gene_sourcehavana | 3.938888491 | 2.15E-06 |
| ENSG00000166535 | A2ML1 | 3.92550175 | 3.54E-14 |
| ENSG00000254732 | gene_sourcehavana | 3.917705535 | 0.00722 |
| ENSG00000186510 | CLCNKA | 3.915298891 | 0.011592 |
| ENSG00000123094 | RASSF8 | 3.911582503 | 4.79E-07 |
| ENSG00000229005 | HNF4A-AS1 | 3.905850898 | 0.0001 |
| ENSG00000255760 | LINC02422 | 3.894337511 | 9.45E-06 |
| ENSG00000135547 | HEY2 | 3.889060257 | 0.000436 |
| ENSG00000134594 | RAB33A | 3.882801784 | 0.005542 |
| ENSG00000246225 | gene_sourcehavana | 3.872210044 | 0.035005 |
| ENSG00000249026 | CTNNA1P1 | 3.851264408 | 0.030479 |
| ENSG00000139289 | PHLDA1 | 3.848146673 | 2.62E-34 |
| ENSG00000136826 | KLF4 | 3.847027197 | 6.48E-19 |
| ENSG00000088756 | ARHGAP28 | 3.844693221 | 0.00123 |
| ENSG00000121653 | MAPK8IP1 | 3.840158657 | 3.16E-28 |
| ENSG00000140836 | ZFHX3 | 3.816752381 | 3.36E-09 |
| ENSG00000111052 | LIN7A | 3.808179382 | 1.09E-14 |
| ENSG00000185669 | SNAI3 | 3.799351934 | 9.62E-14 |
| ENSG00000159409 | CELF3 | 3.788905 | 1.25E-20 |
| ENSG00000023171 | GRAMD1B | 3.785958046 | 0.045567 |
| ENSG00000168126 | OR2W6P | 3.783908857 | 6.91E-05 |
| ENSG00000148926 | ADM | 3.77925536 | 0.000489 |
| ENSG00000117525 | F3 | 3.768149201 | 0.000166 |
| ENSG00000118200 | CAMSAP2 | 3.764603646 | 8.89E-12 |
| ENSG00000068985 | PAGE1 | 3.733101908 | 0.006307 |
| ENSG00000268785 | RPL7P50 | 3.695256565 | 0.048028 |
| ENSG00000188158 | NHS | 3.665606697 | 0.023199 |
| ENSG00000108846 | ABCC3 | 3.664699006 | 0.027604 |
| ENSG00000161921 | CXCL16 | 3.653066209 | 2.12E-08 |
| ENSG00000095397 | WHRN | 3.652529392 | 5.90E-16 |
| ENSG00000266934 | gene_sourcehavana | 3.6414144 | 0.029274 |
| ENSG00000239557 | PPP2R5CP | 3.638978484 | 0.017975 |
| ENSG00000196660 | SLC30A10 | 3.629506768 | 3.49E-116 |
| ENSG00000101162 | TUBB1 | 3.623087627 | 2.39E-106 |
| ENSG00000204421 | LY6G6C | 3.623038702 | 3.70E-05 |
| ENSG00000118785 | SPP1 | 3.619085214 | 0.013192 |

| | | | |
|---|---|---|---|
| ENSG00000282757 | DUXB | 3.615978355 | 0.000112 |
| ENSG00000288825 | H2AC18 | 3.613847614 | 0.0306 |
| ENSG00000147081 | AKAP4 | 3.609085376 | 7.18E-22 |
| ENSG00000270959 | LPP-AS2 | 3.605277152 | 1.46E-09 |
| ENSG00000184545 | DUSP8 | 3.57275845 | 9.53E-06 |
| ENSG00000275527 | gene_sourcehavana | 3.572292878 | 1.55E-63 |
| ENSG00000240497 | gene_sourcehavana | 3.552046961 | 0.034437 |
| ENSG00000263874 | LASP1NB | 3.551582358 | 0.022814 |
| ENSG00000204131 | NHSL2 | 3.547917152 | 0.028769 |
| ENSG00000291138 | gene_sourcehavana | 3.545082775 | 0.002432 |
| ENSG00000213126 | TXNL4AP1 | 3.544177605 | 5.20E-26 |
| ENSG00000206178 | HBZP1 | 3.540062687 | 0.016257 |
| ENSG00000229056 | HECW2-AS1 | 3.535476731 | 8.84E-09 |
| ENSG00000101210 | EEF1A2 | 3.526064693 | 5.16E-75 |
| ENSG00000133019 | CHRM3 | 3.517080638 | 0.000174 |
| ENSG00000142235 | LMTK3 | 3.515444631 | 4.98E-09 |
| ENSG00000224083 | MTCO1P11 | 3.497566479 | 0.004868 |
| ENSG00000011028 | MRC2 | 3.496315849 | 5.82E-68 |
| ENSG00000103855 | CD276 | 3.472926062 | 1.69E-88 |
| ENSG00000241135 | LINC00881 | 3.47137556 | 0.013405 |
| ENSG00000130656 | HBZ | 3.463608867 | 3.65E-99 |
| ENSG00000287632 | gene_sourcehavana_ | 3.462022325 | 2.69E-06 |
| ENSG00000162148 | PPP1R32 | 3.459944498 | 0.016796 |
| ENSG00000170500 | LONRF2 | 3.459650359 | 6.73E-62 |
| ENSG00000239219 | FHL1P1 | 3.451892267 | 1.32E-55 |
| ENSG00000185112 | FAM43A | 3.446472654 | 1.26E-06 |
| ENSG00000115594 | IL1R1 | 3.445204242 | 5.61E-111 |
| ENSG00000162745 | OLFML2B | 3.433052905 | 0.032853 |
| ENSG00000143369 | ECM1 | 3.416338506 | 7.73E-86 |
| ENSG00000184489 | PTP4A3 | 3.412136959 | 4.47E-10 |
| ENSG00000289331 | gene_sourcehavana_ | 3.40839598 | 0.002937 |
| ENSG00000268621 | IGFL2-AS1 | 3.403049156 | 6.68E-28 |
| ENSG00000104213 | PDGFRL | 3.394146664 | 0.00771 |
| ENSG00000108309 | RUNDC3A | 3.386301698 | 1.06E-36 |
| ENSG00000188175 | HEPACAM2 | 3.38143464 | 9.57E-31 |
| ENSG00000227230 | gene_sourcehavana | 3.374790943 | 7.96E-05 |
| ENSG00000168994 | PXDC1 | 3.36329348 | 0.00023 |
| ENSG00000226711 | FAM66C | 3.358539094 | 0.048974 |
| ENSG00000171119 | NRTN | 3.358326052 | 1.43E-10 |
| ENSG00000272905 | gene_sourcehavana | 3.346244499 | 0.01531 |
| ENSG00000058335 | RASGRF1 | 3.343902536 | 0.017761 |
| ENSG00000170271 | FAXDC2 | 3.339954629 | 5.28E-58 |
| ENSG00000143341 | HMCN1 | 3.332700263 | 1.17E-39 |
| ENSG00000249947 | XBP1P1 | 3.322747095 | 0.045745 |
| ENSG00000186642 | PDE2A | 3.316080561 | 0.006594 |
| ENSG00000238365 | RNU7-57P | 3.305458538 | 2.71E-06 |
| ENSG00000196169 | KIF19 | 3.304836303 | 0.010156 |
| ENSG00000268812 | LIF-AS2 | 3.298001744 | 1.28E-05 |
| ENSG00000115902 | SLC1A4 | 3.290916347 | 1.21E-27 |

| | | | |
|---|---|---|---|
| ENSG00000131080 | EDA2R | 3.282647221 | 0.005071 |
| ENSG00000264175 | MIR3189 | 3.280285353 | 1.75E-06 |
| ENSG00000078900 | TP73 | 3.27554889 | 1.61E-07 |
| ENSG00000132975 | GPR12 | 3.271635569 | 5.13E-06 |
| ENSG00000148400 | NOTCH1 | 3.261568223 | 3.57E-11 |
| ENSG00000135083 | CCNJL | 3.249340131 | 1.30E-09 |
| ENSG00000238009 | gene_sourceensembl | 3.247760012 | 0.00135 |
| ENSG00000171345 | KRT19 | 3.246406674 | 2.29E-26 |
| ENSG00000114923 | SLC4A3 | 3.244921345 | 2.20E-11 |
| ENSG00000186522 | SEPTIN10 | 3.238164331 | 4.77E-17 |
| ENSG00000112280 | COL9A1 | 3.23616153 | 0.000143 |
| ENSG00000007516 | BAIAP3 | 3.22759495 | 5.94E-40 |
| ENSG00000255367 | gene_sourcehavana | 3.214110339 | 2.01E-11 |
| ENSG00000276600 | RAB7B | 3.21364694 | 1.92E-07 |
| ENSG00000215704 | CELA2B | 3.211625656 | 5.86E-17 |
| ENSG00000105699 | LSR | 3.210475462 | 3.43E-34 |
| ENSG00000239494 | RN7SL333P | 3.20995271 | 0.037219 |
| ENSG00000088367 | EPB41L1 | 3.208336586 | 1.30E-23 |
| ENSG00000064393 | HIPK2 | 3.206909047 | 2.84E-99 |
| ENSG00000105048 | TNNT1 | 3.200507649 | 1.33E-40 |
| ENSG00000234614 | C2CD4D-AS1 | 3.199379692 | 2.56E-06 |
| ENSG00000101342 | TLDC2 | 3.191457642 | 2.37E-37 |
| ENSG00000236885 | gene_sourcehavana | 3.190869976 | 0.041499 |
| ENSG00000125462 | MIR9-1HG | 3.184064859 | 0.024047 |
| ENSG00000254231 | WWP1-AS1 | 3.183729207 | 0.003335 |
| ENSG00000166831 | RBPMS2 | 3.170195289 | 0.006482 |
| ENSG00000122986 | HVCN1 | 3.166058886 | 1.34E-05 |
| ENSG00000267750 | RUNDC3A-AS1 | 3.164965249 | 2.03E-51 |
| ENSG00000138772 | ANXA3 | 3.163357962 | 0.000321 |
| ENSG00000153234 | NR4A2 | 3.15443791 | 1.56E-28 |
| ENSG00000245293 | CYP2U1-AS1 | 3.153937699 | 0.000909 |
| ENSG00000275880 | NAXD-AS1 | 3.141948227 | 0.008636 |
| ENSG00000128242 | GAL3ST1 | 3.140105019 | 0.042077 |
| ENSG00000161653 | NAGS | 3.138379236 | 5.82E-05 |
| ENSG00000246731 | MGC16275 | 3.132304405 | 4.72E-40 |
| ENSG00000279613 | gene_sourcehavana | 3.11874214 | 0.009267 |
| ENSG00000111879 | FAM184A | 3.118336914 | 9.01E-24 |
| ENSG00000122224 | LY9 | 3.116723431 | 4.86E-12 |
| ENSG00000140678 | ITGAX | 3.108535912 | 0.000951 |
| ENSG00000282024 | gene_sourcehavana | 3.108273605 | 1.89E-38 |
| ENSG00000171759 | PAH | 3.09466072 | 0.022556 |
| ENSG00000124762 | CDKN1A | 3.092106961 | 5.08E-52 |
| ENSG00000196549 | MME | 3.088362443 | 1.32E-29 |
| ENSG00000183578 | TNFAIP8L3 | 3.080784371 | 0.011977 |
| ENSG00000167286 | CD3D | 3.078487586 | 2.52E-09 |
| ENSG00000188582 | PAQR9 | 3.07404907 | 9.93E-107 |
| ENSG00000287906 | gene_sourcehavana_ | 3.073597847 | 8.70E-06 |
| ENSG00000240583 | AQP1 | 3.065266424 | 1.69E-42 |
| ENSG00000110848 | CD69 | 3.061113104 | 2.47E-77 |

| | | | |
|---|---|---|---|
| ENSG00000286672 | gene_sourcehavana_ | 3.048840858 | 5.39E-05 |
| ENSG00000118898 | PPL | 3.042231953 | 0.001988 |
| ENSG00000054356 | PTPRN | 3.038269871 | 0.002824 |
| ENSG00000204613 | TRIM10 | 3.020085595 | 1.54E-197 |
| ENSG00000096696 | DSP | 3.020042677 | 2.70E-05 |
| ENSG00000164855 | TMEM184A | 3.018787503 | 0.001785 |
| ENSG00000164023 | SGMS2 | 3.017220249 | 0.005389 |
| ENSG00000291067 | gene_sourcehavana | 3.01341497 | 2.66E-07 |
| ENSG00000120318 | ARAP3 | 3.00327152 | 5.55E-05 |
| ENSG00000087494 | PTHLH | 2.994407224 | 0.000169 |
| ENSG00000220517 | ASS1P1 | 2.983897857 | 0.01191 |
| ENSG00000135077 | HAVCR2 | 2.981578503 | 1.51E-18 |
| ENSG00000164683 | HEY1 | 2.981246447 | 6.29E-07 |
| ENSG00000228010 | gene_sourcehavana | 2.978179093 | 0.006551 |
| ENSG00000290793 | gene_sourcehavana | 2.975751183 | 0.043464 |
| ENSG00000070614 | NDST1 | 2.975337922 | 4.88E-58 |
| ENSG00000101076 | HNF4A | 2.96795046 | 0.026494 |
| ENSG00000181819 | KCTD9P2 | 2.967231167 | 0.030024 |
| ENSG00000136040 | PLXNC1 | 2.966113692 | 1.48E-64 |
| ENSG00000116299 | ELAPOR1 | 2.965279633 | 0.000266 |
| ENSG00000095303 | PTGS1 | 2.962616332 | 1.57E-122 |
| ENSG00000100600 | LGMN | 2.958350903 | 0.000126 |
| ENSG00000287338 | gene_sourcehavana_ | 2.954153706 | 8.34E-08 |
| ENSG00000154316 | TDH | 2.941213055 | 0.00049 |
| ENSG00000166960 | CCDC178 | 2.941143773 | 0.000211 |
| ENSG00000272791 | gene_sourcehavana | 2.93243605 | 0.021589 |
| ENSG00000272554 | gene_sourcehavana | 2.929704555 | 0.013704 |
| ENSG00000175505 | CLCF1 | 2.927832622 | 6.68E-15 |
| ENSG00000166016 | ABTB2 | 2.908203457 | 7.19E-56 |
| ENSG00000220867 | HSPE1P26 | 2.907381962 | 0.013655 |
| ENSG00000269970 | gene_sourcehavana | 2.896243472 | 0.000272 |
| ENSG00000271992 | gene_sourcehavana | 2.894520426 | 0.000339 |
| ENSG00000160588 | MPZL3 | 2.889533334 | 4.47E-25 |
| ENSG00000182752 | PAPPA | 2.889215848 | 2.36E-44 |
| ENSG00000198948 | MFAP3L | 2.885574232 | 3.17E-43 |
| ENSG00000188672 | RHCE | 2.883233603 | 4.30E-94 |
| ENSG00000169618 | PROKR1 | 2.878618263 | 0.009498 |
| ENSG00000288744 | gene_sourcehavana_ | 2.870954202 | 0.022492 |
| ENSG00000230479 | LNCTSI | 2.861954968 | 0.001021 |
| ENSG00000196139 | AKR1C3 | 2.855768426 | 2.16E-42 |
| ENSG00000163053 | SLC16A14 | 2.855445757 | 0.025688 |
| ENSG00000124657 | OR2B6 | 2.852186766 | 0.018178 |
| ENSG00000151117 | TMEM86A | 2.849078536 | 0.000417 |
| ENSG00000186529 | CYP4F3 | 2.845572973 | 1.99E-05 |
| ENSG00000138119 | MYOF | 2.836553226 | 0.040051 |
| ENSG00000128266 | GNAZ | 2.830030587 | 6.75E-08 |
| ENSG00000039560 | RAI14 | 2.829927149 | 1.16E-05 |
| ENSG00000079385 | CEACAM1 | 2.829497476 | 0.004656 |
| ENSG00000183690 | EFHC2 | 2.811967509 | 0.012478 |

| | | | |
|---|---|---|---|
| ENSG00000101605 | MYOM1 | 2.808757473 | 0.024042 |
| ENSG00000134070 | IRAK2 | 2.805942815 | 1.44E-53 |
| ENSG00000085491 | SLC25A24 | 2.804404193 | 0.001802 |
| ENSG00000233436 | BTBD18 | 2.798786786 | 0.000163 |
| ENSG00000284664 | gene_sourcehavana | 2.795500437 | 0.004577 |
| ENSG00000287315 | gene_sourcehavana | 2.78804411 | 0.033119 |
| ENSG00000166289 | PLEKHF1 | 2.773383671 | 2.86E-11 |
| ENSG00000260923 | LINC02193 | 2.770669385 | 0.001879 |
| ENSG00000187010 | RHD | 2.766473852 | 1.28E-105 |
| ENSG00000130203 | APOE | 2.766399342 | 1.30E-59 |
| ENSG00000291048 | gene_sourcehavana | 2.765458365 | 9.69E-09 |
| ENSG00000261349 | gene_sourcehavana | 2.765326039 | 1.07E-44 |
| ENSG00000148483 | TMEM236 | 2.757186458 | 2.25E-06 |
| ENSG00000095932 | SMIM24 | 2.755593936 | 1.44E-23 |
| ENSG00000100314 | CABP7 | 2.751267668 | 5.42E-51 |
| ENSG00000225891 | DHDDS-AS1 | 2.749899025 | 0.012115 |
| ENSG00000172508 | CARNS1 | 2.748638966 | 5.86E-124 |
| ENSG00000238249 | HMGN2P17 | 2.747233653 | 0.032853 |
| ENSG00000290032 | gene_sourcehavana_ | 2.744696323 | 1.02E-65 |
| ENSG00000175745 | NR2F1 | 2.743987072 | 0.016254 |
| ENSG00000109501 | WFS1 | 2.738641395 | 1.41E-10 |
| ENSG00000135678 | CPM | 2.724719964 | 1.71E-27 |
| ENSG00000161681 | SHANK1 | 2.724036236 | 2.82E-28 |
| ENSG00000126759 | CFP | 2.723104877 | 0.001791 |
| ENSG00000164949 | GEM | 2.723065895 | 8.87E-12 |
| ENSG00000253304 | TMEM200B | 2.719839019 | 1.79E-09 |
| ENSG00000261183 | SPINT1-AS1 | 2.717869915 | 0.016206 |
| ENSG00000111331 | OAS3 | 2.706985641 | 0.001067 |
| ENSG00000172403 | SYNPO2 | 2.706832907 | 0.01856 |
| ENSG00000158406 | H4C8 | 2.70620841 | 4.51E-27 |
| ENSG00000107518 | ATRNL1 | 2.701705942 | 0.004984 |
| ENSG00000180573 | H2AC6 | 2.695583462 | 1.93E-62 |
| ENSG00000287553 | gene_sourcehavana_ | 2.695267482 | 0.019266 |
| ENSG00000164181 | ELOVL7 | 2.694254056 | 2.42E-07 |
| ENSG00000162415 | ZSWIM5 | 2.689934781 | 2.41E-19 |
| ENSG00000232022 | FAAHP1 | 2.689916685 | 0.000453 |
| ENSG00000128283 | CDC42EP1 | 2.684725906 | 2.81E-21 |
| ENSG00000130208 | APOC1 | 2.683204393 | 5.54E-162 |
| ENSG00000128567 | PODXL | 2.681406019 | 1.10E-22 |
| ENSG00000213934 | HBG1 | 2.655733226 | 1.67E-14 |
| ENSG00000171401 | KRT13 | 2.655555704 | 8.32E-42 |
| ENSG00000290853 | gene_sourcehavana | 2.653871439 | 0.000105 |
| ENSG00000177173 | NAP1L4P1 | 2.64469374 | 1.31E-16 |
| ENSG00000169442 | CD52 | 2.63913524 | 7.56E-199 |
| ENSG00000198092 | TMPRSS11F | 2.629125212 | 2.28E-19 |
| ENSG00000149418 | ST14 | 2.629064946 | 7.26E-09 |
| ENSG00000086730 | LAT2 | 2.621479387 | 4.61E-28 |
| ENSG00000197279 | ZNF165 | 2.614184549 | 8.05E-116 |
| ENSG00000234350 | ERICH2-DT | 2.612572346 | 7.86E-30 |

| | | | |
|---|---|---|---|
| ENSG00000289915 | gene_sourcehavana_ | 2.603296536 | 1.11E-07 |
| ENSG00000173334 | TRIB1 | 2.59729321 | 1.33E-34 |
| ENSG00000179913 | B3GNT3 | 2.588660452 | 9.21E-06 |
| ENSG00000169194 | IL13 | 2.588503915 | 0.015463 |
| ENSG00000075399 | VPS9D1 | 2.58019594 | 6.71E-55 |
| ENSG00000213412 | HNRNPA1P33 | 2.565063716 | 2.97E-05 |
| ENSG00000259207 | ITGB3 | 2.560915177 | 0.005649 |
| ENSG00000133169 | BEX1 | 2.55919179 | 1.07E-103 |
| ENSG00000187837 | H1-2 | 2.550659046 | 2.87E-22 |
| ENSG00000105711 | SCN1B | 2.548475384 | 4.83E-05 |
| ENSG00000261158 | gene_sourcehavana | 2.546032766 | 0.031333 |
| ENSG00000181234 | TMEM132C | 2.545936198 | 0.000154 |
| ENSG00000125740 | FOSB | 2.545512655 | 0.000111 |
| ENSG00000104081 | BMF | 2.535752459 | 8.62E-36 |
| ENSG00000135407 | AVIL | 2.533264849 | 0.003733 |
| ENSG00000179403 | VWA1 | 2.532899176 | 1.24E-09 |
| ENSG00000176485 | PLAAT3 | 2.518231253 | 5.95E-06 |
| ENSG00000256706 | gene_sourcehavana | 2.513931627 | 1.56E-05 |
| ENSG00000197903 | H2BC12 | 2.51180102 | 3.31E-80 |
| ENSG00000181019 | NQO1 | 2.503675727 | 6.97E-49 |
| ENSG00000197712 | FAM114A1 | 2.497280621 | 6.87E-09 |
| ENSG00000135454 | B4GALNT1 | 2.495133044 | 3.44E-06 |
| ENSG00000282021 | gene_sourcehavana | 2.493804469 | 0.021898 |
| ENSG00000165029 | ABCA1 | 2.488887333 | 0.000113 |
| ENSG00000225528 | TMA7B | 2.479630909 | 1.10E-11 |
| ENSG00000249790 | LINC02972 | 2.475723414 | 2.61E-153 |
| ENSG00000187164 | SHTN1 | 2.474287166 | 2.04E-19 |
| ENSG00000270022 | gene_sourcehavana | 2.471997562 | 0.007469 |
| ENSG00000158014 | SLC30A2 | 2.470093818 | 1.36E-06 |
| ENSG00000239291 | gene_sourcehavana | 2.465371392 | 0.009832 |
| ENSG00000073910 | FRY | 2.463554316 | 1.70E-08 |
| ENSG00000180891 | CUEDC1 | 2.461364914 | 0.007131 |
| ENSG00000129422 | MTUS1 | 2.454593537 | 9.27E-05 |
| ENSG00000115756 | HPCAL1 | 2.450474896 | 4.46E-85 |
| ENSG00000291156 | gene_sourcehavana | 2.447195313 | 0.012909 |
| ENSG00000197721 | CR1L | 2.446694676 | 3.45E-17 |
| ENSG00000134376 | CRB1 | 2.445868823 | 0.001134 |
| ENSG00000165795 | NDRG2 | 2.445344581 | 0.000145 |
| ENSG00000186594 | MIR22HG | 2.443191106 | 3.37E-64 |
| ENSG00000105707 | HPN | 2.44025479 | 0.015253 |
| ENSG00000025039 | RRAGD | 2.439705997 | 2.78E-31 |
| ENSG00000108370 | RGS9 | 2.435743753 | 0.004538 |
| ENSG00000259030 | FPGT-TNNI3K | 2.435405375 | 0.045692 |
| ENSG00000166900 | STX3 | 2.434836711 | 2.45E-117 |
| ENSG00000067606 | PRKCZ | 2.430763102 | 1.02E-07 |
| ENSG00000157483 | MYO1E | 2.413977519 | 0.046478 |
| ENSG00000102934 | PLLP | 2.407539258 | 0.001046 |
| ENSG00000168246 | UBTD2 | 2.402012793 | 5.69E-05 |
| ENSG00000287190 | gene_sourcehavana_ | 2.398811106 | 0.037619 |

| | | | |
|---|---|---|---|
| ENSG00000187017 | ESPN | 2.39585448 | 2.02E-146 |
| ENSG00000166317 | SYNPO2L | 2.393105634 | 3.64E-06 |
| ENSG00000185697 | MYBL1 | 2.386139667 | 0.002046 |
| ENSG00000152767 | FARP1 | 2.383623234 | 1.49E-66 |
| ENSG00000156042 | CFAP70 | 2.382671177 | 1.31E-08 |
| ENSG00000138678 | GPAT3 | 2.381563287 | 0.000263 |
| ENSG00000174028 | FAM3C2P | 2.378842856 | 1.15E-19 |
| ENSG00000082074 | FYB1 | 2.372297617 | 0.048359 |
| ENSG00000260793 | gene_sourcehavana | 2.370997785 | 2.73E-16 |
| ENSG00000229953 | BCAN-AS2 | 2.368598157 | 0.028421 |
| ENSG00000175471 | MCTP1 | 2.367377661 | 0.044753 |
| ENSG00000008735 | MAPK8IP2 | 2.36720231 | 0.036901 |
| ENSG00000122694 | GLIPR2 | 2.36654731 | 1.35E-42 |
| ENSG00000139625 | MAP3K12 | 2.364364814 | 1.25E-10 |
| ENSG00000130202 | NECTIN2 | 2.362502092 | 4.29E-34 |
| ENSG00000134827 | TCN1 | 2.362085263 | 2.42E-08 |
| ENSG00000164841 | TMEM74 | 2.36183157 | 2.90E-12 |
| ENSG00000024422 | EHD2 | 2.358898736 | 1.90E-101 |
| ENSG00000120693 | SMAD9 | 2.350729676 | 2.82E-11 |
| ENSG00000137094 | DNAJB5 | 2.346063158 | 0.000159 |
| ENSG00000135709 | KIAA0513 | 2.343386978 | 1.89E-27 |
| ENSG00000152952 | PLOD2 | 2.340051829 | 0.000182 |
| ENSG00000181104 | F2R | 2.337299667 | 1.04E-40 |
| ENSG00000080561 | MID2 | 2.336199496 | 3.86E-14 |
| ENSG00000072182 | ASIC4 | 2.330838813 | 3.94E-39 |
| ENSG00000232600 | TONSL-AS1 | 2.329144119 | 0.000222 |
| ENSG00000128872 | TMOD2 | 2.325809428 | 0.028929 |
| ENSG00000166750 | SLFN5 | 2.325198026 | 5.49E-14 |
| ENSG00000131242 | RAB11FIP4 | 2.319175339 | 6.51E-44 |
| ENSG00000279283 | gene_sourcehavana | 2.317550167 | 0.004055 |
| ENSG00000073756 | PTGS2 | 2.315285283 | 0.045498 |
| ENSG00000241570 | PAQR9-AS1 | 2.312430296 | 0.041331 |
| ENSG00000228463 | gene_sourcehavana | 2.306619217 | 6.83E-09 |
| ENSG00000074657 | ZNF532 | 2.305868482 | 0.000202 |
| ENSG00000178301 | AQP11 | 2.305147557 | 6.47E-07 |
| ENSG00000159958 | TNFRSF13C | 2.298492117 | 0.011432 |
| ENSG00000078246 | TULP3 | 2.297843532 | 2.15E-55 |
| ENSG00000111339 | ART4 | 2.29576458 | 4.87E-45 |
| ENSG00000156711 | MAPK13 | 2.293364315 | 2.99E-05 |
| ENSG00000260188 | gene_sourcehavana | 2.291559325 | 0.014163 |
| ENSG00000123358 | NR4A1 | 2.29140957 | 2.33E-28 |
| ENSG00000163683 | SMIM14 | 2.29111554 | 1.16E-05 |
| ENSG00000236209 | gene_sourcehavana | 2.290930818 | 0.01704 |
| ENSG00000255176 | gene_sourcehavana | 2.282006846 | 0.017138 |
| ENSG00000268119 | gene_sourcehavana | 2.276286486 | 3.27E-05 |
| ENSG00000006453 | BAIAP2L1 | 2.262602655 | 2.23E-19 |
| ENSG00000135636 | DYSF | 2.256234149 | 4.14E-102 |
| ENSG00000179059 | ZFP42 | 2.256061459 | 0.049091 |
| ENSG00000259033 | gene_sourcehavana_ | 2.255397836 | 3.40E-09 |

| | | | |
|---|---|---|---|
| ENSG00000167114 | SLC27A4 | 2.24569147 | 4.83E-122 |
| ENSG00000110693 | SOX6 | 2.244617162 | 5.48E-69 |
| ENSG00000105289 | TJP3 | 2.24385956 | 8.78E-06 |
| ENSG00000255200 | PGAM1P8 | 2.240678086 | 6.46E-07 |
| ENSG00000159388 | BTG2 | 2.239247889 | 5.17E-109 |
| ENSG00000273983 | H3C8 | 2.238384596 | 1.37E-07 |
| ENSG00000182732 | RGS6 | 2.237040922 | 1.90E-28 |
| ENSG00000103034 | NDRG4 | 2.236297066 | 4.44E-21 |
| ENSG00000158022 | TRIM63 | 2.235413517 | 0.004449 |
| ENSG00000134352 | IL6ST | 2.232965824 | 1.72E-33 |
| ENSG00000204396 | VWA7 | 2.224595242 | 3.50E-20 |
| ENSG00000288869 | SMANTIS | 2.2244076 | 2.91E-10 |
| ENSG00000136717 | BIN1 | 2.224301466 | 5.66E-20 |
| ENSG00000122035 | RASL11A | 2.22410074 | 1.39E-81 |
| ENSG00000232884 | gene_sourcehavana | 2.222485683 | 2.25E-21 |
| ENSG00000266947 | gene_sourcehavana | 2.222016635 | 0.003138 |
| ENSG00000223442 | TH2LCRR | 2.220635757 | 0.02365 |
| ENSG00000151012 | SLC7A11 | 2.220267545 | 9.62E-87 |
| ENSG00000156535 | CD109 | 2.220201616 | 1.10E-10 |
| ENSG00000198753 | PLXNB3 | 2.219826253 | 9.89E-80 |
| ENSG00000250318 | RPL13AP26 | 2.21981161 | 0.001598 |
| ENSG00000136026 | CKAP4 | 2.217318985 | 5.95E-07 |
| ENSG00000168056 | LTBP3 | 2.217219618 | 2.13E-44 |
| ENSG00000159314 | ARHGAP27 | 2.215921957 | 1.65E-06 |
| ENSG00000166444 | DENND2B | 2.213505026 | 0.001143 |
| ENSG00000229544 | NKX1-2 | 2.205948782 | 3.99E-20 |
| ENSG00000197122 | SRC | 2.20354618 | 1.14E-19 |
| ENSG00000279491 | gene_sourcehavana | 2.202039742 | 1.44E-05 |
| ENSG00000285646 | gene_sourcehavana | 2.196489481 | 0.004296 |
| ENSG00000168672 | LRATD2 | 2.18207235 | 4.64E-05 |
| ENSG00000281026 | N4BP2L2-IT2 | 2.179654554 | 4.53E-10 |
| ENSG00000165801 | ARHGEF40 | 2.178767774 | 9.38E-05 |
| ENSG00000182196 | ARL6IP4 | 2.177705647 | 0.008001 |
| ENSG00000119508 | NR4A3 | 2.177586451 | 0.000175 |
| ENSG00000183779 | ZNF703 | 2.174979158 | 2.50E-05 |
| ENSG00000196420 | S100A5 | 2.170354369 | 0.01858 |
| ENSG00000244491 | gene_sourcehavana | 2.170343255 | 0.038786 |
| ENSG00000118508 | RAB32 | 2.167499838 | 6.34E-21 |
| ENSG00000248671 | ALG1L9P | 2.167224789 | 0.031095 |
| ENSG00000249655 | gene_sourcehavana | 2.165107223 | 0.000428 |
| ENSG00000001617 | SEMA3F | 2.161187465 | 0.004167 |
| ENSG00000104856 | RELB | 2.156654159 | 1.78E-32 |
| ENSG00000004939 | SLC4A1 | 2.155654946 | 4.69E-149 |
| ENSG00000008086 | CDKL5 | 2.15518961 | 4.98E-11 |
| ENSG00000178082 | TWF1P1 | 2.154792368 | 0.047932 |
| ENSG00000087076 | HSD17B14 | 2.153163017 | 0.000236 |
| ENSG00000162772 | ATF3 | 2.150219402 | 4.81E-24 |
| ENSG00000214212 | C19orf38 | 2.149450977 | 0.018974 |
| ENSG00000166432 | ZMAT1 | 2.144953758 | 2.58E-18 |

| | | | |
|---|---|---|---|
| ENSG00000142149 | HUNK | 2.141313291 | 2.60E-53 |
| ENSG00000235527 | HIPK1-AS1 | 2.138645694 | 0.010769 |
| ENSG00000160408 | ST6GALNAC6 | 2.137180184 | 8.22E-10 |
| ENSG00000135931 | ARMC9 | 2.132593601 | 4.84E-18 |
| ENSG00000111344 | RASAL1 | 2.130446324 | 7.14E-09 |
| ENSG00000171246 | NPTX1 | 2.129980687 | 4.33E-87 |
| ENSG00000172738 | TMEM217 | 2.129951251 | 4.04E-16 |
| ENSG00000267221 | C17orf113 | 2.122033394 | 0.001215 |
| ENSG00000262155 | LINC02175 | 2.116641941 | 1.19E-10 |
| ENSG00000274750 | H3C6 | 2.115494416 | 1.67E-06 |
| ENSG00000184601 | C14orf180 | 2.109702154 | 0.010588 |
| ENSG00000257221 | gene_sourcehavana | 2.109346853 | 0.003589 |
| ENSG00000100055 | CYTH4 | 2.108004282 | 4.38E-11 |
| ENSG00000284138 | ATP6V0CP4 | 2.10745386 | 0.036056 |
| ENSG00000089692 | LAG3 | 2.100345401 | 0.025337 |
| ENSG00000208028 | MIR616 | 2.099656837 | 0.04371 |
| ENSG00000285294 | LINC00842 | 2.095706645 | 0.000113 |
| ENSG00000159640 | ACE | 2.087286557 | 0.043685 |
| ENSG00000113742 | CPEB4 | 2.085620911 | 2.11E-120 |
| ENSG00000130766 | SESN2 | 2.083330173 | 2.57E-49 |
| ENSG00000126353 | CCR7 | 2.08119178 | 7.71E-10 |
| ENSG00000178297 | TMPRSS9 | 2.080220711 | 0.018531 |
| ENSG00000163975 | MELTF | 2.076888729 | 9.93E-45 |
| ENSG00000250361 | GYPB | 2.074512911 | 2.30E-110 |
| ENSG00000145365 | TIFA | 2.074073106 | 0.022696 |
| ENSG00000183929 | DUSP5P1 | 2.071791179 | 1.93E-26 |
| ENSG00000122367 | LDB3 | 2.070555941 | 0.014403 |
| ENSG00000020577 | SAMD4A | 2.067355574 | 1.61E-50 |
| ENSG00000185267 | CDNF | 2.066960908 | 0.039688 |
| ENSG00000138646 | HERC5 | 2.06530411 | 1.93E-17 |
| ENSG00000159335 | PTMS | 2.063192195 | 2.01E-32 |
| ENSG00000106078 | COBL | 2.060524173 | 0.015716 |
| ENSG00000291008 | gene_sourcehavana | 2.054566708 | 0.000212 |
| ENSG00000196937 | FAM3C | 2.054382778 | 1.64E-103 |
| ENSG00000228436 | RRAGC-DT | 2.054102161 | 9.94E-06 |
| ENSG00000233818 | CLDN14-AS1 | 2.052665139 | 0.01708 |
| ENSG00000207561 | MIR635 | 2.046374521 | 0.039924 |
| ENSG00000162873 | KLHDC8A | 2.044391062 | 6.88E-102 |
| ENSG00000236320 | SLFN14 | 2.039768752 | 1.34E-105 |
| ENSG00000069667 | RORA | 2.039035883 | 9.79E-58 |
| ENSG00000112576 | CCND3 | 2.038717863 | 7.49E-137 |
| ENSG00000196878 | LAMB3 | 2.035562333 | 2.36E-09 |
| ENSG00000129355 | CDKN2D | 2.03502637 | 1.31E-33 |
| ENSG00000084734 | GCKR | 2.034741963 | 0.043372 |
| ENSG00000151914 | DST | 2.032360702 | 4.96E-98 |
| ENSG00000250075 | gene_sourcehavana | 2.031116505 | 0.048463 |
| ENSG00000160282 | FTCD | 2.028995538 | 2.73E-10 |
| ENSG00000180596 | H2BC4 | 2.028572291 | 4.78E-17 |
| ENSG00000131015 | ULBP2 | 2.02831475 | 0.001477 |

| | | | |
|---|---|---|---|
| ENSG00000078269 | SYNJ2 | 2.027206843 | 2.62E-67 |
| ENSG00000177191 | B3GNT8 | 2.020386495 | 1.30E-06 |
| ENSG00000105499 | PLA2G4C | 2.020217876 | 4.09E-07 |
| ENSG00000079308 | TNS1 | 2.016163457 | 2.98E-51 |
| ENSG00000170871 | KIAA0232 | 2.014924142 | 4.88E-150 |
| ENSG00000279900 | gene_sourcehavana | 2.013900976 | 0.009734 |
| ENSG00000171813 | PWWP2B | 2.011096662 | 0.001241 |
| ENSG00000168490 | PHYHIP | 2.009149113 | 6.74E-13 |
| ENSG00000136929 | HEMGN | 2.008945869 | 2.40E-198 |
| ENSG00000147883 | CDKN2B | 2.008101912 | 2.17E-42 |
| ENSG00000267632 | gene_sourcehavana | 2.007779932 | 8.78E-05 |
| ENSG00000249846 | LINC02021 | 2.005946596 | 0.020931 |
| ENSG00000270084 | GAS5-AS1 | 2.005108764 | 0.034503 |
| ENSG00000205726 | ITSN1 | 2.005082362 | 3.96E-118 |
| ENSG00000104321 | TRPA1 | 2.001619456 | 0.01116 |
| ENSG00000182372 | CLN8 | 2.001467329 | 9.50E-14 |
| ENSG00000124006 | OBSL1 | 1.993847042 | 0.01083 |
| ENSG00000224356 | gene_sourcehavana | 1.990898049 | 0.023187 |
| ENSG00000198580 | gene_sourcehavana | 1.989886828 | 0.017316 |
| ENSG00000287094 | gene_sourcehavana_ | 1.988371736 | 1.98E-05 |
| ENSG00000223552 | CCR5AS | 1.987103623 | 0.001839 |
| ENSG00000109321 | AREG | 1.980417634 | 4.60E-09 |
| ENSG00000221963 | APOL6 | 1.976567245 | 9.71E-26 |
| ENSG00000182809 | CRIP2 | 1.975693822 | 6.63E-12 |
| ENSG00000266907 | gene_sourcehavana | 1.975383105 | 8.93E-07 |
| ENSG00000269921 | gene_sourcehavana | 1.969506707 | 0.00169 |
| ENSG00000285933 | gene_sourcehavana | 1.96533288 | 3.58E-54 |
| ENSG00000114739 | ACVR2B | 1.964390609 | 1.04E-45 |
| ENSG00000131398 | KCNC3 | 1.963665988 | 4.36E-05 |
| ENSG00000164078 | MST1R | 1.958888749 | 0.011145 |
| ENSG00000141540 | TTYH2 | 1.957318465 | 2.17E-57 |
| ENSG00000234684 | SDCBP2-AS1 | 1.95727633 | 0.000203 |
| ENSG00000109689 | STIM2 | 1.956267494 | 4.13E-62 |
| ENSG00000140961 | OSGIN1 | 1.955897663 | 0.000114 |
| ENSG00000169896 | ITGAM | 1.953484377 | 0.02196 |
| ENSG00000051108 | HERPUD1 | 1.951171389 | 5.34E-107 |
| ENSG00000197535 | MYO5A | 1.950521872 | 1.43E-06 |
| ENSG00000170190 | SLC16A5 | 1.948988599 | 0.004845 |
| ENSG00000156966 | B3GNT7 | 1.946057076 | 0.000222 |
| ENSG00000290652 | gene_sourcehavana | 1.944486709 | 1.12E-49 |
| ENSG00000278828 | H3C10 | 1.944022717 | 3.10E-13 |
| ENSG00000223609 | HBD | 1.943107043 | 3.04E-220 |
| ENSG00000206878 | gene_sourceensembl | 1.937452092 | 5.11E-07 |
| ENSG00000225828 | FAM229A | 1.937379381 | 5.31E-20 |
| ENSG00000204580 | DDR1 | 1.935795295 | 1.51E-26 |
| ENSG00000248734 | gene_sourcehavana | 1.933708303 | 0.032195 |
| ENSG00000172183 | ISG20 | 1.932356051 | 2.30E-15 |
| ENSG00000253669 | GASAL1 | 1.931230117 | 0.000747 |
| ENSG00000262140 | gene_sourcehavana | 1.930105379 | 0.01089 |

| | | | |
|---|---|---|---|
| ENSG00000163106 | HPGDS | 1.928325367 | 9.45E-46 |
| ENSG00000170385 | SLC30A1 | 1.922468285 | 1.90E-63 |
| ENSG00000185015 | CA13 | 1.912233481 | 0.028851 |
| ENSG00000143878 | RHOB | 1.91012329 | 5.11E-58 |
| ENSG00000260563 | gene_sourcehavana | 1.90962199 | 1.87E-08 |
| ENSG00000286448 | gene_sourcehavana_ | 1.908507925 | 0.000211 |
| ENSG00000275318 | gene_sourcehavana | 1.904569783 | 0.013053 |
| ENSG00000165914 | TTC7B | 1.904437629 | 8.60E-07 |
| ENSG00000095794 | CREM | 1.898744758 | 1.93E-36 |
| ENSG00000124508 | BTN2A2 | 1.8947724 | 6.75E-47 |
| ENSG00000290792 | gene_sourcehavana | 1.894338656 | 4.08E-23 |
| ENSG00000185909 | KLHDC8B | 1.893571893 | 3.60E-117 |
| ENSG00000274020 | LINC01138 | 1.893149419 | 0.000532 |
| ENSG00000123240 | OPTN | 1.890632067 | 6.41E-97 |
| ENSG00000262877 | gene_sourcehavana | 1.888827372 | 0.008848 |
| ENSG00000279744 | gene_sourcehavana | 1.886971771 | 7.97E-17 |
| ENSG00000146242 | TPBG | 1.884662124 | 0.000834 |
| ENSG00000273402 | gene_sourcehavana | 1.884585682 | 0.029993 |
| ENSG00000166145 | SPINT1 | 1.884349713 | 4.44E-26 |
| ENSG00000187474 | FPR3 | 1.884126906 | 0.000763 |
| ENSG00000051128 | HOMER3 | 1.883374958 | 0.003657 |
| ENSG00000184384 | MAML2 | 1.882921751 | 6.17E-08 |
| ENSG00000147082 | CCNB3 | 1.879860001 | 1.24E-07 |
| ENSG00000115295 | CLIP4 | 1.878762099 | 1.21E-05 |
| ENSG00000189143 | CLDN4 | 1.877486279 | 0.016499 |
| ENSG00000124570 | SERPINB6 | 1.876142166 | 6.05E-13 |
| ENSG00000136404 | TM6SF1 | 1.875039016 | 3.73E-30 |
| ENSG00000177963 | RIC8A | 1.874581433 | 6.09E-100 |
| ENSG00000183688 | RFLNB | 1.874427033 | 6.95E-66 |
| ENSG00000155749 | FLACC1 | 1.873791253 | 7.08E-06 |
| ENSG00000108840 | HDAC5 | 1.872441919 | 4.61E-94 |
| ENSG00000224892 | RPS4XP16 | 1.872125636 | 0.00896 |
| ENSG00000205213 | LGR4 | 1.867478844 | 9.10E-58 |
| ENSG00000181192 | DHTKD1 | 1.866123469 | 3.69E-103 |
| ENSG00000266066 | POLRMTP1 | 1.862757725 | 6.64E-23 |
| ENSG00000169946 | ZFPM2 | 1.862007895 | 5.72E-09 |
| ENSG00000207005 | RNU1-2 | 1.861445216 | 0.048272 |
| ENSG00000087589 | CASS4 | 1.857074441 | 2.35E-22 |
| ENSG00000128564 | VGF | 1.84676096 | 7.74E-10 |
| ENSG00000146112 | PPP1R18 | 1.842903128 | 3.14E-65 |
| ENSG00000136235 | GPNMB | 1.839571755 | 9.29E-12 |
| ENSG00000263766 | KPNB1-DT | 1.839308789 | 0.008939 |
| ENSG00000164733 | CTSB | 1.838920717 | 8.50E-79 |
| ENSG00000271643 | PDCD6IP-DT | 1.836127305 | 2.91E-12 |
| ENSG00000262410 | gene_sourcehavana | 1.834820301 | 0.000147 |
| ENSG00000102032 | RENBP | 1.83462647 | 8.08E-46 |
| ENSG00000266777 | SH3GL1P1 | 1.833689669 | 0.000103 |
| ENSG00000120306 | CYSTM1 | 1.828134399 | 2.37E-70 |
| ENSG00000225964 | NRIR | 1.826654687 | 9.25E-09 |

| | | | |
|---|---|---|---|
| ENSG00000143198 | MGST3 | 1.826336429 | 1.18E-105 |
| ENSG00000166971 | AKTIP | 1.825628239 | 6.51E-10 |
| ENSG00000127184 | COX7C | 1.822225356 | 3.87E-119 |
| ENSG00000184226 | PCDH9 | 1.818438237 | 3.22E-11 |
| ENSG00000224614 | TNK2-AS1 | 1.818030141 | 6.09E-05 |
| ENSG00000091490 | SEL1L3 | 1.812031431 | 5.51E-60 |
| ENSG00000289126 | gene_sourcehavana_ | 1.811187381 | 0.007566 |
| ENSG00000170180 | GYPA | 1.80847695 | 7.32E-147 |
| ENSG00000145335 | SNCA | 1.808332191 | 1.17E-35 |
| ENSG00000277526 | LINC03021 | 1.807307524 | 0.015517 |
| ENSG00000266976 | gene_sourcehavana | 1.806706715 | 5.29E-14 |
| ENSG00000143515 | ATP8B2 | 1.800872463 | 9.34E-72 |
| ENSG00000278921 | EPB41L4A-DT | 1.798387964 | 0.001795 |
| ENSG00000271425 | NBPF10 | 1.798280691 | 3.05E-10 |
| ENSG00000268895 | A1BG-AS1 | 1.796753575 | 0.041761 |
| ENSG00000235663 | SAPCD1-AS1 | 1.795740133 | 0.001217 |
| ENSG00000274210 | RNVU1-27 | 1.792907136 | 0.018179 |
| ENSG00000197405 | C5AR1 | 1.792812012 | 0.007542 |
| ENSG00000165410 | CFL2 | 1.791899007 | 1.03E-39 |
| ENSG00000189060 | H1-0 | 1.790919564 | 3.28E-28 |
| ENSG00000279700 | gene_sourcehavana | 1.788067385 | 0.003863 |
| ENSG00000198478 | SH3BGRL2 | 1.785432348 | 0.006687 |
| ENSG00000248643 | RBM14-RBM4 | 1.783823784 | 0.019951 |
| ENSG00000196421 | C20orf204 | 1.782877907 | 0.001524 |
| ENSG00000114270 | COL7A1 | 1.776106503 | 0.022765 |
| ENSG00000275302 | CCL4 | 1.775759394 | 0.006457 |
| ENSG00000269600 | gene_sourcehavana | 1.775567746 | 1.27E-55 |
| ENSG00000280303 | ERICD | 1.772572478 | 2.82E-07 |
| ENSG00000072682 | P4HA2 | 1.772107105 | 0.007808 |
| ENSG00000068366 | ACSL4 | 1.771878675 | 1.47E-83 |
| ENSG00000213090 | SCYL2P1 | 1.769352369 | 0.030825 |
| ENSG00000088826 | SMOX | 1.764492912 | 8.47E-24 |
| ENSG00000135048 | CEMIP2 | 1.764279955 | 2.90E-120 |
| ENSG00000241860 | gene_sourcehavana | 1.759102677 | 0.036035 |
| ENSG00000107201 | RIGI | 1.756930299 | 0.037355 |
| ENSG00000142949 | PTPRF | 1.756875709 | 6.62E-17 |
| ENSG00000099625 | CBARP | 1.755949668 | 6.57E-10 |
| ENSG00000114166 | KAT2B | 1.755820394 | 3.86E-85 |
| ENSG00000115993 | TRAK2 | 1.752982093 | 8.31E-130 |
| ENSG00000238269 | PAGE2B | 1.748653008 | 0.003185 |
| ENSG00000236377 | gene_sourcehavana | 1.748336077 | 0.031713 |
| ENSG00000159346 | ADIPOR1 | 1.748105733 | 2.55E-144 |
| ENSG00000116016 | EPAS1 | 1.746673452 | 6.41E-63 |
| ENSG00000160191 | PDE9A | 1.745763016 | 0.006135 |
| ENSG00000143595 | AQP10 | 1.744523997 | 4.15E-31 |
| ENSG00000128965 | CHAC1 | 1.742404682 | 5.16E-26 |
| ENSG00000027869 | SH2D2A | 1.742294846 | 3.42E-30 |
| ENSG00000131370 | SH3BP5 | 1.741646195 | 0.009733 |
| ENSG00000095383 | TBC1D2 | 1.741451427 | 4.22E-17 |

| | | | |
|---|---|---|---|
| ENSG00000131400 | NAPSA | 1.735839193 | 0.045653 |
| ENSG00000150637 | CD226 | 1.734406414 | 0.006574 |
| ENSG00000102547 | CAB39L | 1.732763597 | 6.81E-26 |
| ENSG00000147642 | SYBU | 1.732242803 | 3.55E-08 |
| ENSG00000184613 | NELL2 | 1.731946415 | 0.002627 |
| ENSG00000107796 | ACTA2 | 1.731814963 | 0.015236 |
| ENSG00000163251 | FZD5 | 1.731678038 | 2.57E-69 |
| ENSG00000272375 | gene_sourcehavana | 1.727845732 | 0.002668 |
| ENSG00000271795 | gene_sourcehavana | 1.725722983 | 1.77E-05 |
| ENSG00000110002 | VWA5A | 1.72410001 | 1.84E-188 |
| ENSG00000278356 | gene_sourcehavana | 1.723054232 | 0.020615 |
| ENSG00000047644 | WWC3 | 1.720124732 | 2.60E-66 |
| ENSG00000196517 | SLC6A9 | 1.719648382 | 2.22E-79 |
| ENSG00000259498 | TPM1-AS | 1.718917634 | 1.53E-10 |
| ENSG00000160951 | PTGER1 | 1.718186874 | 0.044446 |
| ENSG00000163219 | ARHGAP25 | 1.710504122 | 2.76E-31 |
| ENSG00000137843 | PAK6 | 1.70965127 | 5.24E-07 |
| ENSG00000184619 | KRBA2 | 1.707849368 | 8.77E-08 |
| ENSG00000165259 | HDX | 1.707616052 | 2.72E-05 |
| ENSG00000175040 | CHST2 | 1.7054505 | 1.83E-72 |
| ENSG00000170955 | CAVIN3 | 1.703277544 | 0.002202 |
| ENSG00000257524 | gene_sourcehavana | 1.701240055 | 0.013192 |
| ENSG00000105464 | GRIN2D | 1.701112509 | 7.87E-09 |
| ENSG00000257594 | GALNT4 | 1.700939478 | 0.02086 |
| ENSG00000275131 | PDE4DIPP2 | 1.696335948 | 0.003247 |
| ENSG00000213722 | DDAH2 | 1.694921675 | 7.98E-23 |
| ENSG00000151690 | MFSD6 | 1.694401828 | 6.79E-44 |
| ENSG00000204323 | SMIM5 | 1.69348515 | 2.16E-83 |
| ENSG00000273456 | gene_sourcehavana | 1.693263806 | 0.036174 |
| ENSG00000100249 | C22orf31 | 1.689472861 | 0.016519 |
| ENSG00000137936 | BCAR3 | 1.686649939 | 1.23E-09 |
| ENSG00000057468 | MSH4 | 1.682956431 | 0.044681 |
| ENSG00000184557 | SOCS3 | 1.682166035 | 0.000333 |
| ENSG00000279861 | gene_sourcehavana | 1.681449124 | 3.87E-11 |
| ENSG00000162390 | ACOT11 | 1.679784612 | 0.000282 |
| ENSG00000120899 | PTK2B | 1.67434316 | 7.26E-106 |
| ENSG00000289341 | gene_sourcehavana_ | 1.673064339 | 5.86E-06 |
| ENSG00000184378 | ACTRT3 | 1.672686318 | 1.01E-06 |
| ENSG00000168071 | CCDC88B | 1.671803227 | 2.28E-26 |
| ENSG00000182310 | SPACA6 | 1.670809549 | 8.32E-08 |
| ENSG00000102057 | KCND1 | 1.670333408 | 0.000193 |
| ENSG00000114279 | FGF12 | 1.668174573 | 0.003601 |
| ENSG00000274180 | NATD1 | 1.667384481 | 1.43E-95 |
| ENSG00000173542 | MOB1B | 1.666549745 | 1.34E-89 |
| ENSG00000178104 | PDE4DIP | 1.665422761 | 1.06E-40 |
| ENSG00000112195 | TREML2 | 1.663811573 | 1.54E-14 |
| ENSG00000069974 | RAB27A | 1.662756922 | 3.46E-63 |
| ENSG00000114841 | DNAH1 | 1.661069882 | 2.01E-06 |
| ENSG00000258748 | KIF26A-DT | 1.660195358 | 0.000667 |

| | | | |
|---|---|---|---|
| ENSG00000095637 | SORBS1 | 1.659567479 | 1.54E-58 |
| ENSG00000276410 | H2BC3 | 1.659156381 | 0.001557 |
| ENSG00000243107 | gene_sourcehavana | 1.659149816 | 0.034042 |
| ENSG00000278376 | gene_sourcehavana | 1.658395765 | 0.021796 |
| ENSG00000276478 | gene_sourcehavana | 1.657786764 | 4.18E-06 |
| ENSG00000160325 | CACFD1 | 1.656811047 | 5.83E-05 |
| ENSG00000141934 | PLPP2 | 1.655975314 | 2.65E-10 |
| ENSG00000204335 | SP5 | 1.655940556 | 2.93E-05 |
| ENSG00000125089 | SH3TC1 | 1.653721541 | 0.010623 |
| ENSG00000101844 | ATG4A | 1.651604273 | 5.74E-76 |
| ENSG00000166046 | TCP11L2 | 1.651101292 | 3.27E-23 |
| ENSG00000234156 | gene_sourcehavana | 1.649984074 | 3.79E-11 |
| ENSG00000198711 | SSBP3-AS1 | 1.64679126 | 0.019545 |
| ENSG00000256043 | CTSO | 1.646353708 | 1.60E-08 |
| ENSG00000168016 | TRANK1 | 1.646193652 | 2.35E-46 |
| ENSG00000141655 | TNFRSF11A | 1.645033996 | 5.24E-23 |
| ENSG00000197461 | PDGFA | 1.644483782 | 0.000445 |
| ENSG00000256973 | gene_sourcehavana | 1.643058829 | 0.049989 |
| ENSG00000175197 | DDIT3 | 1.641623254 | 9.65E-17 |
| ENSG00000255326 | gene_sourcehavana | 1.641536245 | 0.006487 |
| ENSG00000165891 | E2F7 | 1.641025171 | 4.58E-63 |
| ENSG00000110911 | SLC11A2 | 1.63659773 | 7.29E-74 |
| ENSG00000042062 | RIPOR3 | 1.634153177 | 1.84E-51 |
| ENSG00000100036 | SLC35E4 | 1.633701865 | 2.04E-06 |
| ENSG00000267189 | ATP5MGP7 | 1.633275654 | 0.013801 |
| ENSG00000102554 | KLF5 | 1.631379626 | 5.12E-10 |
| ENSG00000183723 | CMTM4 | 1.631045936 | 6.44E-28 |
| ENSG00000235128 | gene_sourcehavana | 1.629368622 | 7.02E-07 |
| ENSG00000125895 | TMEM74B | 1.628763295 | 7.39E-17 |
| ENSG00000003249 | DBNDD1 | 1.628694984 | 0.034383 |
| ENSG00000186834 | HEXIM1 | 1.627746578 | 4.54E-79 |
| ENSG00000120217 | CD274 | 1.627516374 | 8.71E-23 |
| ENSG00000130307 | USHBP1 | 1.626600206 | 0.038081 |
| ENSG00000104327 | CALB1 | 1.625796899 | 1.30E-05 |
| ENSG00000198452 | OR14L1P | 1.622776105 | 0.013741 |
| ENSG00000166343 | MSS51 | 1.622503423 | 0.000939 |
| ENSG00000172086 | KRCC1 | 1.621919581 | 1.61E-34 |
| ENSG00000139438 | FAM222A | 1.619739451 | 2.05E-13 |
| ENSG00000130005 | GAMT | 1.612153978 | 8.48E-14 |
| ENSG00000178719 | GRINA | 1.611124128 | 1.93E-83 |
| ENSG00000235652 | EPM2A-DT | 1.608506522 | 0.002421 |
| ENSG00000168386 | FILIP1L | 1.608503844 | 9.67E-09 |
| ENSG00000137942 | FNBP1L | 1.607365983 | 1.11E-37 |
| ENSG00000213626 | LBH | 1.604369608 | 4.41E-71 |
| ENSG00000273820 | USP27X | 1.603488478 | 3.17E-11 |
| ENSG00000137880 | GCHFR | 1.599745145 | 1.10E-80 |
| ENSG00000170542 | SERPINB9 | 1.59707469 | 3.99E-09 |
| ENSG00000185022 | MAFF | 1.595363249 | 1.58E-17 |
| ENSG00000181218 | H2AC25 | 1.594279205 | 6.49E-15 |

| | | | |
|---|---|---|---|
| ENSG00000260852 | FBXL19-AS1 | 1.592658505 | 9.43E-30 |
| ENSG00000197582 | GPX1P1 | 1.591652755 | 6.02E-31 |
| ENSG00000049323 | LTBP1 | 1.591247966 | 1.10E-40 |
| ENSG00000206418 | RAB12 | 1.589163038 | 4.00E-36 |
| ENSG00000149218 | ENDOD1 | 1.588459911 | 2.39E-43 |
| ENSG00000119986 | AVPI1 | 1.587748924 | 1.19E-42 |
| ENSG00000124145 | SDC4 | 1.587097677 | 1.43E-72 |
| ENSG00000281530 | gene_sourcehavana | 1.585091909 | 0.033738 |
| ENSG00000196792 | STRN3 | 1.584542365 | 4.91E-63 |
| ENSG00000280033 | gene_sourcehavana | 1.584186183 | 0.044615 |
| ENSG00000135677 | GNS | 1.583982588 | 1.79E-73 |
| ENSG00000144843 | ADPRH | 1.583300693 | 2.80E-11 |
| ENSG00000203666 | EFCAB2 | 1.576144791 | 0.020472 |
| ENSG00000068971 | PPP2R5B | 1.575031249 | 8.69E-64 |
| ENSG00000167748 | KLK1 | 1.574598192 | 1.23E-32 |
| ENSG00000111229 | ARPC3 | 1.574305362 | 2.06E-88 |
| ENSG00000101670 | LIPG | 1.570371165 | 0.002782 |
| ENSG00000171724 | VAT1L | 1.568402704 | 0.000303 |
| ENSG00000137460 | FHDC1 | 1.566792769 | 3.84E-68 |
| ENSG00000289840 | gene_sourcehavana_ | 1.566360552 | 2.37E-09 |
| ENSG00000280069 | gene_sourcehavana | 1.566174876 | 0.013342 |
| ENSG00000185222 | TCEAL9 | 1.565873359 | 3.75E-12 |
| ENSG00000180667 | YOD1 | 1.565829738 | 4.54E-56 |
| ENSG00000175305 | CCNE2 | 1.565783715 | 2.51E-96 |
| ENSG00000213347 | MXD3 | 1.564613235 | 9.61E-44 |
| ENSG00000128383 | APOBEC3A | 1.561267468 | 0.035925 |
| ENSG00000134321 | RSAD2 | 1.561168565 | 2.78E-53 |
| ENSG00000196369 | SRGAP2B | 1.558423038 | 4.29E-12 |
| ENSG00000011198 | ABHD5 | 1.558232692 | 2.01E-73 |
| ENSG00000065717 | TLE2 | 1.556927212 | 0.025534 |
| ENSG00000259986 | gene_sourcehavana | 1.555166987 | 7.70E-19 |
| ENSG00000167107 | ACSF2 | 1.553777512 | 1.11E-19 |
| ENSG00000148180 | GSN | 1.553368735 | 8.23E-25 |
| ENSG00000077150 | NFKB2 | 1.552503765 | 1.31E-22 |
| ENSG00000164463 | CREBRF | 1.549857516 | 2.84E-47 |
| ENSG00000250334 | LINC00989 | 1.549085892 | 0.006975 |
| ENSG00000156671 | SAMD8 | 1.54906131 | 1.06E-28 |
| ENSG00000157107 | FCHO2 | 1.547987379 | 6.89E-83 |
| ENSG00000087085 | ACHE | 1.547082731 | 4.82E-58 |
| ENSG00000187688 | TRPV2 | 1.54626127 | 2.88E-60 |
| ENSG00000185862 | EVI2B | 1.546019891 | 9.69E-07 |
| ENSG00000142082 | SIRT3 | 1.544517275 | 2.43E-38 |
| ENSG00000116857 | TMEM9 | 1.540917774 | 3.02E-56 |
| ENSG00000084207 | GSTP1 | 1.538827845 | 4.02E-13 |
| ENSG00000144331 | ZNF385B | 1.53242117 | 1.51E-06 |
| ENSG00000284606 | gene_sourcehavana | 1.532194792 | 1.54E-06 |
| ENSG00000151689 | INPP1 | 1.529835371 | 1.46E-18 |
| ENSG00000099994 | SUSD2 | 1.528542477 | 0.018246 |
| ENSG00000143367 | TUFT1 | 1.526278944 | 2.31E-25 |

| | | | |
|---|---|---|---|
| ENSG00000131979 | GCH1 | 1.526197817 | 2.46E-29 |
| ENSG00000141279 | NPEPPS | 1.523308979 | 1.65E-74 |
| ENSG00000064687 | ABCA7 | 1.522980211 | 3.34E-64 |
| ENSG00000286677 | gene_sourcehavana_ | 1.521393166 | 0.004742 |
| ENSG00000115657 | ABCB6 | 1.521181866 | 3.51E-08 |
| ENSG00000197885 | NKIRAS1 | 1.518177681 | 2.52E-10 |
| ENSG00000004799 | PDK4 | 1.515265148 | 0.000132 |
| ENSG00000107738 | VSIR | 1.514991153 | 1.93E-10 |
| ENSG00000233080 | LINC01399 | 1.513930237 | 2.91E-14 |
| ENSG00000100351 | GRAP2 | 1.508260131 | 7.03E-45 |
| ENSG00000122862 | SRGN | 1.507897594 | 1.09E-45 |
| ENSG00000104863 | LIN7B | 1.507737784 | 8.27E-27 |
| ENSG00000124635 | H2BC11 | 1.504655496 | 2.87E-11 |
| ENSG00000111644 | ACRBP | 1.502717486 | 0.032949 |
| ENSG00000111913 | RIPOR2 | 1.499076067 | 0.036419 |
| ENSG00000138642 | HERC6 | 1.498955363 | 7.86E-52 |
| ENSG00000106125 | MINDY4 | 1.498630224 | 0.011808 |
| ENSG00000273270 | LINC03072 | 1.49815235 | 6.82E-06 |
| ENSG00000143842 | SOX13 | 1.495889883 | 0.008707 |
| ENSG00000070669 | ASNS | 1.495619487 | 2.46E-08 |
| ENSG00000165886 | UBTD1 | 1.492189165 | 7.32E-05 |
| ENSG00000143630 | HCN3 | 1.491089815 | 9.49E-14 |
| ENSG00000101000 | PROCR | 1.490983352 | 1.12E-09 |
| ENSG00000121797 | CCRL2 | 1.490475326 | 1.24E-32 |
| ENSG00000078804 | TP53INP2 | 1.490262447 | 9.38E-29 |
| ENSG00000121104 | FAM117A | 1.487958166 | 1.82E-164 |
| ENSG00000182628 | SKA2 | 1.487566366 | 7.79E-38 |
| ENSG00000165168 | CYBB | 1.486191492 | 3.31E-39 |
| ENSG00000127831 | VIL1 | 1.484133604 | 7.49E-05 |
| ENSG00000070159 | PTPN3 | 1.48302054 | 0.00357 |
| ENSG00000128590 | DNAJB9 | 1.481248674 | 6.77E-30 |
| ENSG00000285077 | ARHGAP11B | 1.480659626 | 0.000143 |
| ENSG00000183722 | LHFPL6 | 1.479152918 | 9.53E-09 |
| ENSG00000282988 | gene_sourcehavana | 1.478897857 | 0.002857 |
| ENSG00000109255 | NMU | 1.474061146 | 3.57E-05 |
| ENSG00000285043 | gene_sourceensembl | 1.472707998 | 0.000301 |
| ENSG00000171703 | TCEA2 | 1.471430766 | 4.91E-10 |
| ENSG00000151239 | TWF1 | 1.468302285 | 3.37E-47 |
| ENSG00000285796 | gene_sourcehavana | 1.464663491 | 3.87E-05 |
| ENSG00000019485 | PRDM11 | 1.463432365 | 7.30E-45 |
| ENSG00000023909 | GCLM | 1.463052437 | 1.14E-97 |
| ENSG00000136854 | STXBP1 | 1.459761802 | 5.96E-06 |
| ENSG00000165434 | PGM2L1 | 1.458837577 | 1.51E-42 |
| ENSG00000151135 | TMEM263 | 1.458276728 | 4.26E-45 |
| ENSG00000213445 | SIPA1 | 1.458176407 | 4.73E-56 |
| ENSG00000063180 | CA11 | 1.457654915 | 1.09E-26 |
| ENSG00000030582 | GRN | 1.45705383 | 7.80E-83 |
| ENSG00000172748 | ZNF596 | 1.456074946 | 4.14E-16 |
| ENSG00000152292 | SH2D6 | 1.454458669 | 0.047602 |

| | | | |
|---|---|---|---|
| ENSG00000272405 | gene_sourcehavana | 1.452378026 | 0.012243 |
| ENSG00000137496 | IL18BP | 1.451749954 | 3.66E-35 |
| ENSG00000213977 | TAX1BP3 | 1.451055347 | 0.024702 |
| ENSG00000131069 | ACSS2 | 1.446847999 | 1.32E-92 |
| ENSG00000278949 | gene_sourcehavana | 1.44520707 | 0.014121 |
| ENSG00000224982 | TMEM233 | 1.445065045 | 1.28E-06 |
| ENSG00000133131 | MORC4 | 1.444170678 | 1.82E-07 |
| ENSG00000147650 | LRP12 | 1.443972975 | 1.10E-15 |
| ENSG00000064547 | LPAR2 | 1.442976452 | 0.004247 |
| ENSG00000007968 | E2F2 | 1.44252487 | 5.64E-83 |
| ENSG00000136840 | ST6GALNAC4 | 1.441390468 | 2.44E-69 |
| ENSG00000141504 | SAT2 | 1.440689576 | 3.95E-07 |
| ENSG00000150337 | FCGR1A | 1.440272874 | 7.10E-53 |
| ENSG00000135409 | AMHR2 | 1.43550704 | 2.17E-20 |
| ENSG00000130176 | CNN1 | 1.432010364 | 0.008352 |
| ENSG00000163218 | PGLYRP4 | 1.42998783 | 0.003191 |
| ENSG00000288935 | gene_sourcehavana_ | 1.429751211 | 0.039481 |
| ENSG00000103269 | RHBDL1 | 1.425247698 | 0.001967 |
| ENSG00000163827 | LRRC2 | 1.425058106 | 5.23E-12 |
| ENSG00000213918 | DNASE1 | 1.424728205 | 1.03E-23 |
| ENSG00000076067 | RBMS2 | 1.424446625 | 5.06E-07 |
| ENSG00000282508 | LINC01002 | 1.423447385 | 0.003663 |
| ENSG00000152556 | PFKM | 1.422845511 | 5.40E-47 |
| ENSG00000182795 | C1orf116 | 1.422475376 | 6.58E-08 |
| ENSG00000080819 | CPOX | 1.422020824 | 1.85E-99 |
| ENSG00000271614 | ATP2B1-AS1 | 1.421861759 | 0.039785 |
| ENSG00000137831 | UACA | 1.421373376 | 8.76E-06 |
| ENSG00000291194 | gene_sourcehavana | 1.420938209 | 4.18E-07 |
| ENSG00000286786 | gene_sourcehavana_ | 1.420765627 | 0.028951 |
| ENSG00000106976 | DNM1 | 1.420624662 | 5.63E-05 |
| ENSG00000257315 | ZBED6 | 1.419895399 | 0.000116 |
| ENSG00000276791 | gene_sourcehavana | 1.419836013 | 0.049637 |
| ENSG00000160058 | BSDC1 | 1.418590053 | 5.10E-75 |
| ENSG00000158966 | CACHD1 | 1.417512541 | 4.03E-05 |
| ENSG00000158856 | DMTN | 1.415940153 | 3.85E-94 |
| ENSG00000139626 | ITGB7 | 1.415153599 | 2.23E-33 |
| ENSG00000110047 | EHD1 | 1.413906263 | 1.08E-45 |
| ENSG00000139793 | MBNL2 | 1.413899334 | 1.51E-50 |
| ENSG00000148498 | PARD3 | 1.413692187 | 3.08E-35 |
| ENSG00000076944 | STXBP2 | 1.411600349 | 1.23E-15 |
| ENSG00000049249 | TNFRSF9 | 1.411475458 | 0.035826 |
| ENSG00000134955 | SLC37A2 | 1.410968996 | 1.18E-39 |
| ENSG00000105327 | BBC3 | 1.408059395 | 2.90E-31 |
| ENSG00000198932 | GPRASP1 | 1.407252979 | 0.01052 |
| ENSG00000167220 | HDHD2 | 1.407082905 | 2.64E-27 |
| ENSG00000066735 | KIF26A | 1.406019344 | 3.14E-84 |
| ENSG00000274272 | gene_sourcehavana | 1.404313819 | 0.001069 |
| ENSG00000180089 | TMEM86B | 1.403139981 | 3.72E-20 |
| ENSG00000261840 | gene_sourceensembl | 1.402887161 | 0.044452 |

| | | | |
|---|---|---|---|
| ENSG00000137941 | TTLL7 | 1.402833809 | 0.006818 |
| ENSG00000205502 | C2CD4B | 1.402587878 | 0.041387 |
| ENSG00000081181 | ARG2 | 1.401063845 | 4.78E-29 |
| ENSG00000137198 | GMPR | 1.400660019 | 8.69E-95 |
| ENSG00000261499 | gene_sourcehavana | 1.398744749 | 0.041687 |
| ENSG00000196741 | LINC01560 | 1.397820427 | 0.000424 |
| ENSG00000102024 | PLS3 | 1.396633801 | 5.79E-34 |
| ENSG00000151893 | CACUL1 | 1.39377873 | 2.50E-53 |
| ENSG00000108960 | MMD | 1.393395837 | 4.08E-22 |
| ENSG00000104883 | PEX11G | 1.392723672 | 0.002434 |
| ENSG00000283674 | gene_sourcehavana | 1.392141718 | 1.51E-15 |
| ENSG00000154898 | CCDC144CP | 1.389050794 | 0.00013 |
| ENSG00000269889 | gene_sourcehavana | 1.388721661 | 7.84E-05 |
| ENSG00000172432 | GTPBP2 | 1.388004514 | 7.54E-73 |
| ENSG00000177303 | CASKIN2 | 1.387527383 | 9.40E-10 |
| ENSG00000136044 | APPL2 | 1.387310147 | 2.26E-36 |
| ENSG00000145012 | LPP | 1.384520148 | 1.06E-21 |
| ENSG00000141574 | SECTM1 | 1.38398674 | 8.99E-05 |
| ENSG00000177692 | DNAJC28 | 1.382687054 | 3.72E-12 |
| ENSG00000205913 | SRRM2-AS1 | 1.382056202 | 0.000219 |
| ENSG00000237441 | RGL2 | 1.381280474 | 2.11E-32 |
| ENSG00000165169 | DYNLT3 | 1.381023521 | 2.98E-25 |
| ENSG00000156011 | PSD3 | 1.379654461 | 0.004793 |
| ENSG00000013364 | MVP | 1.379413369 | 6.39E-25 |
| ENSG00000242498 | ARPIN | 1.377758871 | 0.001609 |
| ENSG00000127946 | HIP1 | 1.376918545 | 0.017278 |
| ENSG00000197694 | SPTAN1 | 1.374351016 | 2.69E-28 |
| ENSG00000244119 | PDCL3P4 | 1.37418123 | 0.004387 |
| ENSG00000116717 | GADD45A | 1.373568433 | 1.12E-28 |
| ENSG00000166405 | RIC3 | 1.372923224 | 3.23E-10 |
| ENSG00000133639 | BTG1 | 1.36835319 | 5.66E-38 |
| ENSG00000260592 | gene_sourcehavana | 1.367899096 | 9.44E-17 |
| ENSG00000275719 | gene_sourceensembl | 1.367863577 | 0.000608 |
| ENSG00000145979 | TBC1D7 | 1.367421082 | 2.22E-43 |
| ENSG00000132471 | WBP2 | 1.367356176 | 6.53E-66 |
| ENSG00000103160 | HSDL1 | 1.366706625 | 2.57E-31 |
| ENSG00000268006 | PTOV1-AS1 | 1.365870621 | 0.016249 |
| ENSG00000006756 | ARSD | 1.364479607 | 1.51E-10 |
| ENSG00000253570 | RNF5P1 | 1.362119013 | 0.042401 |
| ENSG00000111266 | DUSP16 | 1.361990193 | 1.74E-11 |
| ENSG00000167103 | PIP5KL1 | 1.361711671 | 0.00935 |
| ENSG00000125967 | NECAB3 | 1.360674998 | 4.44E-32 |
| ENSG00000224040 | HMGN1P4 | 1.360469458 | 0.007769 |
| ENSG00000285103 | gene_sourcehavana | 1.359002853 | 0.000411 |
| ENSG00000138166 | DUSP5 | 1.358900484 | 5.94E-11 |
| ENSG00000285669 | gene_sourcehavana | 1.354871418 | 8.51E-12 |
| ENSG00000167552 | TUBA1A | 1.354637647 | 1.42E-69 |
| ENSG00000133134 | BEX2 | 1.354146926 | 0.001629 |
| ENSG00000125505 | MBOAT7 | 1.353557201 | 4.91E-84 |

| | | | |
|---|---|---|---|
| ENSG00000158023 | CFAP251 | 1.350955791 | 0.000607 |
| ENSG00000116199 | FAM20B | 1.350425056 | 9.03E-66 |
| ENSG00000233654 | NEMP2-DT | 1.349366796 | 0.01811 |
| ENSG00000251434 | gene_sourcehavana | 1.348504126 | 3.63E-05 |
| ENSG00000096433 | ITPR3 | 1.347634344 | 3.41E-17 |
| ENSG00000235272 | RAMACL | 1.346909052 | 0.014774 |
| ENSG00000164695 | CHMP4C | 1.345420994 | 1.03E-17 |
| ENSG00000272688 | gene_sourcehavana | 1.343080826 | 0.006553 |
| ENSG00000047634 | SCML1 | 1.339321318 | 3.57E-20 |
| ENSG00000205740 | gene_sourcehavana | 1.338651879 | 0.046423 |
| ENSG00000089847 | ANKRD24 | 1.338356278 | 0.000664 |
| ENSG00000092295 | TGM1 | 1.337650003 | 0.000224 |
| ENSG00000135127 | BICDL1 | 1.337455152 | 0.006235 |
| ENSG00000099326 | MZF1 | 1.33429046 | 2.91E-47 |
| ENSG00000110665 | C11orf21 | 1.334145906 | 2.62E-71 |
| ENSG00000163932 | PRKCD | 1.33364419 | 6.53E-46 |
| ENSG00000203865 | ATP1A1-AS1 | 1.333631192 | 3.62E-08 |
| ENSG00000114853 | ZBTB47 | 1.333325809 | 0.036039 |
| ENSG00000168802 | CHTF8 | 1.333193264 | 0.042863 |
| ENSG00000272379 | gene_sourcehavana | 1.333118078 | 0.042608 |
| ENSG00000288877 | gene_sourcehavana_ | 1.332175899 | 0.002073 |
| ENSG00000277224 | H2BC7 | 1.332095699 | 0.038715 |
| ENSG00000175482 | POLD4 | 1.331844426 | 3.24E-08 |
| ENSG00000169155 | ZBTB43 | 1.33162451 | 5.16E-32 |
| ENSG00000142669 | SH3BGRL3 | 1.331531254 | 3.28E-71 |
| ENSG00000187231 | SESTD1 | 1.331402197 | 0.011446 |
| ENSG00000023330 | ALAS1 | 1.331079659 | 8.32E-41 |
| ENSG00000115216 | NRBP1 | 1.328302167 | 8.45E-70 |
| ENSG00000164040 | PGRMC2 | 1.327090404 | 1.76E-99 |
| ENSG00000174600 | CMKLR1 | 1.325273407 | 0.01091 |
| ENSG00000181027 | FKRP | 1.323738751 | 3.08E-39 |
| ENSG00000099910 | KLHL22 | 1.320639896 | 3.17E-11 |
| ENSG00000273335 | gene_sourcehavana | 1.320477103 | 0.024003 |
| ENSG00000150760 | DOCK1 | 1.319760409 | 0.028805 |
| ENSG00000227403 | LINC01806 | 1.319436549 | 2.29E-05 |
| ENSG00000011422 | PLAUR | 1.314496198 | 7.83E-15 |
| ENSG00000270344 | POC1B-AS1 | 1.314144921 | 0.004843 |
| ENSG00000163297 | ANTXR2 | 1.31413254 | 5.16E-42 |
| ENSG00000170634 | ACYP2 | 1.31150725 | 1.41E-09 |
| ENSG00000183963 | SMTN | 1.310188584 | 1.80E-30 |
| ENSG00000079819 | EPB41L2 | 1.310123718 | 4.63E-06 |
| ENSG00000173933 | RBM4 | 1.309938437 | 1.98E-16 |
| ENSG00000133657 | ATP13A3 | 1.309549845 | 1.52E-68 |
| ENSG00000242028 | HYPK | 1.309497743 | 0.000496 |
| ENSG00000124721 | DNAH8 | 1.30934408 | 5.61E-05 |
| ENSG00000178498 | DTX3 | 1.309282469 | 1.22E-05 |
| ENSG00000213928 | IRF9 | 1.309005757 | 1.05E-06 |
| ENSG00000181035 | SLC25A42 | 1.307499926 | 1.43E-38 |
| ENSG00000133069 | TMCC2 | 1.307070128 | 5.60E-29 |

| | | | |
|---|---|---|---|
| ENSG00000171873 | ADRA1D | 1.306575444 | 0.039953 |
| ENSG00000198853 | RUSC2 | 1.306439737 | 4.27E-12 |
| ENSG00000141562 | NARF | 1.304297826 | 8.78E-65 |
| ENSG00000131941 | RHPN2 | 1.303997527 | 3.49E-06 |
| ENSG00000206503 | HLA-A | 1.303201764 | 3.01E-63 |
| ENSG00000074660 | SCARF1 | 1.302341367 | 5.63E-15 |
| ENSG00000187678 | SPRY4 | 1.301007597 | 2.98E-12 |
| ENSG00000104783 | KCNN4 | 1.300973142 | 8.01E-35 |
| ENSG00000173442 | EHBP1L1 | 1.3006713 | 3.40E-59 |
| ENSG00000144959 | NCEH1 | 1.300622589 | 3.80E-72 |
| ENSG00000177606 | JUN | 1.300559996 | 7.21E-13 |
| ENSG00000279602 | gene_sourcehavana | 1.299089028 | 0.004662 |
| ENSG00000255624 | C10orf88B | 1.298804744 | 4.98E-10 |
| ENSG00000273802 | H2BC8 | 1.298691874 | 8.40E-05 |
| ENSG00000255477 | gene_sourcehavana | 1.297544215 | 0.000573 |
| ENSG00000181350 | LRRC75A | 1.295652949 | 1.09E-13 |
| ENSG00000273136 | NBPF26 | 1.295020621 | 1.63E-06 |
| ENSG00000245552 | LNCRNA-IUR | 1.294804302 | 1.86E-11 |
| ENSG00000203879 | GDI1 | 1.2937998 | 1.62E-44 |
| ENSG00000243978 | RTL9 | 1.291193892 | 0.047685 |
| ENSG00000139324 | TMTC3 | 1.290883191 | 3.42E-28 |
| ENSG00000173926 | MARCHF3 | 1.289941546 | 1.96E-21 |
| ENSG00000155252 | PI4K2A | 1.289406038 | 2.59E-50 |
| ENSG00000145555 | MYO10 | 1.288836737 | 0.032517 |
| ENSG00000214558 | ZC3H11C | 1.288253064 | 0.0498 |
| ENSG00000052802 | MSMO1 | 1.28798191 | 6.38E-67 |
| ENSG00000263069 | RNF213-AS1 | 1.286195276 | 0.036067 |
| ENSG00000135924 | DNAJB2 | 1.285219572 | 1.25E-40 |
| ENSG00000173598 | NUDT4 | 1.285174101 | 9.74E-64 |
| ENSG00000135241 | PNPLA8 | 1.283537153 | 2.24E-49 |
| ENSG00000112212 | TSPO2 | 1.283047167 | 8.16E-32 |
| ENSG00000136940 | PDCL | 1.282492401 | 2.00E-76 |
| ENSG00000163376 | KBTBD8 | 1.281638033 | 1.41E-66 |
| ENSG00000135047 | CTSL | 1.281523932 | 3.93E-79 |
| ENSG00000128342 | LIF | 1.281424743 | 9.82E-21 |
| ENSG00000170266 | GLB1 | 1.280766728 | 3.98E-49 |
| ENSG00000130222 | GADD45G | 1.279672752 | 0.024271 |
| ENSG00000127125 | PPCS | 1.278320784 | 1.65E-36 |
| ENSG00000149212 | SESN3 | 1.277797833 | 1.33E-61 |
| ENSG00000124201 | ZNFX1 | 1.273815052 | 4.73E-48 |
| ENSG00000204386 | NEU1 | 1.273347157 | 4.15E-50 |
| ENSG00000251279 | SMIM15-AS1 | 1.271696074 | 0.045305 |
| ENSG00000290791 | gene_sourcehavana | 1.270682335 | 1.48E-07 |
| ENSG00000260329 | TMEM263-DT | 1.269010293 | 0.000149 |
| ENSG00000111726 | CMAS | 1.267945392 | 1.71E-41 |
| ENSG00000102265 | TIMP1 | 1.267157445 | 8.68E-44 |
| ENSG00000086065 | CHMP5 | 1.265300723 | 3.21E-43 |
| ENSG00000071242 | RPS6KA2 | 1.265162163 | 0.000163 |
| ENSG00000107829 | FBXW4 | 1.264427823 | 3.94E-39 |

| | | | |
|---|---|---|---|
| ENSG00000172331 | BPGM | 1.263527988 | 8.85E-34 |
| ENSG00000285693 | gene_sourcehavana | 1.262244256 | 0.014062 |
| ENSG00000163171 | CDC42EP3 | 1.260694935 | 0.001192 |
| ENSG00000198561 | CTNND1 | 1.260578591 | 2.87E-14 |
| ENSG00000163191 | S100A11 | 1.260515329 | 1.42E-26 |
| ENSG00000067082 | KLF6 | 1.258974662 | 1.05E-36 |
| ENSG00000161714 | PLCD3 | 1.255684942 | 4.68E-07 |
| ENSG00000123091 | RNF11 | 1.255086392 | 3.18E-48 |
| ENSG00000205464 | ATP6AP1L | 1.25366763 | 0.001087 |
| ENSG00000106804 | C5 | 1.250572386 | 0.006843 |
| ENSG00000155858 | LSM11 | 1.250033188 | 1.38E-28 |
| ENSG00000130653 | PNPLA7 | 1.248806031 | 0.012481 |
| ENSG00000206172 | HBA1 | 1.247324717 | 1.53E-11 |
| ENSG00000158301 | GPRASP2 | 1.246484816 | 2.65E-10 |
| ENSG00000117016 | RIMS3 | 1.246378415 | 2.16E-33 |
| ENSG00000290086 | gene_sourcehavana_ | 1.24614772 | 0.043003 |
| ENSG00000272140 | gene_sourcehavana | 1.246131492 | 0.00326 |
| ENSG00000084764 | MAPRE3 | 1.24548904 | 2.31E-16 |
| ENSG00000111885 | MAN1A1 | 1.245461446 | 5.29E-38 |
| ENSG00000068697 | LAPTM4A | 1.245281082 | 3.32E-40 |
| ENSG00000134107 | BHLHE40 | 1.245280697 | 1.38E-23 |
| ENSG00000171298 | GAA | 1.243374454 | 3.11E-16 |
| ENSG00000177169 | ULK1 | 1.242286764 | 1.54E-34 |
| ENSG00000134294 | SLC38A2 | 1.241699646 | 9.71E-52 |
| ENSG00000068878 | PSME4 | 1.241626817 | 2.45E-57 |
| ENSG00000100473 | COCH | 1.241525352 | 1.79E-46 |
| ENSG00000117758 | STX12 | 1.240286876 | 8.65E-32 |
| ENSG00000144674 | GOLGA4 | 1.240030151 | 7.47E-45 |
| ENSG00000130940 | CASZ1 | 1.239832984 | 4.32E-13 |
| ENSG00000033627 | ATP6V0A1 | 1.239690615 | 1.49E-52 |
| ENSG00000126088 | UROD | 1.239191878 | 5.81E-108 |
| ENSG00000173530 | TNFRSF10D | 1.238639075 | 8.63E-09 |
| ENSG00000106299 | WASL | 1.237875838 | 5.96E-23 |
| ENSG00000154620 | TMSB4Y | 1.237766356 | 0.002739 |
| ENSG00000289956 | gene_sourcehavana_ | 1.23749697 | 9.36E-06 |
| ENSG00000139410 | SDSL | 1.23610095 | 2.03E-15 |
| ENSG00000158373 | H2BC5 | 1.235950597 | 1.93E-10 |
| ENSG00000184350 | MRGPRE | 1.235832019 | 1.19E-51 |
| ENSG00000196218 | RYR1 | 1.235395623 | 0.000477 |
| ENSG00000140564 | FURIN | 1.235000437 | 7.83E-54 |
| ENSG00000196747 | H2AC13 | 1.234833764 | 0.000128 |
| ENSG00000165804 | ZNF219 | 1.23323661 | 9.89E-15 |
| ENSG00000001561 | ENPP4 | 1.231863204 | 7.51E-35 |
| ENSG00000259891 | gene_sourcehavana | 1.231528065 | 1.08E-05 |
| ENSG00000173611 | SCAI | 1.231158834 | 4.77E-39 |
| ENSG00000121064 | SCPEP1 | 1.230949895 | 5.55E-50 |
| ENSG00000104899 | AMH | 1.230700316 | 6.96E-05 |
| ENSG00000108551 | RASD1 | 1.230190701 | 5.49E-13 |
| ENSG00000101004 | NINL | 1.229883589 | 0.029509 |

| | | | |
|---|---|---|---|
| ENSG00000159023 | EPB41 | 1.22950484 | 5.22E-109 |
| ENSG00000212916 | MAP10 | 1.229110496 | 0.04044 |
| ENSG00000188747 | NOXA1 | 1.229102921 | 0.008295 |
| ENSG00000056558 | TRAF1 | 1.228372298 | 0.021684 |
| ENSG00000108187 | PBLD | 1.227038413 | 2.75E-07 |
| ENSG00000290877 | ROCK1P1 | 1.223847652 | 0.001306 |
| ENSG00000270629 | NBPF14 | 1.222591265 | 1.01E-06 |
| ENSG00000176809 | LRRC37A3 | 1.219821173 | 0.000216 |
| ENSG00000231663 | COA6-AS1 | 1.219365298 | 0.043685 |
| ENSG00000112511 | PHF1 | 1.219027404 | 2.43E-38 |
| ENSG00000256269 | HMBS | 1.217274746 | 1.96E-91 |
| ENSG00000164938 | TP53INP1 | 1.217132474 | 6.80E-39 |
| ENSG00000064201 | TSPAN32 | 1.216422725 | 4.61E-22 |
| ENSG00000067225 | PKM | 1.215367947 | 1.29E-93 |
| ENSG00000213533 | STIMATE | 1.212315844 | 0.018149 |
| ENSG00000049769 | PPP1R3F | 1.211439309 | 0.015795 |
| ENSG00000168779 | SHOX2 | 1.210374289 | 0.000122 |
| ENSG00000158769 | F11R | 1.205019413 | 2.65E-37 |
| ENSG00000132003 | ZSWIM4 | 1.205000684 | 1.55E-09 |
| ENSG00000153048 | CARHSP1 | 1.204973221 | 2.90E-28 |
| ENSG00000165244 | ZNF367 | 1.20366774 | 1.21E-45 |
| ENSG00000146676 | PURB | 1.20244712 | 2.80E-48 |
| ENSG00000284946 | gene_sourcehavana | 1.201840177 | 1.66E-16 |
| ENSG00000099785 | MARCHF2 | 1.20174288 | 3.09E-33 |
| ENSG00000141682 | PMAIP1 | 1.201490635 | 9.24E-15 |
| ENSG00000174358 | SLC6A19 | 1.200663986 | 1.34E-56 |
| ENSG00000126775 | ATG14 | 1.200383842 | 7.90E-52 |
| ENSG00000167065 | DUSP18 | 1.200311641 | 2.50E-12 |
| ENSG00000204520 | MICA | 1.200158286 | 1.53E-26 |
| ENSG00000174516 | PELI3 | 1.198366552 | 9.77E-05 |
| ENSG00000047597 | XK | 1.196149508 | 5.79E-75 |
| ENSG00000276903 | H2AC16 | 1.195452043 | 0.006357 |
| ENSG00000008056 | SYN1 | 1.194960904 | 1.04E-05 |
| ENSG00000164976 | MYORG | 1.19347068 | 0.011341 |
| ENSG00000111727 | HCFC2 | 1.191407102 | 3.81E-24 |
| ENSG00000135404 | CD63 | 1.190708651 | 2.98E-57 |
| ENSG00000082458 | DLG3 | 1.190150234 | 0.043575 |
| ENSG00000204116 | CHIC1 | 1.18891694 | 4.11E-20 |
| ENSG00000249898 | MCPH1-AS1 | 1.188548448 | 2.33E-09 |
| ENSG00000147454 | SLC25A37 | 1.188526167 | 2.38E-109 |
| ENSG00000075539 | FRYL | 1.188469716 | 9.10E-53 |
| ENSG00000178934 | LGALS7B | 1.188373159 | 3.39E-10 |
| ENSG00000140859 | KIFC3 | 1.187662604 | 5.17E-24 |
| ENSG00000272447 | gene_sourcehavana | 1.18751017 | 9.56E-05 |
| ENSG00000113361 | CDH6 | 1.187361302 | 4.82E-60 |
| ENSG00000119737 | GPR75 | 1.187208556 | 3.92E-09 |
| ENSG00000171928 | TVP23B | 1.186017423 | 5.52E-21 |
| ENSG00000188177 | ZC3H6 | 1.185533186 | 8.66E-18 |
| ENSG00000072071 | ADGRL1 | 1.185464832 | 6.12E-16 |

| | | | |
|---|---|---|---|
| ENSG00000277687 | gene_sourcehavana | 1.183793372 | 0.003009 |
| ENSG00000162366 | PDZK1IP1 | 1.183511035 | 3.09E-17 |
| ENSG00000178127 | NDUFV2 | 1.183317669 | 0.003998 |
| ENSG00000265590 | CFAP298-TCP10L | 1.180465697 | 1.29E-11 |
| ENSG00000006062 | MAP3K14 | 1.180016849 | 1.42E-05 |
| ENSG00000271869 | DCTN6-DT | 1.177943926 | 0.036714 |
| ENSG00000010810 | FYN | 1.177683892 | 0.014567 |
| ENSG00000197872 | CYRIA | 1.172047907 | 2.01E-36 |
| ENSG00000269176 | gene_sourcehavana | 1.171974646 | 0.00265 |
| ENSG00000196152 | ZNF79 | 1.171296785 | 2.32E-20 |
| ENSG00000166532 | RIMKLB | 1.171015403 | 3.67E-41 |
| ENSG00000105856 | HBP1 | 1.170391566 | 1.61E-41 |
| ENSG00000140931 | CMTM3 | 1.170386507 | 8.16E-05 |
| ENSG00000206341 | HLA-H | 1.169262056 | 0.004087 |
| ENSG00000228107 | gene_sourcehavana | 1.169186368 | 0.001344 |
| ENSG00000187134 | AKR1C1 | 1.168222033 | 3.00E-05 |
| ENSG00000232859 | LYRM9 | 1.167147694 | 0.001354 |
| ENSG00000111276 | CDKN1B | 1.167041049 | 1.00E-50 |
| ENSG00000237977 | EIF4HP2 | 1.165503526 | 4.85E-05 |
| ENSG00000171903 | CYP4F11 | 1.165186652 | 0.003366 |
| ENSG00000184678 | H2BC21 | 1.165140502 | 0.002277 |
| ENSG00000125772 | GPCPD1 | 1.163818489 | 3.36E-24 |
| ENSG00000119139 | TJP2 | 1.163290421 | 3.49E-09 |
| ENSG00000179954 | SSC5D | 1.16311595 | 0.017772 |
| ENSG00000261455 | LINC01003 | 1.162888833 | 0.00023 |
| ENSG00000196642 | RABL6 | 1.162519847 | 1.86E-49 |
| ENSG00000137331 | IER3 | 1.160797087 | 2.22E-05 |
| ENSG00000291043 | GLUD1P3 | 1.160238016 | 0.001326 |
| ENSG00000117155 | SSX2IP | 1.158892949 | 3.19E-50 |
| ENSG00000270872 | SRGAP2D | 1.15727691 | 0.000697 |
| ENSG00000158615 | PPP1R15B | 1.156264743 | 3.15E-58 |
| ENSG00000129028 | THAP10 | 1.155547169 | 4.23E-05 |
| ENSG00000065029 | ZNF76 | 1.155041574 | 1.28E-36 |
| ENSG00000083857 | FAT1 | 1.153892812 | 2.80E-06 |
| ENSG00000231856 | gene_sourcehavana | 1.153576473 | 0.008531 |
| ENSG00000267309 | ZNF566-AS1 | 1.153335517 | 0.019554 |
| ENSG00000286532 | PARTICL | 1.152106559 | 0.00012 |
| ENSG00000054965 | FAM168A | 1.151948218 | 1.16E-20 |
| ENSG00000142733 | MAP3K6 | 1.1513865 | 4.24E-19 |
| ENSG00000111540 | RAB5B | 1.151306914 | 8.03E-47 |
| ENSG00000137500 | CCDC90B | 1.150196501 | 4.70E-30 |
| ENSG00000277481 | PKD1L3 | 1.148627098 | 2.99E-05 |
| ENSG00000138495 | COX17 | 1.148559411 | 1.63E-26 |
| ENSG00000183856 | IQGAP3 | 1.148500159 | 3.98E-48 |
| ENSG00000233276 | GPX1 | 1.148375121 | 3.09E-13 |
| ENSG00000289339 | gene_sourcehavana_ | 1.147550907 | 0.003539 |
| ENSG00000143382 | ADAMTSL4 | 1.147040347 | 5.01E-35 |
| ENSG00000149591 | TAGLN | 1.146900348 | 5.38E-07 |
| ENSG00000155980 | KIF5A | 1.146249054 | 5.00E-43 |

| | | | |
|---|---|---|---|
| ENSG00000120688 | WBP4 | 1.145776135 | 2.66E-49 |
| ENSG00000291147 | gene_sourcehavana | 1.145748588 | 0.001563 |
| ENSG00000051382 | PIK3CB | 1.145100966 | 4.83E-35 |
| ENSG00000263232 | ATP5F1AP3 | 1.14505593 | 0.000301 |
| ENSG00000289625 | IUR1 | 1.144513447 | 0.005569 |
| ENSG00000259704 | gene_sourcehavana | 1.1428198 | 0.043301 |
| ENSG00000140009 | ESR2 | 1.142803019 | 1.31E-08 |
| ENSG00000106526 | ACTR3C | 1.141899898 | 2.16E-06 |
| ENSG00000288663 | gene_sourcehavana | 1.141225411 | 0.02159 |
| ENSG00000143774 | GUK1 | 1.140380772 | 5.29E-45 |
| ENSG00000244055 | gene_sourcehavana | 1.140360408 | 0.030085 |
| ENSG00000124357 | NAGK | 1.140267228 | 8.95E-29 |
| ENSG00000104177 | MYEF2 | 1.139127517 | 1.28E-14 |
| ENSG00000177951 | BET1L | 1.13911946 | 5.20E-33 |
| ENSG00000233609 | RPL10P19 | 1.137396419 | 0.002067 |
| ENSG00000133794 | BMAL1 | 1.136941413 | 7.22E-19 |
| ENSG00000156500 | PABIR3 | 1.136934819 | 1.41E-17 |
| ENSG00000180263 | FGD6 | 1.136250143 | 3.10E-15 |
| ENSG00000235823 | OLMALINC | 1.136236323 | 4.77E-08 |
| ENSG00000085788 | DDHD2 | 1.136072819 | 1.91E-61 |
| ENSG00000158578 | ALAS2 | 1.135664204 | 2.49E-49 |
| ENSG00000154640 | BTG3 | 1.134389442 | 1.63E-33 |
| ENSG00000185716 | MOSMO | 1.134096594 | 4.06E-14 |
| ENSG00000133107 | TRPC4 | 1.133805146 | 0.001507 |
| ENSG00000115159 | GPD2 | 1.133363159 | 7.14E-42 |
| ENSG00000040487 | SLC66A1 | 1.132959267 | 1.43E-20 |
| ENSG00000266236 | NARF-IT1 | 1.131954708 | 0.000165 |
| ENSG00000173264 | GPR137 | 1.131311944 | 7.44E-21 |
| ENSG00000241404 | EGFL8 | 1.130585272 | 2.11E-05 |
| ENSG00000197063 | MAFG | 1.129724992 | 1.31E-45 |
| ENSG00000156381 | ANKRD9 | 1.128796465 | 1.11E-26 |
| ENSG00000124098 | FAM210B | 1.127937996 | 1.45E-79 |
| ENSG00000144655 | CSRNP1 | 1.127604168 | 1.49E-17 |
| ENSG00000079482 | OPHN1 | 1.1272464 | 0.001225 |
| ENSG00000204334 | ERICH2 | 1.126461318 | 5.09E-11 |
| ENSG00000196263 | ZNF471 | 1.125133519 | 1.74E-07 |
| ENSG00000131100 | ATP6V1E1 | 1.124544776 | 1.82E-40 |
| ENSG00000224505 | HEXIM2-AS1 | 1.124223884 | 0.03741 |
| ENSG00000069956 | MAPK6 | 1.122148937 | 1.91E-52 |
| ENSG00000187699 | C2orf88 | 1.120700366 | 2.23E-31 |
| ENSG00000177432 | NAP1L5 | 1.119356347 | 8.25E-19 |
| ENSG00000198168 | SVIP | 1.118116421 | 1.03E-49 |
| ENSG00000100711 | ZFYVE21 | 1.117688073 | 3.74E-37 |
| ENSG00000197147 | LRRC8B | 1.117028189 | 1.30E-38 |
| ENSG00000069493 | CLEC2D | 1.116818388 | 8.65E-06 |
| ENSG00000240731 | gene_sourcehavana | 1.116227334 | 0.011081 |
| ENSG00000101255 | TRIB3 | 1.114364709 | 1.11E-07 |
| ENSG00000004809 | SLC22A16 | 1.113462222 | 1.01E-14 |
| ENSG00000130830 | MPP1 | 1.113290797 | 2.25E-70 |

| | | | |
|---|---|---|---|
| ENSG00000244274 | DBNDD2 | 1.112787156 | 0.043003 |
| ENSG00000083097 | DOP1A | 1.112616987 | 3.66E-27 |
| ENSG00000196233 | LCOR | 1.110008548 | 1.41E-32 |
| ENSG00000177595 | PIDD1 | 1.109967142 | 2.13E-25 |
| ENSG00000110429 | FBXO3 | 1.108417101 | 2.13E-33 |
| ENSG00000180769 | WDFY3-AS2 | 1.108094832 | 0.005867 |
| ENSG00000197153 | H3C12 | 1.108035356 | 0.01452 |
| ENSG00000232545 | gene_sourcehavana | 1.107651971 | 0.009927 |
| ENSG00000181904 | C5orf24 | 1.106983676 | 1.45E-36 |
| ENSG00000100478 | AP4S1 | 1.10646016 | 1.52E-11 |
| ENSG00000198039 | ZNF273 | 1.104443606 | 7.39E-22 |
| ENSG00000166483 | WEE1 | 1.10390353 | 8.28E-40 |
| ENSG00000185567 | AHNAK2 | 1.103427002 | 1.31E-22 |
| ENSG00000006576 | PHTF2 | 1.101286272 | 1.36E-49 |
| ENSG00000117616 | RSRP1 | 1.100938863 | 8.83E-49 |
| ENSG00000148488 | ST8SIA6 | 1.100845536 | 1.61E-34 |
| ENSG00000105443 | CYTH2 | 1.100123731 | 3.92E-38 |
| ENSG00000118292 | C1orf54 | 1.099976423 | 0.047356 |
| ENSG00000167565 | SERTAD3 | 1.099903189 | 1.91E-21 |
| ENSG00000288961 | gene_sourcehavana_ | 1.099211575 | 0.002007 |
| ENSG00000183718 | TRIM52 | 1.09864351 | 1.67E-25 |
| ENSG00000187902 | SHISA7 | 1.096666255 | 2.63E-33 |
| ENSG00000158109 | TPRG1L | 1.096632629 | 3.73E-36 |
| ENSG00000128245 | YWHAH | 1.094890244 | 1.87E-66 |
| ENSG00000166801 | FAM111A | 1.094676584 | 8.87E-50 |
| ENSG00000180979 | LRRC57 | 1.093331238 | 1.75E-22 |
| ENSG00000102760 | RGCC | 1.093310904 | 5.29E-40 |
| ENSG00000167193 | CRK | 1.093280776 | 2.95E-41 |
| ENSG00000197465 | GYPE | 1.092694788 | 1.62E-41 |
| ENSG00000224259 | LINC01133 | 1.092400706 | 3.89E-07 |
| ENSG00000092841 | MYL6 | 1.092340925 | 5.58E-52 |
| ENSG00000260231 | KDM7A-DT | 1.092212403 | 1.12E-05 |
| ENSG00000109680 | TBC1D19 | 1.092078238 | 0.019774 |
| ENSG00000140319 | SRP14 | 1.091490496 | 1.25E-48 |
| ENSG00000170382 | LRRN2 | 1.091167227 | 2.39E-12 |
| ENSG00000152484 | USP12 | 1.090775102 | 1.94E-37 |
| ENSG00000188536 | HBA2 | 1.090256289 | 6.46E-45 |
| ENSG00000166073 | GPR176 | 1.089904447 | 6.26E-17 |
| ENSG00000165782 | PIP4P1 | 1.089675345 | 1.04E-28 |
| ENSG00000170296 | GABARAP | 1.089621961 | 2.18E-08 |
| ENSG00000164543 | STK17A | 1.089357595 | 6.36E-36 |
| ENSG00000236104 | ZBTB22 | 1.08766712 | 1.22E-25 |
| ENSG00000286602 | gene_sourcehavana_ | 1.087157858 | 0.000205 |
| ENSG00000116815 | CD58 | 1.087093985 | 4.12E-37 |
| ENSG00000159348 | CYB5R1 | 1.084564362 | 6.18E-30 |
| ENSG00000173846 | PLK3 | 1.083440797 | 9.39E-12 |
| ENSG00000220205 | VAMP2 | 1.082984571 | 3.26E-25 |
| ENSG00000160991 | ORAI2 | 1.08261549 | 1.38E-27 |
| ENSG00000153214 | TMEM87B | 1.081909076 | 7.62E-31 |

| | | | |
|---|---|---|---|
| ENSG00000215811 | BTNL10P | 1.081728575 | 5.66E-05 |
| ENSG00000119801 | YPEL5 | 1.081674464 | 4.95E-44 |
| ENSG00000166946 | CCNDBP1 | 1.081395497 | 1.83E-58 |
| ENSG00000141854 | MISP3 | 1.080882511 | 4.33E-07 |
| ENSG00000122779 | TRIM24 | 1.080857089 | 1.36E-39 |
| ENSG00000150787 | PTS | 1.080476889 | 2.59E-17 |
| ENSG00000083290 | ULK2 | 1.080114504 | 6.26E-18 |
| ENSG00000206344 | HCG27 | 1.079269292 | 1.66E-06 |
| ENSG00000169967 | MAP3K2 | 1.078939367 | 7.06E-26 |
| ENSG00000182718 | ANXA2 | 1.076161214 | 5.16E-35 |
| ENSG00000152242 | C18orf25 | 1.071826662 | 1.49E-52 |
| ENSG00000157350 | ST3GAL2 | 1.071811964 | 1.17E-42 |
| ENSG00000214309 | MBLAC1 | 1.069931677 | 0.011472 |
| ENSG00000115998 | C2orf42 | 1.068900436 | 5.84E-19 |
| ENSG00000153814 | JAZF1 | 1.068247834 | 4.46E-54 |
| ENSG00000100196 | KDELR3 | 1.067280325 | 4.04E-09 |
| ENSG00000102362 | SYTL4 | 1.067007657 | 2.34E-32 |
| ENSG00000037042 | TUBG2 | 1.066216217 | 1.55E-53 |
| ENSG00000186591 | UBE2H | 1.065738508 | 1.03E-35 |
| ENSG00000277157 | H4C4 | 1.065070937 | 0.041013 |
| ENSG00000181061 | HIGD1A | 1.06305496 | 3.34E-27 |
| ENSG00000025423 | HSD17B6 | 1.062840333 | 0.033012 |
| ENSG00000106366 | SERPINE1 | 1.06170842 | 4.24E-06 |
| ENSG00000125812 | GZF1 | 1.061382312 | 3.89E-22 |
| ENSG00000167186 | COQ7 | 1.061283442 | 8.66E-20 |
| ENSG00000150457 | LATS2 | 1.061173034 | 1.44E-17 |
| ENSG00000197019 | SERTAD1 | 1.060716948 | 1.81E-11 |
| ENSG00000272106 | gene_sourcehavana | 1.06015694 | 0.000277 |
| ENSG00000213930 | GALT | 1.059391539 | 3.31E-06 |
| ENSG00000274925 | ZKSCAN2-DT | 1.059094717 | 2.91E-05 |
| ENSG00000167315 | ACAA2 | 1.05745298 | 2.43E-59 |
| ENSG00000174796 | THAP6 | 1.056765414 | 4.80E-14 |
| ENSG00000246705 | H2AJ | 1.055671744 | 4.69E-24 |
| ENSG00000110841 | PPFIBP1 | 1.054955703 | 0.024256 |
| ENSG00000005238 | ATOSB | 1.054350175 | 1.06E-21 |
| ENSG00000143811 | PYCR2 | 1.05387518 | 5.17E-38 |
| ENSG00000289432 | gene_sourcehavana_ | 1.053619465 | 0.000192 |
| ENSG00000198198 | SZT2 | 1.052931049 | 4.38E-33 |
| ENSG00000204592 | HLA-E | 1.051782964 | 2.42E-32 |
| ENSG00000169877 | AHSP | 1.05108682 | 9.44E-61 |
| ENSG00000135720 | DYNC1LI2 | 1.049537952 | 1.70E-42 |
| ENSG00000096070 | BRPF3 | 1.049275603 | 1.12E-44 |
| ENSG00000127220 | ABHD8 | 1.048959861 | 3.18E-11 |
| ENSG00000011426 | ANLN | 1.04823834 | 1.43E-67 |
| ENSG00000115170 | ACVR1 | 1.047324543 | 4.10E-07 |
| ENSG00000167536 | DHRS13 | 1.046837697 | 5.52E-33 |
| ENSG00000243335 | KCTD7 | 1.046819609 | 5.39E-15 |
| ENSG00000122203 | KIAA1191 | 1.046263589 | 2.96E-26 |
| ENSG00000239382 | ALKBH6 | 1.046082189 | 5.55E-06 |

| | | | |
|---|---|---|---|
| ENSG00000074370 | ATP2A3 | 1.045878808 | 0.03296 |
| ENSG00000188690 | UROS | 1.045777626 | 5.70E-65 |
| ENSG00000126903 | SLC10A3 | 1.044971782 | 1.46E-22 |
| ENSG00000198400 | NTRK1 | 1.043979004 | 1.62E-19 |
| ENSG00000196187 | TMEM63A | 1.043556217 | 3.82E-37 |
| ENSG00000135926 | TMBIM1 | 1.043455945 | 7.75E-61 |
| ENSG00000133247 | KMT5C | 1.041768984 | 7.92E-23 |
| ENSG00000227373 | RABGAP1L-DT | 1.041732495 | 0.043688 |
| ENSG00000198925 | ATG9A | 1.041195536 | 4.43E-41 |
| ENSG00000173064 | HECTD4 | 1.040423316 | 9.92E-39 |
| ENSG00000059378 | PARP12 | 1.038970601 | 1.85E-07 |
| ENSG00000204388 | HSPA1B | 1.038392976 | 9.84E-31 |
| ENSG00000102781 | KATNAL1 | 1.037379725 | 2.35E-26 |
| ENSG00000230615 | gene_sourcehavana | 1.03689768 | 0.023719 |
| ENSG00000105472 | CLEC11A | 1.036726998 | 3.14E-09 |
| ENSG00000102931 | ARL2BP | 1.03551797 | 2.82E-38 |
| ENSG00000204178 | MACO1 | 1.034823006 | 2.26E-47 |
| ENSG00000071967 | CYBRD1 | 1.034802803 | 2.99E-43 |
| ENSG00000198736 | MSRB1 | 1.0347551 | 9.90E-13 |
| ENSG00000260589 | STAM-DT | 1.034338037 | 0.047342 |
| ENSG00000100307 | CBX7 | 1.034171595 | 6.39E-13 |
| ENSG00000164327 | RICTOR | 1.033868681 | 3.91E-32 |
| ENSG00000021355 | SERPINB1 | 1.033844504 | 1.74E-17 |
| ENSG00000236397 | DDX11L2 | 1.032368696 | 0.004418 |
| ENSG00000160446 | ZDHHC12 | 1.032207654 | 4.37E-22 |
| ENSG00000197779 | ZNF81 | 1.031925377 | 1.69E-22 |
| ENSG00000183060 | LYSMD4 | 1.031799846 | 1.38E-17 |
| ENSG00000139112 | GABARAPL1 | 1.029943434 | 3.60E-09 |
| ENSG00000276644 | DACH1 | 1.027802535 | 2.04E-05 |
| ENSG00000147036 | LANCL3 | 1.027768486 | 4.49E-12 |
| ENSG00000139190 | VAMP1 | 1.027556961 | 1.96E-05 |
| ENSG00000173531 | MST1 | 1.02744816 | 1.48E-07 |
| ENSG00000156869 | FRRS1 | 1.027249085 | 2.71E-38 |
| ENSG00000100554 | ATP6V1D | 1.027157267 | 1.14E-25 |
| ENSG00000196576 | PLXNB2 | 1.026932616 | 2.19E-35 |
| ENSG00000253483 | gene_sourcehavana | 1.026798942 | 0.027921 |
| ENSG00000119242 | CCDC92 | 1.025218813 | 1.72E-06 |
| ENSG00000075426 | FOSL2 | 1.024402222 | 6.11E-05 |
| ENSG00000264112 | gene_sourcehavana | 1.023999987 | 5.26E-08 |
| ENSG00000154265 | ABCA5 | 1.023792644 | 0.001227 |
| ENSG00000154222 | CC2D1B | 1.022892567 | 5.70E-25 |
| ENSG00000161011 | SQSTM1 | 1.021839131 | 2.43E-07 |
| ENSG00000111912 | NCOA7 | 1.021388353 | 1.11E-27 |
| ENSG00000274290 | H2BC6 | 1.021377192 | 0.023108 |
| ENSG00000227533 | SLC2A1-DT | 1.020679154 | 0.018767 |
| ENSG00000137133 | HINT2 | 1.020634855 | 0.005295 |
| ENSG00000203705 | TATDN3 | 1.020516627 | 6.19E-22 |
| ENSG00000100949 | RABGGTA | 1.020174241 | 1.05E-23 |
| ENSG00000006712 | PAF1 | 1.01917209 | 1.04E-39 |

| | | | |
|---|---|---|---|
| ENSG00000090013 | BLVRB | 1.019170768 | 2.27E-57 |
| ENSG00000215908 | CROCCP2 | 1.017890879 | 4.92E-12 |
| ENSG00000135776 | ABCB10 | 1.017407399 | 1.40E-62 |
| ENSG00000235316 | DUSP8P5 | 1.01699341 | 0.006055 |
| ENSG00000117394 | SLC2A1 | 1.016670833 | 6.02E-34 |
| ENSG00000068394 | GPKOW | 1.016361648 | 8.85E-33 |
| ENSG00000104419 | NDRG1 | 1.015378461 | 6.65E-35 |
| ENSG00000279631 | gene_sourcehavana | 1.015120597 | 0.025015 |
| ENSG00000183508 | TENT5C | 1.013919925 | 4.62E-36 |
| ENSG00000133742 | CA1 | 1.013693848 | 2.21E-54 |
| ENSG00000185658 | BRWD1 | 1.013446424 | 6.66E-40 |
| ENSG00000207547 | MIR25 | 1.012786275 | 0.022496 |
| ENSG00000172461 | FUT9 | 1.01236574 | 3.53E-20 |
| ENSG00000157227 | MMP14 | 1.012276309 | 0.028766 |
| ENSG00000079691 | CARMIL1 | 1.011770825 | 4.40E-19 |
| ENSG00000151474 | FRMD4A | 1.011055017 | 3.61E-08 |
| ENSG00000198626 | RYR2 | 1.01036513 | 0.011498 |
| ENSG00000176912 | TYMSOS | 1.010104012 | 3.47E-05 |
| ENSG00000139168 | ZCRB1 | 1.009564833 | 2.78E-31 |
| ENSG00000099204 | ABLIM1 | 1.008041703 | 0.003095 |
| ENSG00000123643 | SLC36A1 | 1.006339972 | 4.38E-63 |
| ENSG00000247934 | gene_sourcehavana | 1.006294462 | 0.006141 |
| ENSG00000165650 | PDZD8 | 1.005547345 | 4.30E-44 |
| ENSG00000174405 | LIG4 | 1.005387451 | 1.53E-26 |
| ENSG00000155304 | HSPA13 | 1.003429128 | 3.03E-29 |
| ENSG00000157873 | TNFRSF14 | 1.002706113 | 1.94E-19 |
| ENSG00000183784 | DOCK8-AS1 | 1.002679763 | 0.017974 |
| ENSG00000166579 | NDEL1 | 1.002081733 | 2.49E-28 |
| ENSG00000178567 | EPM2AIP1 | 1.00185302 | 4.22E-28 |
| ENSG00000155090 | KLF10 | 1.001518628 | 4.56E-46 |
| ENSG00000126705 | AHDC1 | 1.000181242 | 1.51E-21 |
| ENSG00000106049 | HIBADH | -1.000293737 | 3.16E-17 |
| ENSG00000151176 | PLBD2 | -1.001755718 | 6.29E-38 |
| ENSG00000127533 | F2RL3 | -1.002126374 | 2.46E-09 |
| ENSG00000162694 | EXTL2 | -1.002617023 | 2.81E-12 |
| ENSG00000113273 | ARSB | -1.002823311 | 5.74E-19 |
| ENSG00000106603 | COA1 | -1.004728212 | 5.64E-23 |
| ENSG00000170322 | NFRKB | -1.005658606 | 4.44E-24 |
| ENSG00000119684 | MLH3 | -1.005827341 | 7.66E-17 |
| ENSG00000139722 | VPS37B | -1.006130923 | 5.96E-45 |
| ENSG00000170464 | DNAJC18 | -1.006982582 | 0.000803 |
| ENSG00000165185 | KIAA1958 | -1.007294335 | 2.54E-08 |
| ENSG00000112208 | BAG2 | -1.007463513 | 2.29E-23 |
| ENSG00000188921 | HACD4 | -1.007916489 | 6.36E-06 |
| ENSG00000126351 | THRA | -1.00792011 | 1.03E-21 |
| ENSG00000148296 | SURF6 | -1.00973821 | 7.13E-35 |
| ENSG00000170365 | SMAD1 | -1.009899722 | 5.97E-09 |
| ENSG00000240445 | FOXO3B | -1.010509916 | 1.45E-07 |
| ENSG00000186687 | LYRM7 | -1.010877628 | 1.46E-21 |

| | | | |
|---|---|---|---|
| ENSG00000100462 | PRMT5 | -1.011014898 | 3.02E-30 |
| ENSG00000165898 | ISCA2 | -1.011195762 | 5.16E-25 |
| ENSG00000183087 | GAS6 | -1.011931458 | 1.01E-13 |
| ENSG00000173801 | JUP | -1.012681399 | 1.68E-21 |
| ENSG00000142207 | URB1 | -1.012788468 | 4.04E-31 |
| ENSG00000288398 | gene_sourcehavana | -1.013559698 | 1.14E-05 |
| ENSG00000148288 | GBGT1 | -1.013859936 | 1.52E-42 |
| ENSG00000281398 | SNHG4 | -1.014127023 | 3.26E-16 |
| ENSG00000249741 | gene_sourcehavana | -1.014151642 | 0.013175 |
| ENSG00000205542 | TMSB4X | -1.014827245 | 4.93E-59 |
| ENSG00000088035 | ALG6 | -1.015215304 | 3.33E-09 |
| ENSG00000108592 | FTSJ3 | -1.01532416 | 1.52E-44 |
| ENSG00000176946 | THAP4 | -1.015446116 | 5.85E-35 |
| ENSG00000127314 | RAP1B | -1.016057938 | 3.47E-31 |
| ENSG00000117899 | MESD | -1.016162036 | 2.63E-33 |
| ENSG00000078081 | LAMP3 | -1.016532897 | 0.02707 |
| ENSG00000177370 | TIMM22 | -1.016677594 | 2.19E-20 |
| ENSG00000128928 | IVD | -1.016997873 | 6.62E-32 |
| ENSG00000104356 | POP1 | -1.018167976 | 1.57E-18 |
| ENSG00000196950 | SLC39A10 | -1.018265814 | 4.89E-25 |
| ENSG00000130382 | MLLT1 | -1.018551934 | 9.31E-26 |
| ENSG00000172889 | EGFL7 | -1.01933728 | 3.87E-33 |
| ENSG00000204264 | PSMB8 | -1.019567153 | 5.81E-09 |
| ENSG00000011143 | MKS1 | -1.02060654 | 1.73E-19 |
| ENSG00000130300 | PLVAP | -1.020950283 | 0.027667 |
| ENSG00000173145 | NOC3L | -1.021196603 | 1.47E-30 |
| ENSG00000254682 | gene_sourcehavana | -1.021844656 | 0.002486 |
| ENSG00000125827 | TMX4 | -1.022691742 | 4.10E-40 |
| ENSG00000117597 | UTP25 | -1.022820939 | 1.16E-35 |
| ENSG00000274383 | gene_sourcehavana | -1.023080501 | 0.031444 |
| ENSG00000198369 | SPRED2 | -1.023270351 | 1.62E-31 |
| ENSG00000100364 | KIAA0930 | -1.023466843 | 2.39E-07 |
| ENSG00000184216 | IRAK1 | -1.023518833 | 8.79E-47 |
| ENSG00000058056 | USP13 | -1.02550768 | 0.000944 |
| ENSG00000134490 | TMEM241 | -1.025754919 | 8.02E-10 |
| ENSG00000126522 | ASL | -1.025811367 | 2.40E-18 |
| ENSG00000138640 | FAM13A | -1.026666114 | 0.002739 |
| ENSG00000137486 | ARRB1 | -1.026704348 | 7.33E-14 |
| ENSG00000122884 | P4HA1 | -1.027584618 | 2.95E-13 |
| ENSG00000178802 | MPI | -1.027855301 | 3.19E-07 |
| ENSG00000236144 | TMEM147-AS1 | -1.028872403 | 9.30E-13 |
| ENSG00000143207 | COP1 | -1.029688347 | 3.16E-19 |
| ENSG00000168394 | TAP1 | -1.029726233 | 1.63E-12 |
| ENSG00000145214 | DGKQ | -1.030747738 | 4.87E-15 |
| ENSG00000172292 | CERS6 | -1.031629014 | 3.14E-21 |
| ENSG00000168286 | THAP11 | -1.031928647 | 2.82E-18 |
| ENSG00000168264 | IRF2BP2 | -1.032240932 | 7.16E-50 |
| ENSG00000112425 | EPM2A | -1.032585247 | 8.90E-06 |
| ENSG00000142089 | IFITM3 | -1.032728734 | 0.017009 |

| | | | |
|---|---|---|---|
| ENSG00000049167 | ERCC8 | -1.032987553 | 6.48E-16 |
| ENSG00000184492 | FOXD4L1 | -1.033742637 | 1.75E-05 |
| ENSG00000213965 | NUDT19 | -1.036481093 | 1.05E-26 |
| ENSG00000138617 | PARP16 | -1.037747687 | 3.77E-09 |
| ENSG00000136877 | FPGS | -1.038233888 | 5.24E-30 |
| ENSG00000084693 | AGBL5 | -1.039542119 | 5.85E-19 |
| ENSG00000174282 | ZBTB4 | -1.041050992 | 3.05E-30 |
| ENSG00000083635 | NUFIP1 | -1.041052937 | 9.09E-14 |
| ENSG00000122188 | LAX1 | -1.0411165 | 0.001271 |
| ENSG00000085117 | CD82 | -1.041901159 | 3.38E-45 |
| ENSG00000267575 | LINC02987 | -1.042845218 | 6.97E-10 |
| ENSG00000140382 | HMG20A | -1.043940224 | 1.04E-26 |
| ENSG00000219665 | ZNF433-AS1 | -1.044017102 | 0.000127 |
| ENSG00000197016 | ZNF470 | -1.04443385 | 4.34E-08 |
| ENSG00000176438 | SYNE3 | -1.046317704 | 4.40E-19 |
| ENSG00000263731 | gene_sourcehavana | -1.046370918 | 0.00346 |
| ENSG00000144867 | SRPRB | -1.046420333 | 9.17E-33 |
| ENSG00000152782 | PANK1 | -1.047362968 | 1.40E-06 |
| ENSG00000182108 | DEXI | -1.047455662 | 1.19E-24 |
| ENSG00000141295 | SCRN2 | -1.047712554 | 3.16E-07 |
| ENSG00000073060 | SCARB1 | -1.048082161 | 1.02E-37 |
| ENSG00000100225 | FBXO7 | -1.048440914 | 2.00E-45 |
| ENSG00000232445 | EMSLR | -1.048465659 | 2.26E-16 |
| ENSG00000166321 | NUDT13 | -1.049901604 | 0.023643 |
| ENSG00000262814 | MRPL12 | -1.04991402 | 3.58E-11 |
| ENSG00000291317 | TMEM276 | -1.050945395 | 0.00053 |
| ENSG00000100612 | DHRS7 | -1.052362899 | 6.62E-16 |
| ENSG00000101407 | TTI1 | -1.052834345 | 5.09E-28 |
| ENSG00000250479 | CHCHD10 | -1.053051468 | 1.24E-19 |
| ENSG00000115738 | ID2 | -1.053106677 | 1.47E-67 |
| ENSG00000171714 | ANO5 | -1.054030096 | 3.60E-13 |
| ENSG00000198673 | TAFA2 | -1.054142778 | 1.15E-06 |
| ENSG00000164086 | DUSP7 | -1.054965423 | 4.23E-12 |
| ENSG00000141576 | RNF157 | -1.055360416 | 1.66E-15 |
| ENSG00000173218 | VANGL1 | -1.056408786 | 2.19E-20 |
| ENSG00000189007 | ADAT2 | -1.057514434 | 1.89E-24 |
| ENSG00000166963 | MAP1A | -1.057897214 | 1.01E-10 |
| ENSG00000143819 | EPHX1 | -1.060104553 | 1.41E-18 |
| ENSG00000126457 | PRMT1 | -1.061064197 | 5.94E-38 |
| ENSG00000234261 | gene_sourcehavana | -1.061792492 | 1.42E-07 |
| ENSG00000118242 | MREG | -1.063025479 | 0.003024 |
| ENSG00000109062 | NHERF1 | -1.063740371 | 3.29E-22 |
| ENSG00000175538 | KCNE3 | -1.063839702 | 2.55E-05 |
| ENSG00000108091 | CCDC6 | -1.063868478 | 7.45E-44 |
| ENSG00000224078 | SNHG14 | -1.064351359 | 3.02E-30 |
| ENSG00000235092 | ID2-AS1 | -1.065116802 | 3.69E-07 |
| ENSG00000158292 | GPR153 | -1.065471018 | 5.51E-06 |
| ENSG00000163754 | GYG1 | -1.06620721 | 8.84E-23 |
| ENSG00000137073 | UBAP2 | -1.066254754 | 1.81E-35 |

| | | | |
|---|---|---|---|
| ENSG00000151229 | SLC2A13 | -1.066277067 | 1.57E-10 |
| ENSG00000198300 | PEG3 | -1.066772336 | 9.93E-15 |
| ENSG00000084774 | CAD | -1.066936015 | 1.26E-30 |
| ENSG00000100100 | PIK3IP1 | -1.066947053 | 1.26E-05 |
| ENSG00000162735 | PEX19 | -1.066960139 | 3.76E-16 |
| ENSG00000175602 | CCDC85B | -1.067643833 | 0.011843 |
| ENSG00000148187 | MRRF | -1.067862873 | 1.27E-19 |
| ENSG00000205981 | DNAJC19 | -1.067921474 | 2.07E-24 |
| ENSG00000138162 | TACC2 | -1.067934841 | 1.31E-35 |
| ENSG00000138074 | SLC5A6 | -1.068434796 | 3.92E-29 |
| ENSG00000225264 | ZNRF2P2 | -1.069016161 | 3.39E-05 |
| ENSG00000100350 | FOXRED2 | -1.07019745 | 6.29E-28 |
| ENSG00000163626 | COX18 | -1.071356188 | 7.74E-20 |
| ENSG00000213281 | NRAS | -1.071504602 | 1.48E-42 |
| ENSG00000235961 | PNMA6A | -1.072232941 | 0.000532 |
| ENSG00000171604 | CXXC5 | -1.07257315 | 1.97E-21 |
| ENSG00000140876 | NUDT7 | -1.072733683 | 6.13E-06 |
| ENSG00000132004 | FBXW9 | -1.073179495 | 6.84E-12 |
| ENSG00000108924 | HLF | -1.073788738 | 0.001378 |
| ENSG00000180817 | PPA1 | -1.074350247 | 1.07E-36 |
| ENSG00000165832 | TRUB1 | -1.074920899 | 1.43E-22 |
| ENSG00000103037 | SETD6 | -1.075825208 | 6.84E-14 |
| ENSG00000132330 | SCLY | -1.076638762 | 2.92E-05 |
| ENSG00000117174 | ZNHIT6 | -1.077035916 | 2.67E-34 |
| ENSG00000154359 | LONRF1 | -1.077458712 | 3.64E-11 |
| ENSG00000113790 | EHHADH | -1.077938911 | 2.66E-14 |
| ENSG00000167646 | DNAAF3 | -1.078831361 | 0.00653 |
| ENSG00000267278 | MAP3K14-AS1 | -1.078896849 | 1.19E-05 |
| ENSG00000122729 | ACO1 | -1.079116912 | 1.43E-37 |
| ENSG00000080503 | SMARCA2 | -1.079604867 | 1.13E-07 |
| ENSG00000168936 | TMEM129 | -1.079676338 | 3.82E-14 |
| ENSG00000006047 | YBX2 | -1.080889256 | 1.13E-43 |
| ENSG00000162039 | MEIOB | -1.080926382 | 0.004419 |
| ENSG00000141425 | RPRD1A | -1.082249944 | 1.70E-50 |
| ENSG00000109323 | MANBA | -1.082953032 | 8.27E-14 |
| ENSG00000149313 | AASDHPPT | -1.083595564 | 8.66E-35 |
| ENSG00000136169 | SETDB2 | -1.083734061 | 1.51E-17 |
| ENSG00000157404 | KIT | -1.083820393 | 3.84E-90 |
| ENSG00000254093 | PINX1 | -1.085871247 | 9.18E-20 |
| ENSG00000273149 | gene_sourcehavana | -1.086473268 | 0.031841 |
| ENSG00000100099 | HPS4 | -1.086994144 | 1.32E-49 |
| ENSG00000103966 | EHD4 | -1.08747689 | 1.18E-13 |
| ENSG00000187792 | ZNF70 | -1.087965643 | 5.25E-06 |
| ENSG00000173588 | CEP83 | -1.088134706 | 7.84E-19 |
| ENSG00000213398 | LCAT | -1.088432279 | 0.003832 |
| ENSG00000288701 | PRRC2B | -1.088628478 | 1.05E-55 |
| ENSG00000214425 | LRRC37A4P | -1.089066132 | 6.71E-08 |
| ENSG00000196843 | ARID5A | -1.089368438 | 8.88E-05 |
| ENSG00000214194 | SMIM30 | -1.089450044 | 1.50E-43 |

| | | | |
|---|---|---|---|
| ENSG00000152056 | AP1S3 | -1.09029997 | 0.004584 |
| ENSG00000250159 | PKD2L2-DT | -1.090572452 | 0.011803 |
| ENSG00000196653 | ZNF502 | -1.090580948 | 0.014924 |
| ENSG00000139597 | N4BP2L1 | -1.091025465 | 6.04E-09 |
| ENSG00000150977 | RILPL2 | -1.091059813 | 2.61E-19 |
| ENSG00000165506 | DNAAF2 | -1.091060284 | 1.52E-14 |
| ENSG00000272391 | POM121C | -1.091096951 | 1.46E-29 |
| ENSG00000178105 | DDX10 | -1.091624389 | 1.62E-17 |
| ENSG00000215845 | TSTD1 | -1.092487617 | 0.010521 |
| ENSG00000122873 | CISD1 | -1.092702317 | 1.86E-19 |
| ENSG00000139083 | ETV6 | -1.093483659 | 9.51E-10 |
| ENSG00000215417 | MIR17HG | -1.09348859 | 1.23E-05 |
| ENSG00000169860 | P2RY1 | -1.094208876 | 2.84E-11 |
| ENSG00000005187 | ACSM3 | -1.094533051 | 1.52E-16 |
| ENSG00000129562 | DAD1 | -1.096245426 | 1.72E-06 |
| ENSG00000151552 | QDPR | -1.09625753 | 9.71E-28 |
| ENSG00000068024 | HDAC4 | -1.09697844 | 1.78E-25 |
| ENSG00000141391 | PRELID3A | -1.098250075 | 0.017588 |
| ENSG00000182165 | TP53TG1 | -1.098984643 | 0.00244 |
| ENSG00000165271 | NOL6 | -1.100170866 | 3.65E-38 |
| ENSG00000167775 | CD320 | -1.10078779 | 5.48E-25 |
| ENSG00000172728 | FUT10 | -1.100847833 | 7.64E-15 |
| ENSG00000142684 | ZNF593 | -1.100963322 | 1.00E-18 |
| ENSG00000109586 | GALNT7 | -1.101997282 | 2.08E-29 |
| ENSG00000105755 | ETHE1 | -1.102212686 | 1.97E-07 |
| ENSG00000164626 | KCNK5 | -1.102237501 | 7.41E-32 |
| ENSG00000115419 | GLS | -1.102609447 | 6.90E-22 |
| ENSG00000225173 | gene_sourcehavana | -1.102936597 | 0.040329 |
| ENSG00000258920 | FOXN3-AS1 | -1.105281335 | 0.048108 |
| ENSG00000130540 | SULT4A1 | -1.106201668 | 0.007914 |
| ENSG00000118600 | RXYLT1 | -1.107421242 | 1.71E-17 |
| ENSG00000227704 | gene_sourcehavana | -1.107545657 | 0.022492 |
| ENSG00000124406 | ATP8A1 | -1.108922647 | 0.01879 |
| ENSG00000104369 | JPH1 | -1.109186669 | 5.58E-07 |
| ENSG00000168916 | ZNF608 | -1.109372628 | 0.000648 |
| ENSG00000101911 | PRPS2 | -1.109661621 | 2.37E-27 |
| ENSG00000103253 | HAGHL | -1.109948738 | 5.22E-14 |
| ENSG00000275342 | PRAG1 | -1.110645714 | 1.31E-16 |
| ENSG00000157870 | PRXL2B | -1.111913103 | 5.08E-13 |
| ENSG00000107954 | NEURL1 | -1.112092472 | 2.22E-16 |
| ENSG00000130338 | TULP4 | -1.112381332 | 4.31E-27 |
| ENSG00000289011 | gene_sourcehavana_ | -1.113284542 | 0.030787 |
| ENSG00000172115 | CYCS | -1.114110621 | 1.68E-52 |
| ENSG00000262585 | LINC01979 | -1.114291942 | 0.029099 |
| ENSG00000150867 | PIP4K2A | -1.115094625 | 1.91E-49 |
| ENSG00000131844 | MCCC2 | -1.116302124 | 5.99E-35 |
| ENSG00000035141 | FAM136A | -1.116839607 | 2.79E-38 |
| ENSG00000231527 | FAM27C | -1.116968178 | 0.023074 |
| ENSG00000235852 | gene_sourcehavana | -1.117775487 | 0.04965 |

| | | | |
|---|---|---|---|
| ENSG00000203995 | ZYG11A | -1.118595782 | 7.18E-16 |
| ENSG00000171016 | PYGO1 | -1.11927095 | 4.65E-16 |
| ENSG00000152465 | NMT2 | -1.120900595 | 5.17E-10 |
| ENSG00000198542 | ITGBL1 | -1.121552442 | 2.08E-24 |
| ENSG00000134987 | WDR36 | -1.122844298 | 7.55E-39 |
| ENSG00000105364 | MRPL4 | -1.123029641 | 1.73E-30 |
| ENSG00000185480 | PARPBP | -1.123871393 | 1.98E-26 |
| ENSG00000279369 | gene_sourcehavana | -1.12472682 | 1.92E-22 |
| ENSG00000105186 | ANKRD27 | -1.125640741 | 2.79E-62 |
| ENSG00000268555 | gene_sourcehavana | -1.125655895 | 3.39E-14 |
| ENSG00000172578 | KLHL6 | -1.126392372 | 1.43E-31 |
| ENSG00000215883 | CYB5RL | -1.126547854 | 1.37E-18 |
| ENSG00000144120 | TMEM177 | -1.12802637 | 6.34E-15 |
| ENSG00000130707 | ASS1 | -1.128099204 | 1.23E-05 |
| ENSG00000008382 | MPND | -1.128619489 | 2.50E-19 |
| ENSG00000237499 | WAKMAR2 | -1.129159549 | 0.035859 |
| ENSG00000135632 | SMYD5 | -1.129935257 | 2.62E-21 |
| ENSG00000134516 | DOCK2 | -1.130722814 | 3.21E-40 |
| ENSG00000156052 | GNAQ | -1.130912633 | 4.00E-66 |
| ENSG00000072135 | PTPN18 | -1.131191809 | 2.24E-39 |
| ENSG00000260804 | LINC01963 | -1.132626252 | 2.29E-06 |
| ENSG00000077348 | EXOSC5 | -1.13265595 | 1.61E-19 |
| ENSG00000149547 | EI24 | -1.132898561 | 1.24E-54 |
| ENSG00000170921 | TANC2 | -1.133236188 | 3.50E-22 |
| ENSG00000160255 | ITGB2 | -1.133578705 | 3.48E-09 |
| ENSG00000162004 | CCDC78 | -1.133806854 | 8.84E-07 |
| ENSG00000120685 | PROSER1 | -1.133977925 | 1.99E-33 |
| ENSG00000138698 | RAP1GDS1 | -1.134805562 | 6.01E-24 |
| ENSG00000159176 | CSRP1 | -1.136072458 | 1.07E-40 |
| ENSG00000111679 | PTPN6 | -1.136165439 | 2.98E-26 |
| ENSG00000119401 | TRIM32 | -1.136938886 | 4.28E-19 |
| ENSG00000139668 | WDFY2 | -1.138306642 | 5.58E-16 |
| ENSG00000114383 | TUSC2 | -1.139263603 | 1.44E-19 |
| ENSG00000120800 | UTP20 | -1.139354629 | 2.23E-51 |
| ENSG00000156958 | GALK2 | -1.140768972 | 1.07E-22 |
| ENSG00000204228 | HSD17B8 | -1.140923347 | 8.81E-13 |
| ENSG00000125871 | MGME1 | -1.142273359 | 5.73E-46 |
| ENSG00000185621 | LMLN | -1.142427502 | 3.84E-22 |
| ENSG00000166002 | SMCO4 | -1.143155319 | 0.001467 |
| ENSG00000117143 | UAP1 | -1.144559395 | 1.42E-32 |
| ENSG00000290948 | gene_sourcehavana | -1.146026673 | 3.99E-06 |
| ENSG00000128626 | MRPS12 | -1.146960464 | 4.34E-37 |
| ENSG00000167700 | MFSD3 | -1.147706477 | 9.05E-14 |
| ENSG00000100285 | NEFH | -1.148087436 | 2.68E-52 |
| ENSG00000160293 | VAV2 | -1.148310751 | 1.78E-13 |
| ENSG00000075651 | PLD1 | -1.148333967 | 1.33E-34 |
| ENSG00000184220 | CMSS1 | -1.149113828 | 3.32E-38 |
| ENSG00000291223 | FAM86DP | -1.149118592 | 3.35E-13 |
| ENSG00000005810 | MYCBP2 | -1.149204193 | 8.20E-34 |

| | | | |
|---|---|---|---|
| ENSG00000182612 | TSPAN10 | -1.149283992 | 0.001026 |
| ENSG00000287038 | gene_sourcehavana | -1.149408726 | 1.21E-15 |
| ENSG00000131467 | PSME3 | -1.150809261 | 1.16E-39 |
| ENSG00000185875 | THNSL1 | -1.151519577 | 1.60E-19 |
| ENSG00000064995 | TAF11 | -1.151596634 | 1.72E-34 |
| ENSG00000110721 | CHKA | -1.151608573 | 1.73E-30 |
| ENSG00000289984 | gene_sourcehavana_ | -1.153068687 | 0.031195 |
| ENSG00000235314 | LINC00957 | -1.156066925 | 1.38E-07 |
| ENSG00000146416 | AIG1 | -1.156477503 | 2.35E-20 |
| ENSG00000196628 | TCF4 | -1.157846713 | 8.37E-29 |
| ENSG00000257337 | TNS2-AS1 | -1.158195853 | 0.011828 |
| ENSG00000125841 | NRSN2 | -1.160027897 | 7.50E-09 |
| ENSG00000238098 | ABCA17P | -1.161097829 | 4.11E-12 |
| ENSG00000089351 | GRAMD1A | -1.162767732 | 1.45E-34 |
| ENSG00000057252 | SOAT1 | -1.163759056 | 2.00E-15 |
| ENSG00000118513 | MYB | -1.166787728 | 3.15E-73 |
| ENSG00000056736 | IL17RB | -1.167532361 | 1.87E-05 |
| ENSG00000196313 | POM121 | -1.168171805 | 2.43E-29 |
| ENSG00000166197 | NOLC1 | -1.168654375 | 2.65E-48 |
| ENSG00000255471 | PRSS23-AS1 | -1.168939705 | 0.001002 |
| ENSG00000197860 | SGTB | -1.169830448 | 4.71E-07 |
| ENSG00000186469 | GNG2 | -1.169974165 | 3.87E-38 |
| ENSG00000100105 | PATZ1 | -1.169977128 | 1.43E-37 |
| ENSG00000136933 | RABEPK | -1.170549778 | 7.28E-11 |
| ENSG00000279044 | gene_sourcehavana | -1.170827662 | 0.047735 |
| ENSG00000158716 | DUSP23 | -1.171461767 | 1.43E-11 |
| ENSG00000085998 | POMGNT1 | -1.171766187 | 4.39E-17 |
| ENSG00000175832 | ETV4 | -1.172077489 | 3.01E-05 |
| ENSG00000158747 | NBL1 | -1.172574416 | 2.47E-43 |
| ENSG00000238243 | OR2W3 | -1.174317416 | 5.54E-08 |
| ENSG00000204160 | ZDHHC18 | -1.174794089 | 1.38E-19 |
| ENSG00000169220 | RGS14 | -1.174829467 | 4.68E-18 |
| ENSG00000108578 | BLMH | -1.174969304 | 1.43E-28 |
| ENSG00000134215 | VAV3 | -1.175634346 | 1.61E-05 |
| ENSG00000137267 | TUBB2A | -1.175723743 | 1.79E-33 |
| ENSG00000158195 | WASF2 | -1.177467166 | 2.72E-72 |
| ENSG00000130270 | ATP8B3 | -1.177720378 | 1.41E-25 |
| ENSG00000076248 | UNG | -1.177863598 | 9.82E-39 |
| ENSG00000088543 | C3orf18 | -1.178276011 | 0.002515 |
| ENSG00000182957 | SPATA13 | -1.179531479 | 4.40E-17 |
| ENSG00000028277 | POU2F2 | -1.181463326 | 8.93E-05 |
| ENSG00000012232 | EXTL3 | -1.182088535 | 5.09E-40 |
| ENSG00000115318 | LOXL3 | -1.182277685 | 0.018538 |
| ENSG00000134333 | LDHA | -1.182407876 | 4.71E-06 |
| ENSG00000198363 | ASPH | -1.182968031 | 1.95E-18 |
| ENSG00000089057 | SLC23A2 | -1.183483254 | 1.43E-11 |
| ENSG00000146828 | SLC12A9 | -1.184793632 | 1.05E-18 |
| ENSG00000151748 | SAV1 | -1.185181648 | 2.35E-37 |
| ENSG00000114850 | SSR3 | -1.185521974 | 1.99E-50 |

| | | | |
|---|---|---|---|
| ENSG00000154217 | PITPNC1 | -1.186998719 | 4.07E-18 |
| ENSG00000184371 | CSF1 | -1.187174773 | 3.53E-34 |
| ENSG00000180822 | PSMG4 | -1.187769006 | 8.84E-26 |
| ENSG00000132881 | CPLANE2 | -1.187975539 | 2.57E-06 |
| ENSG00000068654 | POLR1A | -1.188207722 | 1.02E-48 |
| ENSG00000085644 | ZNF213 | -1.189474918 | 6.41E-15 |
| ENSG00000152147 | GEMIN6 | -1.189707455 | 4.09E-15 |
| ENSG00000069482 | GAL | -1.18973536 | 1.23E-22 |
| ENSG00000027001 | MIPEP | -1.190056464 | 1.45E-40 |
| ENSG00000077713 | SLC25A43 | -1.190189414 | 2.93E-14 |
| ENSG00000138738 | PRDM5 | -1.193489862 | 1.31E-10 |
| ENSG00000121057 | AKAP1 | -1.193706442 | 3.05E-35 |
| ENSG00000291127 | ANKRD36BP2 | -1.193901578 | 4.28E-13 |
| ENSG00000153767 | GTF2E1 | -1.194474106 | 5.12E-28 |
| ENSG00000196776 | CD47 | -1.194710716 | 1.96E-73 |
| ENSG00000184178 | SCFD2 | -1.194790347 | 6.86E-22 |
| ENSG00000186326 | RGS9BP | -1.195336306 | 0.008104 |
| ENSG00000196943 | NOP9 | -1.195698778 | 2.68E-37 |
| ENSG00000175220 | ARHGAP1 | -1.19640106 | 3.82E-30 |
| ENSG00000265688 | MILIP | -1.197870789 | 1.93E-07 |
| ENSG00000081154 | PCNP | -1.198064705 | 4.74E-38 |
| ENSG00000176105 | YES1 | -1.198159412 | 2.10E-23 |
| ENSG00000189046 | ALKBH2 | -1.198827156 | 1.37E-23 |
| ENSG00000163872 | YEATS2 | -1.199316433 | 1.86E-30 |
| ENSG00000104983 | CCDC61 | -1.199815657 | 8.94E-12 |
| ENSG00000125910 | S1PR4 | -1.199982719 | 6.63E-33 |
| ENSG00000168564 | CDKN2AIP | -1.201458429 | 1.27E-63 |
| ENSG00000060642 | PIGV | -1.201462952 | 1.70E-17 |
| ENSG00000290058 | gene_sourcehavana_ | -1.201730457 | 2.67E-08 |
| ENSG00000144485 | HES6 | -1.201905438 | 3.03E-32 |
| ENSG00000244187 | TMEM141 | -1.202652553 | 4.00E-17 |
| ENSG00000170854 | RIOX2 | -1.203495575 | 3.05E-35 |
| ENSG00000162999 | DUSP19 | -1.203656445 | 5.33E-05 |
| ENSG00000204681 | GABBR1 | -1.205321343 | 0.016175 |
| ENSG00000118420 | UBE3D | -1.205524043 | 1.82E-16 |
| ENSG00000183092 | BEGAIN | -1.205532205 | 3.13E-08 |
| ENSG00000103356 | EARS2 | -1.206255801 | 1.42E-26 |
| ENSG00000278571 | MIR7161 | -1.2069938 | 0.013139 |
| ENSG00000251209 | LINC00923 | -1.207047473 | 0.000475 |
| ENSG00000162337 | LRP5 | -1.207332283 | 2.98E-52 |
| ENSG00000224536 | CSRP1-AS1 | -1.208418669 | 0.005716 |
| ENSG00000157654 | PALM2AKAP2 | -1.208509551 | 1.93E-83 |
| ENSG00000141968 | VAV1 | -1.209380179 | 9.05E-27 |
| ENSG00000212719 | LINC02693 | -1.209449271 | 1.88E-08 |
| ENSG00000169299 | PGM2 | -1.210804163 | 9.27E-46 |
| ENSG00000173894 | CBX2 | -1.211124015 | 2.59E-18 |
| ENSG00000020633 | RUNX3 | -1.213797416 | 9.81E-17 |
| ENSG00000122877 | EGR2 | -1.214894955 | 6.74E-16 |
| ENSG00000167880 | EVPL | -1.215484547 | 2.15E-25 |

| | | | |
|---|---|---|---|
| ENSG00000171130 | ATP6V0E2 | -1.2155514 | 2.69E-15 |
| ENSG00000139835 | GRTP1 | -1.215702339 | 0.030195 |
| ENSG00000289145 | gene_sourcehavana_ | -1.216634999 | 0.018745 |
| ENSG00000106852 | LHX6 | -1.216642399 | 1.44E-10 |
| ENSG00000162613 | FUBP1 | -1.217427883 | 1.95E-33 |
| ENSG00000126790 | L3HYPDH | -1.218477222 | 3.91E-05 |
| ENSG00000136444 | RSAD1 | -1.219240837 | 3.29E-16 |
| ENSG00000184922 | FMNL1 | -1.220142099 | 1.40E-46 |
| ENSG00000159199 | ATP5MC1 | -1.220726491 | 9.99E-43 |
| ENSG00000146083 | RNF44 | -1.220888836 | 1.46E-34 |
| ENSG00000105851 | PIK3CG | -1.221437579 | 8.02E-13 |
| ENSG00000170632 | ARMC10 | -1.223262488 | 4.44E-35 |
| ENSG00000275004 | ZNF280B | -1.22476071 | 2.05E-06 |
| ENSG00000149054 | ZNF215 | -1.225378636 | 0.01152 |
| ENSG00000144815 | NXPE3 | -1.226844988 | 2.61E-22 |
| ENSG00000114767 | RRP9 | -1.22722106 | 1.39E-31 |
| ENSG00000048162 | NOP16 | -1.229499671 | 4.41E-44 |
| ENSG00000100365 | NCF4 | -1.229910989 | 4.51E-46 |
| ENSG00000164048 | ZNF589 | -1.230294625 | 2.25E-12 |
| ENSG00000125485 | DDX31 | -1.231627655 | 1.74E-27 |
| ENSG00000246082 | NUDT16L2P | -1.231958154 | 1.22E-12 |
| ENSG00000142765 | SYTL1 | -1.232366617 | 1.82E-05 |
| ENSG00000169105 | CHST14 | -1.233530331 | 0.000803 |
| ENSG00000085872 | CHERP | -1.234740882 | 1.25E-61 |
| ENSG00000184428 | TOP1MT | -1.234753966 | 3.21E-40 |
| ENSG00000140105 | WARS1 | -1.235009848 | 1.15E-69 |
| ENSG00000128604 | IRF5 | -1.235490222 | 0.0348 |
| ENSG00000135821 | GLUL | -1.235585081 | 3.02E-108 |
| ENSG00000132382 | MYBBP1A | -1.236176261 | 9.90E-49 |
| ENSG00000104331 | BPNT2 | -1.237240965 | 4.45E-32 |
| ENSG00000137393 | RNF144B | -1.237702163 | 8.33E-08 |
| ENSG00000100522 | GNPNAT1 | -1.238277161 | 2.99E-29 |
| ENSG00000110987 | BCL7A | -1.238356167 | 5.41E-22 |
| ENSG00000164933 | SLC25A32 | -1.240253954 | 0.000667 |
| ENSG00000144118 | RALB | -1.241479294 | 9.60E-73 |
| ENSG00000137404 | NRM | -1.24231509 | 2.36E-15 |
| ENSG00000174804 | FZD4 | -1.242876404 | 2.89E-19 |
| ENSG00000165879 | FRAT1 | -1.245608191 | 8.46E-09 |
| ENSG00000147010 | SH3KBP1 | -1.248569076 | 6.72E-16 |
| ENSG00000085832 | EPS15 | -1.248606272 | 1.03E-51 |
| ENSG00000101752 | MIB1 | -1.248695148 | 9.64E-32 |
| ENSG00000055483 | USP36 | -1.250068149 | 8.95E-49 |
| ENSG00000075826 | SEC31B | -1.250397684 | 1.10E-06 |
| ENSG00000103942 | HOMER2 | -1.251252712 | 8.48E-40 |
| ENSG00000151376 | ME3 | -1.252403997 | 2.50E-05 |
| ENSG00000186575 | NF2 | -1.253146616 | 1.99E-33 |
| ENSG00000133627 | ACTR3B | -1.254153396 | 3.06E-10 |
| ENSG00000136830 | NIBAN2 | -1.254228038 | 7.33E-68 |
| ENSG00000131323 | TRAF3 | -1.254808675 | 9.37E-33 |

| | | | |
|---|---|---|---|
| ENSG00000101158 | NELFCD | -1.254941605 | 3.38E-38 |
| ENSG00000188130 | MAPK12 | -1.255032425 | 1.14E-08 |
| ENSG00000154814 | OXNAD1 | -1.256175808 | 3.22E-28 |
| ENSG00000100336 | APOL4 | -1.25694407 | 5.38E-10 |
| ENSG00000277161 | PIGW | -1.257539879 | 1.78E-34 |
| ENSG00000115526 | CHST10 | -1.258861482 | 7.03E-23 |
| ENSG00000015475 | BID | -1.259511274 | 9.87E-26 |
| ENSG00000123600 | METTL8 | -1.260751071 | 5.25E-27 |
| ENSG00000188368 | PRR19 | -1.261535346 | 1.81E-07 |
| ENSG00000162729 | IGSF8 | -1.261949501 | 4.56E-07 |
| ENSG00000189144 | ZNF573 | -1.2627725 | 8.05E-07 |
| ENSG00000177732 | SOX12 | -1.262895165 | 2.46E-24 |
| ENSG00000291151 | NSUN5P1 | -1.263279583 | 0.000397 |
| ENSG00000107815 | TWNK | -1.264326593 | 2.87E-41 |
| ENSG00000196793 | ZNF239 | -1.264489536 | 6.32E-18 |
| ENSG00000198821 | CD247 | -1.264916739 | 0.000217 |
| ENSG00000186280 | KDM4D | -1.265442888 | 0.015122 |
| ENSG00000124787 | RPP40 | -1.266412144 | 1.08E-19 |
| ENSG00000290758 | gene_sourcehavana | -1.26661674 | 2.41E-05 |
| ENSG00000115504 | EHBP1 | -1.268229582 | 6.84E-28 |
| ENSG00000167588 | GPD1 | -1.268382061 | 0.011722 |
| ENSG00000291237 | SOD2 | -1.269002014 | 3.70E-51 |
| ENSG00000184986 | TMEM121 | -1.269845953 | 3.66E-05 |
| ENSG00000067113 | PLPP1 | -1.271064477 | 0.010774 |
| ENSG00000116096 | SPR | -1.271415216 | 8.81E-13 |
| ENSG00000110917 | MLEC | -1.272721109 | 6.44E-49 |
| ENSG00000006695 | COX10 | -1.272851375 | 1.14E-33 |
| ENSG00000105613 | MAST1 | -1.273146175 | 0.000572 |
| ENSG00000149970 | CNKSR2 | -1.27438829 | 2.53E-12 |
| ENSG00000183579 | ZNRF3 | -1.275071057 | 1.10E-16 |
| ENSG00000197183 | NOL4L | -1.276478586 | 2.18E-07 |
| ENSG00000272335 | gene_sourcehavana | -1.276787931 | 1.51E-07 |
| ENSG00000180353 | HCLS1 | -1.276996701 | 4.64E-16 |
| ENSG00000132635 | PCED1A | -1.277515002 | 1.41E-15 |
| ENSG00000156398 | SFXN2 | -1.277936217 | 0.000429 |
| ENSG00000226833 | PPP1CB-DT | -1.278906711 | 0.027146 |
| ENSG00000278530 | CHMP1B2P | -1.278932814 | 2.35E-13 |
| ENSG00000168297 | PXK | -1.279135422 | 2.19E-22 |
| ENSG00000147465 | STAR | -1.280893443 | 3.83E-05 |
| ENSG00000170417 | TMEM182 | -1.280914925 | 6.62E-05 |
| ENSG00000101003 | GINS1 | -1.281608297 | 1.46E-33 |
| ENSG00000127561 | SYNGR3 | -1.28529181 | 3.36E-15 |
| ENSG00000013016 | EHD3 | -1.285579358 | 5.77E-07 |
| ENSG00000116852 | KIF21B | -1.286009354 | 4.87E-15 |
| ENSG00000112218 | GPR63 | -1.287651741 | 1.37E-05 |
| ENSG00000065057 | NTHL1 | -1.287654771 | 1.73E-13 |
| ENSG00000213339 | QTRT1 | -1.28766942 | 1.62E-05 |
| ENSG00000114529 | C3orf52 | -1.288010929 | 0.000958 |
| ENSG00000103495 | MAZ | -1.289427647 | 1.21E-52 |

| | | | |
|---|---|---|---|
| ENSG00000156521 | TYSND1 | -1.290397232 | 1.39E-17 |
| ENSG00000124019 | FAM124B | -1.291324758 | 7.49E-28 |
| ENSG00000165097 | KDM1B | -1.295061248 | 2.62E-47 |
| ENSG00000119900 | OGFRL1 | -1.296219784 | 1.24E-19 |
| ENSG00000140043 | PTGR2 | -1.297295936 | 9.80E-05 |
| ENSG00000119946 | CNNM1 | -1.298494342 | 1.91E-20 |
| ENSG00000153936 | HS2ST1 | -1.300194616 | 5.27E-34 |
| ENSG00000116649 | SRM | -1.301772385 | 1.58E-37 |
| ENSG00000198838 | RYR3 | -1.302877022 | 3.03E-21 |
| ENSG00000167543 | TP53I13 | -1.303094884 | 7.15E-09 |
| ENSG00000096401 | CDC5L | -1.303470427 | 3.08E-11 |
| ENSG00000177192 | PUS1 | -1.304133522 | 2.01E-35 |
| ENSG00000134461 | ANKRD16 | -1.305057585 | 2.61E-10 |
| ENSG00000004399 | PLXND1 | -1.305619588 | 5.75E-11 |
| ENSG00000291071 | FAM86JP | -1.306008309 | 0.000124 |
| ENSG00000185404 | SP140L | -1.30740637 | 5.63E-10 |
| ENSG00000179627 | ZBTB42 | -1.308295082 | 3.95E-05 |
| ENSG00000134138 | MEIS2 | -1.308443322 | 1.84E-22 |
| ENSG00000155313 | USP25 | -1.308456109 | 2.23E-61 |
| ENSG00000182670 | TTC3 | -1.310054226 | 1.17E-65 |
| ENSG00000105426 | PTPRS | -1.310416251 | 0.000419 |
| ENSG00000215021 | PHB2 | -1.311952635 | 1.87E-13 |
| ENSG00000149948 | HMGA2 | -1.311995572 | 1.84E-07 |
| ENSG00000197457 | STMN3 | -1.312370444 | 1.14E-16 |
| ENSG00000226742 | HSBP1L1 | -1.31282234 | 9.16E-16 |
| ENSG00000072506 | HSD17B10 | -1.31309446 | 2.65E-70 |
| ENSG00000148814 | LRRC27 | -1.313494197 | 0.000189 |
| ENSG00000198298 | ZNF485 | -1.314015014 | 7.05E-13 |
| ENSG00000279348 | gene_sourcehavana | -1.315249831 | 0.000605 |
| ENSG00000181827 | RFX7 | -1.31968084 | 1.10E-34 |
| ENSG00000237943 | PRKCQ-AS1 | -1.319785542 | 3.22E-24 |
| ENSG00000261167 | gene_sourcehavana | -1.31997857 | 1.63E-06 |
| ENSG00000153107 | ANAPC1 | -1.32125786 | 1.62E-41 |
| ENSG00000235381 | gene_sourcehavana | -1.321693361 | 0.003961 |
| ENSG00000164120 | HPGD | -1.322999546 | 1.22E-06 |
| ENSG00000288947 | gene_sourcehavana_ | -1.323133035 | 0.004512 |
| ENSG00000183401 | CCDC159 | -1.326441295 | 4.91E-11 |
| ENSG00000237669 | gene_sourcehavana | -1.326778729 | 0.043615 |
| ENSG00000159842 | ABR | -1.327776331 | 7.45E-42 |
| ENSG00000274265 | gene_sourcehavana | -1.328451807 | 0.005192 |
| ENSG00000143627 | PKLR | -1.328940656 | 3.41E-09 |
| ENSG00000165215 | CLDN3 | -1.330864203 | 5.59E-06 |
| ENSG00000102967 | DHODH | -1.331778208 | 5.75E-20 |
| ENSG00000259802 | MARCHF6-DT | -1.333072192 | 0.018705 |
| ENSG00000198894 | CIPC | -1.334817353 | 4.14E-21 |
| ENSG00000133739 | LRRCC1 | -1.334819917 | 3.36E-45 |
| ENSG00000105983 | LMBR1 | -1.334837568 | 1.36E-32 |
| ENSG00000138031 | ADCY3 | -1.335459412 | 1.10E-42 |
| ENSG00000064102 | INTS13 | -1.335517151 | 4.60E-71 |

| | | | |
|---|---|---|---|
| ENSG00000196810 | CTBP1-DT | -1.336908968 | 3.52E-19 |
| ENSG00000160214 | RRP1 | -1.337769779 | 2.37E-62 |
| ENSG00000242173 | ARHGDIG | -1.340036255 | 0.002632 |
| ENSG00000228175 | GEMIN8P4 | -1.340490665 | 0.011635 |
| ENSG00000291178 | gene_sourcehavana | -1.340903319 | 4.98E-09 |
| ENSG00000182796 | TMEM198B | -1.340962104 | 7.53E-12 |
| ENSG00000214063 | TSPAN4 | -1.341005388 | 1.97E-16 |
| ENSG00000103490 | PYCARD | -1.341018438 | 1.26E-05 |
| ENSG00000164953 | TMEM67 | -1.345740429 | 1.42E-10 |
| ENSG00000015285 | WAS | -1.346991244 | 4.31E-20 |
| ENSG00000134602 | STK26 | -1.347392834 | 9.37E-38 |
| ENSG00000163602 | RYBP | -1.3473968 | 2.23E-60 |
| ENSG00000232648 | ING2-DT | -1.347721251 | 0.010276 |
| ENSG00000187630 | DHRS4L2 | -1.34791478 | 3.25E-22 |
| ENSG00000073536 | NLE1 | -1.348913626 | 1.93E-23 |
| ENSG00000142197 | DOP1B | -1.349253216 | 1.96E-16 |
| ENSG00000097096 | SYDE2 | -1.350218558 | 9.31E-08 |
| ENSG00000185482 | STAC3 | -1.350521691 | 0.000115 |
| ENSG00000059588 | TARBP1 | -1.351052914 | 1.54E-36 |
| ENSG00000236810 | ELOA-AS1 | -1.351099102 | 3.18E-06 |
| ENSG00000284753 | EEF1AKMT4 | -1.352438668 | 7.09E-07 |
| ENSG00000233058 | ATP13A3-DT | -1.352859921 | 4.30E-14 |
| ENSG00000117479 | SLC19A2 | -1.354704975 | 8.86E-13 |
| ENSG00000109861 | CTSC | -1.356181586 | 8.40E-40 |
| ENSG00000165424 | ZCCHC24 | -1.356701866 | 2.38E-07 |
| ENSG00000284691 | gene_sourceensembl | -1.359042114 | 2.19E-11 |
| ENSG00000169598 | DFFB | -1.359228997 | 7.42E-18 |
| ENSG00000290374 | FAM86EP | -1.360543054 | 2.58E-05 |
| ENSG00000111110 | PPM1H | -1.360812114 | 1.04E-30 |
| ENSG00000146776 | ATXN7L1 | -1.361287399 | 8.29E-28 |
| ENSG00000247095 | MIR210HG | -1.361530137 | 0.000488 |
| ENSG00000109084 | TMEM97 | -1.361662274 | 1.49E-08 |
| ENSG00000166166 | TRMT61A | -1.363258258 | 1.13E-32 |
| ENSG00000235989 | MORC2-AS1 | -1.363375344 | 0.002277 |
| ENSG00000290831 | NSUN5P2 | -1.363378269 | 0.036401 |
| ENSG00000129480 | DTD2 | -1.363817383 | 1.66E-19 |
| ENSG00000142347 | MYO1F | -1.363957587 | 1.96E-06 |
| ENSG00000183242 | WT1-AS | -1.364357323 | 1.52E-07 |
| ENSG00000204946 | ZNF783 | -1.365626701 | 3.16E-23 |
| ENSG00000243056 | EIF4EBP3 | -1.366262508 | 0.013232 |
| ENSG00000187840 | EIF4EBP1 | -1.366844394 | 1.18E-45 |
| ENSG00000165752 | STK32C | -1.366875956 | 5.09E-08 |
| ENSG00000122490 | SLC66A2 | -1.367090569 | 4.71E-45 |
| ENSG00000206535 | LNP1 | -1.367170924 | 5.04E-05 |
| ENSG00000185499 | MUC1 | -1.368311244 | 3.22E-08 |
| ENSG00000135372 | NAT10 | -1.370411387 | 2.94E-47 |
| ENSG00000290596 | gene_sourcehavana | -1.372080264 | 0.000165 |
| ENSG00000113749 | HRH2 | -1.372162403 | 9.45E-17 |
| ENSG00000128274 | A4GALT | -1.372186901 | 6.66E-37 |

| | | | |
|---|---|---|---|
| ENSG00000167566 | NCKAP5L | -1.373356879 | 0.000168 |
| ENSG00000102230 | PCYT1B | -1.373368414 | 9.85E-23 |
| ENSG00000159753 | CARMIL2 | -1.374346487 | 3.41E-09 |
| ENSG00000259248 | USP3-AS1 | -1.377754442 | 0.015706 |
| ENSG00000130751 | NPAS1 | -1.37817429 | 2.79E-06 |
| ENSG00000145819 | ARHGAP26 | -1.379378118 | 0.000346 |
| ENSG00000161956 | SENP3 | -1.379865164 | 3.20E-17 |
| ENSG00000118965 | WDR35 | -1.381049621 | 9.83E-31 |
| ENSG00000276529 | gene_sourcehavana | -1.381858181 | 0.000222 |
| ENSG00000170011 | MYRIP | -1.382757214 | 0.013854 |
| ENSG00000197442 | MAP3K5 | -1.38314524 | 9.28E-17 |
| ENSG00000083807 | SLC27A5 | -1.385047541 | 5.32E-12 |
| ENSG00000127364 | TAS2R4 | -1.385366926 | 0.038469 |
| ENSG00000107331 | ABCA2 | -1.385691558 | 4.72E-12 |
| ENSG00000229124 | VIM-AS1 | -1.385784409 | 1.08E-08 |
| ENSG00000118707 | TGIF2 | -1.389519682 | 1.58E-15 |
| ENSG00000253154 | gene_sourcehavana | -1.39073433 | 6.10E-11 |
| ENSG00000186026 | ZNF284 | -1.39185097 | 0.003232 |
| ENSG00000090376 | IRAK3 | -1.392565129 | 1.87E-40 |
| ENSG00000132591 | ERAL1 | -1.392681797 | 2.11E-39 |
| ENSG00000111696 | NT5DC3 | -1.393485426 | 3.76E-22 |
| ENSG00000161267 | BDH1 | -1.39371675 | 2.30E-39 |
| ENSG00000158457 | TSPAN33 | -1.396338865 | 0.000782 |
| ENSG00000161010 | MRNIP | -1.396980545 | 6.43E-23 |
| ENSG00000177674 | AGTRAP | -1.397808377 | 2.62E-30 |
| ENSG00000223960 | CHROMR | -1.398823654 | 4.29E-15 |
| ENSG00000183605 | SFXN4 | -1.39915065 | 4.10E-42 |
| ENSG00000137965 | IFI44 | -1.399240082 | 0.015927 |
| ENSG00000154930 | ACSS1 | -1.400997469 | 2.56E-08 |
| ENSG00000188211 | NCR3LG1 | -1.404481528 | 2.56E-52 |
| ENSG00000142657 | PGD | -1.406516295 | 2.89E-61 |
| ENSG00000117877 | POLR1G | -1.407120367 | 1.21E-37 |
| ENSG00000159216 | RUNX1 | -1.408307237 | 1.02E-47 |
| ENSG00000166896 | ATP23 | -1.408491314 | 3.32E-21 |
| ENSG00000115266 | APC2 | -1.409760073 | 1.60E-16 |
| ENSG00000153790 | C7orf31 | -1.410052817 | 0.001425 |
| ENSG00000197448 | GSTK1 | -1.410662658 | 4.44E-08 |
| ENSG00000065308 | TRAM2 | -1.412301634 | 3.07E-45 |
| ENSG00000115255 | REEP6 | -1.412423303 | 2.97E-12 |
| ENSG00000095485 | CWF19L1 | -1.414439813 | 6.57E-44 |
| ENSG00000153885 | KCTD15 | -1.416859698 | 2.89E-20 |
| ENSG00000130590 | SAMD10 | -1.416950916 | 2.41E-13 |
| ENSG00000148204 | CRB2 | -1.417180948 | 1.79E-10 |
| ENSG00000261373 | VPS9D1-AS1 | -1.417904394 | 7.40E-25 |
| ENSG00000212747 | RTL8B | -1.41799235 | 7.80E-23 |
| ENSG00000198865 | CCDC152 | -1.419989142 | 1.01E-09 |
| ENSG00000261294 | gene_sourcehavana | -1.423115802 | 3.95E-07 |
| ENSG00000131188 | PRR7 | -1.425425348 | 1.09E-06 |
| ENSG00000129103 | SUMF2 | -1.426726335 | 1.34E-08 |

| | | | |
|---|---|---|---|
| ENSG00000101194 | SLC17A9 | -1.426919771 | 2.58E-38 |
| ENSG00000173200 | PARP15 | -1.430193341 | 4.44E-31 |
| ENSG00000133943 | DGLUCY | -1.430754129 | 3.12E-36 |
| ENSG00000101236 | RNF24 | -1.431427193 | 4.09E-45 |
| ENSG00000159423 | ALDH4A1 | -1.431546615 | 3.25E-56 |
| ENSG00000100583 | SAMD15 | -1.432482351 | 0.004183 |
| ENSG00000156471 | PTDSS1 | -1.433208764 | 3.08E-49 |
| ENSG00000204267 | TAP2 | -1.433274005 | 3.56E-18 |
| ENSG00000110651 | CD81 | -1.43471007 | 1.72E-39 |
| ENSG00000128694 | OSGEPL1 | -1.435286117 | 1.64E-13 |
| ENSG00000119599 | DCAF4 | -1.437131472 | 3.23E-21 |
| ENSG00000272142 | LYRM4-AS1 | -1.437135744 | 0.003904 |
| ENSG00000125901 | MRPS26 | -1.438115386 | 1.33E-34 |
| ENSG00000117862 | TXNDC12 | -1.441046866 | 3.37E-06 |
| ENSG00000135245 | HILPDA | -1.446306734 | 5.18E-08 |
| ENSG00000184937 | WT1 | -1.446689477 | 1.49E-05 |
| ENSG00000224635 | gene_sourcehavana_ | -1.448163932 | 0.048337 |
| ENSG00000272325 | NUDT3 | -1.448221853 | 4.53E-36 |
| ENSG00000020256 | ZFP64 | -1.448388074 | 6.38E-26 |
| ENSG00000231890 | DARS1-AS1 | -1.448886423 | 0.013692 |
| ENSG00000155893 | PXYLP1 | -1.449045284 | 0.000309 |
| ENSG00000171492 | LRRC8D | -1.449925614 | 9.58E-34 |
| ENSG00000112041 | TULP1 | -1.451246027 | 0.031841 |
| ENSG00000198464 | ZNF480 | -1.451857173 | 1.62E-41 |
| ENSG00000027697 | IFNGR1 | -1.452099016 | 5.20E-29 |
| ENSG00000164292 | RHOBTB3 | -1.452592846 | 1.28E-55 |
| ENSG00000176919 | C8G | -1.453073345 | 0.030171 |
| ENSG00000109099 | PMP22 | -1.453320224 | 2.01E-62 |
| ENSG00000083123 | BCKDHB | -1.45373676 | 5.28E-24 |
| ENSG00000134278 | SPIRE1 | -1.455179003 | 3.30E-16 |
| ENSG00000075618 | FSCN1 | -1.457441844 | 2.09E-61 |
| ENSG00000259953 | LINC02977 | -1.457576189 | 1.48E-10 |
| ENSG00000106415 | GLCCI1 | -1.458166028 | 1.09E-38 |
| ENSG00000103653 | CSK | -1.458305916 | 2.05E-65 |
| ENSG00000100109 | TFIP11 | -1.458323133 | 2.41E-107 |
| ENSG00000176124 | DLEU1 | -1.458636789 | 1.72E-08 |
| ENSG00000141384 | TAF4B | -1.460312841 | 2.19E-11 |
| ENSG00000174721 | FGFBP3 | -1.460902272 | 2.61E-05 |
| ENSG00000184361 | SPATA32 | -1.461757157 | 6.27E-07 |
| ENSG00000156697 | UTP14A | -1.463603615 | 6.59E-11 |
| ENSG00000176834 | VSIG10 | -1.464411462 | 0.008945 |
| ENSG00000067992 | PDK3 | -1.466153544 | 2.98E-26 |
| ENSG00000168792 | ABHD15 | -1.469243268 | 7.53E-09 |
| ENSG00000246582 | TNFRSF10A-DT | -1.469888613 | 1.89E-12 |
| ENSG00000168026 | TTC21A | -1.470000869 | 0.008456 |
| ENSG00000151065 | DCP1B | -1.470047165 | 8.29E-25 |
| ENSG00000166557 | TMED3 | -1.470157896 | 1.09E-60 |
| ENSG00000164402 | SEPTIN8 | -1.471015232 | 6.54E-58 |
| ENSG00000110400 | NECTIN1 | -1.472436814 | 1.75E-78 |

| | | | |
|---|---|---|---|
| ENSG00000110876 | SELPLG | -1.472818168 | 1.68E-08 |
| ENSG00000133983 | COX16 | -1.473280124 | 0.001778 |
| ENSG00000065485 | PDIA5 | -1.474701425 | 1.37E-26 |
| ENSG00000163006 | CCDC138 | -1.47494092 | 5.34E-20 |
| ENSG00000243468 | INGX | -1.475916701 | 0.015938 |
| ENSG00000137513 | NARS2 | -1.47682353 | 3.10E-32 |
| ENSG00000214954 | LRRC69 | -1.476910819 | 0.0068 |
| ENSG00000106080 | FKBP14 | -1.477249598 | 1.43E-17 |
| ENSG00000076555 | ACACB | -1.479864571 | 3.48E-12 |
| ENSG00000129295 | DNAAF11 | -1.480634091 | 5.12E-10 |
| ENSG00000164300 | SERINC5 | -1.482293884 | 6.04E-74 |
| ENSG00000289548 | gene_sourcehavana_ | -1.482359628 | 4.82E-21 |
| ENSG00000105552 | BCAT2 | -1.48598095 | 8.17E-35 |
| ENSG00000123104 | ITPR2 | -1.485984813 | 1.00E-59 |
| ENSG00000145685 | LHFPL2 | -1.488145201 | 3.56E-53 |
| ENSG00000167272 | POP5 | -1.488642199 | 1.68E-25 |
| ENSG00000169570 | DTWD2 | -1.489217624 | 8.30E-12 |
| ENSG00000145911 | N4BP3 | -1.489267531 | 2.74E-09 |
| ENSG00000135338 | LCA5 | -1.490715447 | 2.04E-06 |
| ENSG00000164039 | BDH2 | -1.49139397 | 7.71E-08 |
| ENSG00000101187 | SLCO4A1 | -1.493736363 | 1.98E-29 |
| ENSG00000187024 | PTRH1 | -1.493783256 | 1.39E-10 |
| ENSG00000082153 | BZW1 | -1.493826857 | 1.78E-110 |
| ENSG00000183048 | SLC25A10 | -1.494098798 | 1.33E-13 |
| ENSG00000172197 | MBOAT1 | -1.495075461 | 2.02E-30 |
| ENSG00000152620 | NADK2 | -1.495440265 | 4.23E-57 |
| ENSG00000226942 | IL9RP3 | -1.496951605 | 0.001648 |
| ENSG00000123179 | EBPL | -1.49727001 | 3.03E-27 |
| ENSG00000072818 | ACAP1 | -1.497755425 | 1.50E-52 |
| ENSG00000137699 | TRIM29 | -1.497827118 | 6.28E-57 |
| ENSG00000069998 | HDHD5 | -1.497991378 | 1.57E-21 |
| ENSG00000010318 | PHF7 | -1.499889181 | 2.91E-46 |
| ENSG00000160973 | FOXH1 | -1.501358898 | 0.000322 |
| ENSG00000132768 | DPH2 | -1.502433895 | 1.14E-34 |
| ENSG00000174669 | SLC29A2 | -1.502546405 | 1.17E-28 |
| ENSG00000108961 | RANGRF | -1.503392586 | 2.55E-38 |
| ENSG00000287151 | C2orf27A | -1.504092307 | 0.003654 |
| ENSG00000183454 | GRIN2A | -1.504333673 | 2.22E-06 |
| ENSG00000172349 | IL16 | -1.506661871 | 1.62E-05 |
| ENSG00000134285 | FKBP11 | -1.509474483 | 4.25E-19 |
| ENSG00000010818 | HIVEP2 | -1.510795142 | 0.001414 |
| ENSG00000260230 | FRRS1L | -1.510810541 | 4.87E-24 |
| ENSG00000104635 | SLC39A14 | -1.51104396 | 1.42E-48 |
| ENSG00000126947 | ARMCX1 | -1.513170104 | 1.16E-49 |
| ENSG00000186235 | LINC02610 | -1.513188052 | 6.83E-10 |
| ENSG00000047648 | ARHGAP6 | -1.514526372 | 1.96E-12 |
| ENSG00000124813 | RUNX2 | -1.514964292 | 8.29E-19 |
| ENSG00000173898 | SPTBN2 | -1.521063087 | 4.56E-48 |
| ENSG00000109736 | MFSD10 | -1.521631813 | 4.88E-29 |

| | | | |
|---|---|---|---|
| ENSG00000245213 | GALNT7-DT | -1.521687226 | 0.041946 |
| ENSG00000105963 | ADAP1 | -1.522605777 | 0.04415 |
| ENSG00000269906 | gene_sourcehavana | -1.524410628 | 0.019384 |
| ENSG00000119922 | IFIT2 | -1.525631185 | 5.17E-09 |
| ENSG00000143847 | PPFIA4 | -1.526791919 | 6.80E-06 |
| ENSG00000145604 | SKP2 | -1.528054086 | 7.37E-12 |
| ENSG00000008394 | MGST1 | -1.528734754 | 9.76E-44 |
| ENSG00000107833 | NPM3 | -1.530891754 | 1.63E-33 |
| ENSG00000110104 | CCDC86 | -1.531151441 | 1.95E-43 |
| ENSG00000181619 | GPR135 | -1.532317905 | 0.005851 |
| ENSG00000267475 | NUDT19-DT | -1.533088451 | 5.54E-11 |
| ENSG00000188015 | S100A3 | -1.533438711 | 0.000946 |
| ENSG00000279970 | gene_sourcehavana | -1.535107256 | 0.000356 |
| ENSG00000172137 | CALB2 | -1.535201731 | 0.030223 |
| ENSG00000272604 | EFCAB10-AS1 | -1.538462158 | 0.019234 |
| ENSG00000121690 | DEPDC7 | -1.5411631 | 8.70E-17 |
| ENSG00000247626 | MARS2 | -1.542266292 | 3.43E-23 |
| ENSG00000115935 | WIPF1 | -1.543358647 | 4.36E-18 |
| ENSG00000115956 | PLEK | -1.543933035 | 0.000114 |
| ENSG00000172731 | LRRC20 | -1.544043353 | 1.80E-11 |
| ENSG00000143498 | TAF1A | -1.549322065 | 4.98E-14 |
| ENSG00000135439 | AGAP2 | -1.550090213 | 4.57E-36 |
| ENSG00000196743 | GM2A | -1.550106931 | 7.51E-35 |
| ENSG00000136531 | SCN2A | -1.551053926 | 1.61E-21 |
| ENSG00000269210 | gene_sourcehavana | -1.552970932 | 3.62E-07 |
| ENSG00000171067 | C11orf24 | -1.556497968 | 2.48E-34 |
| ENSG00000226419 | SLC16A1-AS1 | -1.556965514 | 1.46E-12 |
| ENSG00000174749 | FAM241A | -1.557051096 | 6.61E-27 |
| ENSG00000163864 | NMNAT3 | -1.557100159 | 4.57E-30 |
| ENSG00000087053 | MTMR2 | -1.558241624 | 2.51E-64 |
| ENSG00000014919 | COX15 | -1.560096687 | 8.27E-34 |
| ENSG00000102007 | PLP2 | -1.560671065 | 1.59E-51 |
| ENSG00000146281 | PM20D2 | -1.563008116 | 6.02E-23 |
| ENSG00000136997 | MYC | -1.56660327 | 8.42E-66 |
| ENSG00000101384 | JAG1 | -1.566668893 | 4.61E-28 |
| ENSG00000226887 | ERVMER34-1 | -1.568007425 | 1.28E-15 |
| ENSG00000178921 | PFAS | -1.568009715 | 1.44E-11 |
| ENSG00000163513 | TGFBR2 | -1.568509142 | 3.59E-29 |
| ENSG00000115204 | MPV17 | -1.568617882 | 2.14E-06 |
| ENSG00000100504 | PYGL | -1.570851629 | 7.70E-53 |
| ENSG00000291177 | gene_sourcehavana | -1.571579201 | 0.000617 |
| ENSG00000145248 | SLC10A4 | -1.573677807 | 1.40E-38 |
| ENSG00000136059 | VILL | -1.574196893 | 3.47E-15 |
| ENSG00000163093 | BBS5 | -1.576719847 | 6.04E-07 |
| ENSG00000167925 | GHDC | -1.577544934 | 6.62E-18 |
| ENSG00000146072 | TNFRSF21 | -1.580004288 | 2.24E-05 |
| ENSG00000075035 | WSCD2 | -1.580746959 | 5.61E-32 |
| ENSG00000257698 | GIHCG | -1.58153943 | 2.70E-29 |
| ENSG00000131187 | F12 | -1.582329951 | 0.000123 |

| | | | |
|---|---|---|---|
| ENSG00000116337 | AMPD2 | -1.585614846 | 1.58E-38 |
| ENSG00000184949 | FAM227A | -1.585775996 | 0.002195 |
| ENSG00000158517 | NCF1 | -1.58733297 | 0.033049 |
| ENSG00000163576 | EFHB | -1.591209461 | 0.000141 |
| ENSG00000089041 | P2RX7 | -1.592252048 | 8.29E-06 |
| ENSG00000289089 | gene_sourcehavana_ | -1.593965303 | 0.00935 |
| ENSG00000203709 | MIR29B2CHG | -1.594951147 | 3.03E-07 |
| ENSG00000172315 | TP53RK | -1.594993764 | 1.48E-38 |
| ENSG00000114631 | PODXL2 | -1.596221494 | 5.00E-51 |
| ENSG00000196372 | ASB13 | -1.596795046 | 2.19E-21 |
| ENSG00000271754 | gene_sourcehavana | -1.598726225 | 0.02148 |
| ENSG00000197905 | TEAD4 | -1.599321388 | 1.27E-20 |
| ENSG00000266094 | RASSF5 | -1.600972881 | 1.53E-54 |
| ENSG00000232677 | LINC00665 | -1.606248977 | 1.06E-09 |
| ENSG00000262468 | LINC01569 | -1.607100221 | 4.12E-08 |
| ENSG00000239789 | MRPS17 | -1.60804099 | 4.09E-12 |
| ENSG00000139737 | SLAIN1 | -1.609009349 | 2.64E-18 |
| ENSG00000221955 | SLC12A8 | -1.60975414 | 5.51E-06 |
| ENSG00000287055 | gene_sourcehavana_ | -1.612964143 | 0.001626 |
| ENSG00000133256 | PDE6B | -1.615435738 | 1.33E-15 |
| ENSG00000065183 | WDR3 | -1.61565954 | 3.27E-58 |
| ENSG00000247796 | MOCS2-DT | -1.616380268 | 1.48E-09 |
| ENSG00000267757 | EML2-AS1 | -1.617759292 | 5.31E-06 |
| ENSG00000116128 | BCL9 | -1.620671204 | 1.84E-20 |
| ENSG00000106948 | AKNA | -1.620821736 | 0.012757 |
| ENSG00000111145 | ELK3 | -1.622906627 | 6.34E-19 |
| ENSG00000151131 | NOPCHAP1 | -1.624237266 | 2.48E-20 |
| ENSG00000248538 | PPP1R3B-DT | -1.625220527 | 0.000572 |
| ENSG00000130768 | SMPDL3B | -1.625296634 | 1.04E-10 |
| ENSG00000254726 | MEX3A | -1.626280219 | 0.010905 |
| ENSG00000246627 | CACNA1C-AS1 | -1.629331651 | 1.50E-05 |
| ENSG00000048740 | CELF2 | -1.631849183 | 3.78E-64 |
| ENSG00000135966 | TGFBRAP1 | -1.632716993 | 3.62E-66 |
| ENSG00000172264 | MACROD2 | -1.636677676 | 0.003042 |
| ENSG00000256525 | POLG2 | -1.639641245 | 7.80E-41 |
| ENSG00000184979 | USP18 | -1.640527488 | 4.10E-07 |
| ENSG00000175164 | ABO | -1.643321149 | 6.30E-15 |
| ENSG00000260196 | gene_sourcehavana | -1.643678486 | 7.58E-25 |
| ENSG00000285761 | gene_sourcehavana | -1.645115232 | 0.007908 |
| ENSG00000136159 | NUDT15 | -1.648192643 | 4.05E-23 |
| ENSG00000197766 | CFD | -1.648533756 | 4.19E-11 |
| ENSG00000290994 | FAM86C2P | -1.652956657 | 3.25E-05 |
| ENSG00000145220 | LYAR | -1.654504837 | 4.17E-75 |
| ENSG00000188343 | CIBAR1 | -1.65529136 | 1.40E-12 |
| ENSG00000173262 | SLC2A14 | -1.657630021 | 0.032666 |
| ENSG00000151458 | ANKRD50 | -1.659164542 | 1.58E-60 |
| ENSG00000144749 | LRIG1 | -1.660494944 | 8.43E-06 |
| ENSG00000082175 | PGR | -1.667031973 | 3.16E-78 |
| ENSG00000196418 | ZNF124 | -1.667200299 | 1.52E-19 |

| | | | |
|---|---|---|---|
| ENSG00000100368 | CSF2RB | -1.669520044 | 6.54E-139 |
| ENSG00000111348 | ARHGDIB | -1.670808481 | 5.73E-74 |
| ENSG00000082516 | GEMIN5 | -1.671754871 | 2.38E-53 |
| ENSG00000099284 | MACROH2A2 | -1.672590639 | 0.019951 |
| ENSG00000160471 | COX6B2 | -1.67290432 | 1.95E-09 |
| ENSG00000175711 | B3GNTL1 | -1.672929511 | 4.31E-12 |
| ENSG00000159363 | ATP13A2 | -1.673066641 | 8.82E-67 |
| ENSG00000159674 | SPON2 | -1.673640334 | 1.95E-10 |
| ENSG00000116918 | TSNAX | -1.675511438 | 8.90E-78 |
| ENSG00000244513 | EOGT-DT | -1.675518641 | 0.004151 |
| ENSG00000167202 | TBC1D2B | -1.67570837 | 2.62E-16 |
| ENSG00000181458 | TMEM45A | -1.676605423 | 0.000675 |
| ENSG00000188807 | TMEM201 | -1.678951451 | 4.12E-47 |
| ENSG00000257718 | CPNE8-AS1 | -1.681935917 | 0.028746 |
| ENSG00000102743 | SLC25A15 | -1.682834242 | 1.44E-55 |
| ENSG00000107186 | MPDZ | -1.684490687 | 8.24E-38 |
| ENSG00000143674 | MAP3K21 | -1.684686903 | 2.50E-46 |
| ENSG00000101079 | NDRG3 | -1.685059545 | 6.82E-48 |
| ENSG00000143061 | IGSF3 | -1.689308997 | 3.15E-06 |
| ENSG00000271601 | LIX1L | -1.693209983 | 9.92E-53 |
| ENSG00000151623 | NR3C2 | -1.696462785 | 0.047949 |
| ENSG00000228649 | SNHG26 | -1.697798693 | 9.53E-18 |
| ENSG00000109906 | ZBTB16 | -1.697844251 | 3.14E-20 |
| ENSG00000248578 | NPM1P21 | -1.699311844 | 0.018809 |
| ENSG00000133935 | ERG28 | -1.70515392 | 1.93E-50 |
| ENSG00000291049 | FAM86B3P | -1.706584772 | 2.15E-16 |
| ENSG00000169239 | CA5B | -1.707275441 | 5.31E-10 |
| ENSG00000153574 | RPIA | -1.708082864 | 1.90E-47 |
| ENSG00000225439 | BOLA3-DT | -1.709780632 | 3.68E-31 |
| ENSG00000079150 | FKBP7 | -1.712068184 | 7.82E-05 |
| ENSG00000291220 | gene_sourceensembl | -1.714623446 | 0.045593 |
| ENSG00000289372 | gene_sourcehavana_ | -1.715888317 | 0.001054 |
| ENSG00000119865 | CNRIP1 | -1.715985098 | 9.47E-54 |
| ENSG00000278291 | gene_sourcehavana | -1.719800367 | 0.017706 |
| ENSG00000272395 | IFNL4 | -1.721380166 | 0.001162 |
| ENSG00000168778 | TCTN2 | -1.721912106 | 4.91E-19 |
| ENSG00000249639 | gene_sourcehavana | -1.722817481 | 0.032293 |
| ENSG00000065675 | PRKCQ | -1.722983271 | 7.18E-51 |
| ENSG00000288046 | gene_sourcehavana_ | -1.723474753 | 9.76E-09 |
| ENSG00000134202 | GSTM3 | -1.724728498 | 9.04E-16 |
| ENSG00000187049 | TMEM216 | -1.727902587 | 2.50E-23 |
| ENSG00000241743 | XACT | -1.732660974 | 3.11E-32 |
| ENSG00000254854 | NECTIN1-DT | -1.73336576 | 0.039876 |
| ENSG00000116771 | AGMAT | -1.733504211 | 7.40E-19 |
| ENSG00000105926 | PALS2 | -1.733773601 | 7.50E-37 |
| ENSG00000287672 | gene_sourcehavana_ | -1.736276664 | 0.000125 |
| ENSG00000237686 | SCIRT | -1.738134169 | 0.00311 |
| ENSG00000106484 | MEST | -1.739961562 | 1.48E-17 |
| ENSG00000076351 | SLC46A1 | -1.741836022 | 1.23E-07 |

| | | | |
|---|---|---|---|
| ENSG00000101166 | PRELID3B | -1.742764404 | 3.65E-99 |
| ENSG00000140470 | ADAMTS17 | -1.743506744 | 0.042589 |
| ENSG00000130038 | CRACR2A | -1.743561698 | 4.10E-42 |
| ENSG00000180448 | ARHGAP45 | -1.744210076 | 1.52E-68 |
| ENSG00000137098 | SPAG8 | -1.744627924 | 0.035698 |
| ENSG00000157600 | TMEM164 | -1.74811269 | 9.71E-92 |
| ENSG00000089558 | KCNH4 | -1.751040649 | 0.012688 |
| ENSG00000103485 | QPRT | -1.751410013 | 6.01E-44 |
| ENSG00000065054 | NHERF2 | -1.752728661 | 1.21E-08 |
| ENSG00000186918 | ZNF395 | -1.755248266 | 7.27E-12 |
| ENSG00000160161 | CILP2 | -1.755725672 | 8.25E-23 |
| ENSG00000285600 | gene_sourcehavana | -1.756063021 | 0.000239 |
| ENSG00000243701 | DUBR | -1.756724227 | 3.12E-06 |
| ENSG00000184785 | SMIM10 | -1.757564882 | 9.32E-74 |
| ENSG00000124134 | KCNS1 | -1.758148409 | 2.94E-55 |
| ENSG00000011376 | LARS2 | -1.758323114 | 4.38E-61 |
| ENSG00000151364 | KCTD14 | -1.759923008 | 0.014656 |
| ENSG00000167600 | CYP2S1 | -1.76072332 | 1.78E-08 |
| ENSG00000198890 | PRMT6 | -1.764165615 | 1.57E-64 |
| ENSG00000115257 | PCSK4 | -1.769346594 | 4.11E-10 |
| ENSG00000168421 | RHOH | -1.769481141 | 5.08E-41 |
| ENSG00000204745 | gene_sourcehavana | -1.769669664 | 9.27E-10 |
| ENSG00000099365 | STX1B | -1.77193551 | 5.26E-05 |
| ENSG00000279307 | NDUFAF2P1 | -1.775874125 | 0.002216 |
| ENSG00000152558 | TMEM123 | -1.776211835 | 2.15E-10 |
| ENSG00000162430 | SELENON | -1.781688946 | 3.25E-40 |
| ENSG00000182511 | FES | -1.787398368 | 2.15E-38 |
| ENSG00000226777 | FAM30A | -1.788006188 | 9.77E-07 |
| ENSG00000126368 | NR1D1 | -1.790139415 | 1.09E-16 |
| ENSG00000144560 | VGLL4 | -1.790360382 | 1.54E-07 |
| ENSG00000288973 | gene_sourcehavana_ | -1.79213683 | 6.87E-77 |
| ENSG00000119714 | GPR68 | -1.793815434 | 0.032179 |
| ENSG00000104312 | RIPK2 | -1.802420528 | 2.34E-43 |
| ENSG00000179241 | LDLRAD3 | -1.807273169 | 3.94E-29 |
| ENSG00000284237 | LINC02767 | -1.80737852 | 0.000798 |
| ENSG00000153721 | CNKSR3 | -1.807704557 | 1.78E-12 |
| ENSG00000164182 | NDUFAF2 | -1.811009168 | 1.00E-38 |
| ENSG00000100234 | TIMP3 | -1.812781353 | 1.56E-13 |
| ENSG00000168890 | TMEM150A | -1.813795901 | 1.52E-07 |
| ENSG00000178773 | CPNE7 | -1.815352663 | 1.64E-11 |
| ENSG00000140905 | GCSH | -1.816133664 | 0.000915 |
| ENSG00000242715 | CCDC169 | -1.817538319 | 9.89E-17 |
| ENSG00000171766 | GATM | -1.817717895 | 0.025058 |
| ENSG00000290094 | gene_sourcehavana_ | -1.818424439 | 6.65E-05 |
| ENSG00000146858 | ZC3HAV1L | -1.818793438 | 2.77E-18 |
| ENSG00000186603 | HPDL | -1.819760343 | 2.08E-39 |
| ENSG00000153551 | CMTM7 | -1.821331043 | 3.24E-13 |
| ENSG00000100104 | SRRD | -1.822405497 | 2.78E-57 |
| ENSG00000109089 | CDR2L | -1.825568227 | 0.003659 |

| | | | |
|---|---|---|---|
| ENSG00000155846 | PPARGC1B | -1.829642835 | 5.81E-42 |
| ENSG00000213654 | GPSM3 | -1.829745801 | 1.57E-30 |
| ENSG00000156968 | MPV17L | -1.831644788 | 8.81E-24 |
| ENSG00000278341 | gene_sourcehavana | -1.833306385 | 0.039154 |
| ENSG00000135905 | DOCK10 | -1.834990039 | 2.16E-15 |
| ENSG00000198816 | ZNF358 | -1.83649209 | 0.002613 |
| ENSG00000018280 | SLC11A1 | -1.836573729 | 1.30E-29 |
| ENSG00000177283 | FZD8 | -1.8397622 | 1.72E-13 |
| ENSG00000251602 | MTA1-DT | -1.841043846 | 1.21E-07 |
| ENSG00000158467 | AHCYL2 | -1.842881386 | 4.28E-20 |
| ENSG00000159761 | C16orf86 | -1.843468589 | 5.17E-06 |
| ENSG00000103187 | COTL1 | -1.847859412 | 2.87E-09 |
| ENSG00000274605 | PCCA-DT | -1.84853656 | 6.70E-11 |
| ENSG00000004776 | HSPB6 | -1.853803892 | 0.01356 |
| ENSG00000065882 | TBC1D1 | -1.85530615 | 9.21E-52 |
| ENSG00000164687 | FABP5 | -1.856040006 | 2.75E-62 |
| ENSG00000140323 | DISP2 | -1.856544359 | 0.000822 |
| ENSG00000259673 | IQCH-AS1 | -1.856619374 | 6.63E-08 |
| ENSG00000186854 | TRABD2A | -1.857128397 | 5.45E-11 |
| ENSG00000123427 | EEF1AKMT3 | -1.85837287 | 1.54E-15 |
| ENSG00000289268 | gene_sourcehavana_ | -1.859729289 | 1.65E-05 |
| ENSG00000278619 | MRM1 | -1.859950685 | 8.02E-38 |
| ENSG00000176532 | PRR15 | -1.860777776 | 0.000428 |
| ENSG00000148730 | EIF4EBP2 | -1.861241832 | 4.88E-147 |
| ENSG00000261770 | gene_sourcehavana | -1.862677579 | 0.047143 |
| ENSG00000178078 | STAP2 | -1.863867649 | 9.67E-08 |
| ENSG00000169047 | IRS1 | -1.864070847 | 1.01E-15 |
| ENSG00000110660 | SLC35F2 | -1.864980941 | 3.26E-16 |
| ENSG00000091127 | PUS7 | -1.868393375 | 1.02E-15 |
| ENSG00000008277 | ADAM22 | -1.870461254 | 1.28E-12 |
| ENSG00000235173 | HGH1 | -1.872946351 | 6.12E-14 |
| ENSG00000162836 | ACP6 | -1.875598765 | 1.48E-06 |
| ENSG00000176244 | ACBD7 | -1.87702908 | 2.80E-16 |
| ENSG00000104290 | FZD3 | -1.878469702 | 4.55E-48 |
| ENSG00000108641 | B9D1 | -1.883369038 | 8.99E-22 |
| ENSG00000167799 | NUDT8 | -1.883468946 | 5.59E-12 |
| ENSG00000105227 | PRX | -1.88627801 | 3.60E-10 |
| ENSG00000155792 | DEPTOR | -1.886904843 | 1.68E-11 |
| ENSG00000184675 | AMER1 | -1.889960321 | 1.78E-30 |
| ENSG00000169991 | IFFO2 | -1.890396736 | 9.78E-25 |
| ENSG00000005249 | PRKAR2B | -1.890678486 | 9.64E-80 |
| ENSG00000099985 | OSM | -1.894143021 | 7.98E-36 |
| ENSG00000189337 | KAZN | -1.895965132 | 3.60E-13 |
| ENSG00000248971 | KRT8P46 | -1.898438071 | 0.03378 |
| ENSG00000242193 | CRYZL2P | -1.898716993 | 9.82E-12 |
| ENSG00000115825 | PRKD3 | -1.89879895 | 6.07E-08 |
| ENSG00000172081 | MOB3A | -1.902225951 | 1.46E-49 |
| ENSG00000198765 | SYCP1 | -1.902390171 | 9.68E-08 |
| ENSG00000187210 | GCNT1 | -1.905141432 | 8.33E-48 |

| | | | |
|---|---|---|---|
| ENSG00000130193 | THEM6 | -1.908743421 | 8.91E-37 |
| ENSG00000128596 | CCDC136 | -1.909301986 | 5.28E-05 |
| ENSG00000232063 | gene_sourcehavana | -1.90992715 | 8.17E-31 |
| ENSG00000132530 | XAF1 | -1.91002027 | 7.80E-08 |
| ENSG00000184916 | JAG2 | -1.910129623 | 1.07E-42 |
| ENSG00000230387 | gene_sourcehavana | -1.91036073 | 0.001018 |
| ENSG00000112394 | SLC16A10 | -1.911750032 | 0.004711 |
| ENSG00000136213 | CHST12 | -1.912569918 | 2.64E-66 |
| ENSG00000112238 | PRDM13 | -1.915340674 | 7.12E-05 |
| ENSG00000081985 | IL12RB2 | -1.917974251 | 1.06E-14 |
| ENSG00000286646 | MAP3K5-AS2 | -1.92038256 | 8.25E-07 |
| ENSG00000183807 | FAM162B | -1.920419347 | 1.11E-07 |
| ENSG00000178409 | BEND3 | -1.933032213 | 2.20E-36 |
| ENSG00000120915 | EPHX2 | -1.936454457 | 4.81E-15 |
| ENSG00000106070 | GRB10 | -1.937272276 | 3.80E-101 |
| ENSG00000122778 | KIAA1549 | -1.939089402 | 0.025282 |
| ENSG00000188706 | ZDHHC9 | -1.943336778 | 2.90E-38 |
| ENSG00000121207 | LRAT | -1.94863722 | 0.000544 |
| ENSG00000100321 | SYNGR1 | -1.948878148 | 2.57E-12 |
| ENSG00000280106 | gene_sourcehavana | -1.950052439 | 0.01755 |
| ENSG00000026297 | RNASET2 | -1.951380298 | 6.57E-12 |
| ENSG00000261524 | ABCB10P3 | -1.953870956 | 0.000355 |
| ENSG00000185615 | PDIA2 | -1.95391909 | 2.25E-23 |
| ENSG00000171791 | BCL2 | -1.956656545 | 1.86E-05 |
| ENSG00000136521 | NDUFB5 | -1.957581331 | 3.34E-55 |
| ENSG00000102678 | FGF9 | -1.964764554 | 0.001185 |
| ENSG00000104313 | EYA1 | -1.968276697 | 0.016737 |
| ENSG00000161940 | BCL6B | -1.978183892 | 0.0419 |
| ENSG00000113389 | NPR3 | -1.979105453 | 6.19E-68 |
| ENSG00000189410 | SH2D5 | -1.980504832 | 0.041848 |
| ENSG00000127903 | ZNF835 | -1.983304705 | 2.56E-06 |
| ENSG00000255561 | FDXACB1 | -1.983925918 | 7.27E-09 |
| ENSG00000108262 | GIT1 | -1.985043035 | 4.06E-65 |
| ENSG00000236081 | ELFN1-AS1 | -1.985622115 | 8.81E-12 |
| ENSG00000124783 | SSR1 | -1.992283798 | 6.06E-109 |
| ENSG00000168404 | MLKL | -1.994733987 | 1.94E-31 |
| ENSG00000243989 | ACY1 | -1.99534204 | 0.00036 |
| ENSG00000181163 | NPM1 | -1.996655102 | 1.02E-109 |
| ENSG00000164236 | ANKRD33B | -1.999711249 | 0.0041 |
| ENSG00000061273 | HDAC7 | -2.003138746 | 3.68E-169 |
| ENSG00000225062 | CATIP-AS1 | -2.003563119 | 0.00081 |
| ENSG00000100599 | RIN3 | -2.00627828 | 1.14E-77 |
| ENSG00000173638 | SLC19A1 | -2.008483126 | 1.53E-65 |
| ENSG00000136379 | ABHD17C | -2.010333692 | 1.23E-17 |
| ENSG00000136720 | HS6ST1 | -2.014697513 | 1.05E-51 |
| ENSG00000100605 | ITPK1 | -2.015817479 | 1.48E-90 |
| ENSG00000138449 | SLC40A1 | -2.018085906 | 3.14E-119 |
| ENSG00000164237 | CMBL | -2.026995878 | 2.65E-118 |
| ENSG00000130669 | PAK4 | -2.035071661 | 1.88E-113 |

| | | | |
|---|---|---|---|
| ENSG00000186523 | FAM86B1 | -2.038810625 | 2.54E-11 |
| ENSG00000158089 | GALNT14 | -2.041845637 | 2.05E-32 |
| ENSG00000245848 | CEBPA | -2.053626222 | 4.02E-09 |
| ENSG00000274099 | ABCB10P1 | -2.054099176 | 4.30E-06 |
| ENSG00000172819 | RARG | -2.05564806 | 1.07E-16 |
| ENSG00000168497 | CAVIN2 | -2.056021455 | 1.49E-06 |
| ENSG00000159208 | CIART | -2.058921474 | 5.96E-23 |
| ENSG00000134508 | CABLES1 | -2.060740046 | 1.36E-41 |
| ENSG00000198885 | ITPRIPL1 | -2.061619156 | 1.30E-05 |
| ENSG00000143119 | CD53 | -2.062343403 | 0.000176 |
| ENSG00000155380 | SLC16A1 | -2.062361294 | 5.29E-23 |
| ENSG00000213468 | FIRRE | -2.063376881 | 7.70E-25 |
| ENSG00000133740 | E2F5 | -2.063637055 | 7.17E-19 |
| ENSG00000288829 | gene_sourcehavana_ | -2.068304198 | 0.038822 |
| ENSG00000162733 | DDR2 | -2.074241872 | 0.025197 |
| ENSG00000184221 | OLIG1 | -2.075435887 | 7.72E-34 |
| ENSG00000234494 | SP2-AS1 | -2.08005246 | 1.77E-06 |
| ENSG00000140199 | SLC12A6 | -2.083030523 | 4.41E-130 |
| ENSG00000129675 | ARHGEF6 | -2.084170242 | 2.69E-87 |
| ENSG00000260053 | ABCB10P4 | -2.090014231 | 1.69E-09 |
| ENSG00000287166 | gene_sourcehavana_ | -2.091201384 | 0.035122 |
| ENSG00000174137 | FAM53A | -2.095599424 | 0.002777 |
| ENSG00000236915 | CLCA4-AS1 | -2.096629914 | 0.001234 |
| ENSG00000223749 | MIR503HG | -2.103250036 | 2.21E-06 |
| ENSG00000052749 | RRP12 | -2.108003202 | 1.46E-112 |
| ENSG00000123191 | ATP7B | -2.115563041 | 9.54E-15 |
| ENSG00000175600 | SUGCT | -2.118928314 | 6.85E-09 |
| ENSG00000161405 | IKZF3 | -2.123477854 | 4.52E-61 |
| ENSG00000213881 | NPM1P6 | -2.12713862 | 0.002052 |
| ENSG00000004139 | SARM1 | -2.132257967 | 1.41E-07 |
| ENSG00000184831 | APOO | -2.140071503 | 2.51E-21 |
| ENSG00000157613 | CREB3L1 | -2.140557654 | 1.30E-45 |
| ENSG00000224167 | LINC01357 | -2.140729483 | 0.036414 |
| ENSG00000139926 | FRMD6 | -2.144547954 | 0.007147 |
| ENSG00000143499 | SMYD2 | -2.14829482 | 1.59E-40 |
| ENSG00000198246 | SLC29A3 | -2.149849326 | 4.93E-17 |
| ENSG00000249353 | NPM1P27 | -2.151506064 | 3.18E-95 |
| ENSG00000188372 | ZP3 | -2.151991992 | 3.03E-08 |
| ENSG00000262768 | gene_sourcehavana | -2.1529789 | 0.001201 |
| ENSG00000225159 | NPM1P39 | -2.154814791 | 3.30E-07 |
| ENSG00000169174 | PCSK9 | -2.157207132 | 2.23E-16 |
| ENSG00000198720 | ANKRD13B | -2.163576798 | 3.15E-28 |
| ENSG00000152518 | ZFP36L2 | -2.172165347 | 6.99E-168 |
| ENSG00000116711 | PLA2G4A | -2.172853753 | 5.58E-14 |
| ENSG00000170899 | GSTA4 | -2.174013269 | 6.05E-10 |
| ENSG00000023892 | DEF6 | -2.177473089 | 1.91E-53 |
| ENSG00000257913 | DDN-AS1 | -2.180646424 | 5.07E-06 |
| ENSG00000174004 | NRROS | -2.180742329 | 3.11E-77 |
| ENSG00000070404 | FSTL3 | -2.181086603 | 6.87E-29 |

| | | | |
|---|---|---|---|
| ENSG00000186615 | KTN1-AS1 | -2.18123159 | 3.88E-13 |
| ENSG00000178695 | KCTD12 | -2.183106036 | 0.008414 |
| ENSG00000185252 | ZNF74 | -2.188855771 | 1.08E-28 |
| ENSG00000263961 | RHEX | -2.192840733 | 3.85E-21 |
| ENSG00000138835 | RGS3 | -2.193701199 | 1.28E-42 |
| ENSG00000289810 | gene_sourcehavana | -2.194410989 | 2.03E-05 |
| ENSG00000123384 | LRP1 | -2.197490724 | 4.91E-10 |
| ENSG00000174500 | GCSAM | -2.211643251 | 1.12E-08 |
| ENSG00000162614 | NEXN | -2.213434908 | 0.031349 |
| ENSG00000073792 | IGF2BP2 | -2.223902227 | 4.03E-106 |
| ENSG00000167701 | GPT | -2.228505567 | 5.30E-14 |
| ENSG00000111728 | ST8SIA1 | -2.229490976 | 4.22E-08 |
| ENSG00000168477 | TNXB | -2.238106509 | 1.35E-09 |
| ENSG00000150764 | DIXDC1 | -2.240399297 | 6.28E-62 |
| ENSG00000203943 | SAMD13 | -2.244701925 | 0.023872 |
| ENSG00000169247 | SH3TC2 | -2.246633996 | 1.90E-19 |
| ENSG00000173991 | TCAP | -2.25890529 | 2.29E-05 |
| ENSG00000196482 | ESRRG | -2.259431717 | 6.94E-81 |
| ENSG00000266208 | GJD3-AS1 | -2.263763808 | 1.01E-13 |
| ENSG00000103184 | SEC14L5 | -2.270569644 | 0.037995 |
| ENSG00000134874 | DZIP1 | -2.27183722 | 0.010687 |
| ENSG00000275793 | RIMBP3 | -2.271968088 | 0.046407 |
| ENSG00000272476 | gene_sourcehavana | -2.276136932 | 0.014104 |
| ENSG00000214050 | FBXO16 | -2.278602072 | 8.88E-05 |
| ENSG00000243224 | TWF2-DT | -2.281233961 | 0.016397 |
| ENSG00000186648 | CARMIL3 | -2.28195337 | 8.33E-10 |
| ENSG00000171608 | PIK3CD | -2.282520274 | 0.0326 |
| ENSG00000228594 | FNDC10 | -2.282685929 | 1.08E-08 |
| ENSG00000236305 | SLC12A9-AS1 | -2.288741906 | 0.010449 |
| ENSG00000138613 | APH1B | -2.293014559 | 1.23E-26 |
| ENSG00000234753 | FOXP4-AS1 | -2.293028526 | 6.04E-05 |
| ENSG00000271270 | TMCC1-DT | -2.294192954 | 8.61E-15 |
| ENSG00000198429 | ZNF69 | -2.294858555 | 3.92E-06 |
| ENSG00000095539 | SEMA4G | -2.296683625 | 0.031553 |
| ENSG00000272711 | HK2-DT | -2.299554847 | 2.41E-06 |
| ENSG00000135414 | GDF11 | -2.30136006 | 1.26E-05 |
| ENSG00000197965 | MPZL1 | -2.302638538 | 2.94E-50 |
| ENSG00000272129 | gene_sourcehavana | -2.305437209 | 0.000354 |
| ENSG00000135426 | TESPA1 | -2.309946422 | 1.39E-10 |
| ENSG00000162520 | SYNC | -2.310932839 | 3.84E-18 |
| ENSG00000159399 | HK2 | -2.312579272 | 4.12E-19 |
| ENSG00000185905 | C16orf54 | -2.314370413 | 1.74E-05 |
| ENSG00000107104 | KANK1 | -2.322828372 | 1.08E-30 |
| ENSG00000228956 | SATB1-AS1 | -2.331039797 | 0.004024 |
| ENSG00000181754 | AMIGO1 | -2.33210313 | 0.013095 |
| ENSG00000175445 | LPL | -2.332274411 | 2.19E-30 |
| ENSG00000138030 | KHK | -2.33513002 | 5.42E-51 |
| ENSG00000125285 | SOX21 | -2.335867476 | 2.91E-07 |
| ENSG00000131378 | RFTN1 | -2.33806302 | 1.66E-23 |

| | | | |
|---|---|---|---|
| ENSG00000280202 | gene_sourcehavana | -2.338444158 | 2.97E-18 |
| ENSG00000138028 | CGREF1 | -2.34127249 | 3.08E-10 |
| ENSG00000110934 | BIN2 | -2.342577244 | 1.07E-75 |
| ENSG00000138395 | CDK15 | -2.343946 | 3.45E-43 |
| ENSG00000006740 | ARHGAP44 | -2.344480549 | 0.021473 |
| ENSG00000087253 | LPCAT2 | -2.347564439 | 1.04E-49 |
| ENSG00000120949 | TNFRSF8 | -2.348023191 | 6.41E-36 |
| ENSG00000091409 | ITGA6 | -2.349229996 | 0.031169 |
| ENSG00000124766 | SOX4 | -2.349624626 | 4.13E-16 |
| ENSG00000043462 | LCP2 | -2.350562123 | 3.98E-66 |
| ENSG00000181274 | FRAT2 | -2.352950085 | 1.91E-97 |
| ENSG00000130475 | FCHO1 | -2.355584503 | 4.55E-51 |
| ENSG00000063241 | ISOC2 | -2.357526789 | 1.12E-16 |
| ENSG00000137124 | ALDH1B1 | -2.363719399 | 2.10E-14 |
| ENSG00000229536 | LINC02572 | -2.365888949 | 0.005932 |
| ENSG00000033170 | FUT8 | -2.367514759 | 3.19E-52 |
| ENSG00000106348 | IMPDH1 | -2.368285509 | 5.60E-84 |
| ENSG00000148219 | ASTN2 | -2.372637212 | 1.73E-17 |
| ENSG00000138439 | FAM117B | -2.372975387 | 1.48E-08 |
| ENSG00000158286 | RNF207 | -2.374290895 | 1.49E-11 |
| ENSG00000179431 | FJX1 | -2.377106481 | 2.12E-41 |
| ENSG00000138080 | EMILIN1 | -2.377799869 | 3.68E-08 |
| ENSG00000016391 | CHDH | -2.386860549 | 6.55E-25 |
| ENSG00000119699 | TGFB3 | -2.388205115 | 2.16E-15 |
| ENSG00000184584 | STING1 | -2.393130197 | 6.39E-67 |
| ENSG00000172995 | ARPP21 | -2.400433606 | 2.60E-05 |
| ENSG00000185664 | PMEL | -2.400456145 | 0.025492 |
| ENSG00000241399 | CD302 | -2.403383063 | 1.21E-05 |
| ENSG00000147180 | ZNF711 | -2.403492933 | 1.41E-32 |
| ENSG00000214694 | ARHGEF33 | -2.406487287 | 5.17E-15 |
| ENSG00000278041 | gene_sourcehavana | -2.408642879 | 2.22E-05 |
| ENSG00000105808 | RASA4 | -2.413888255 | 5.78E-05 |
| ENSG00000120669 | SOHLH2 | -2.418336462 | 0.002206 |
| ENSG00000182747 | SLC35D3 | -2.420073173 | 1.01E-05 |
| ENSG00000120645 | IQSEC3 | -2.422613428 | 5.31E-17 |
| ENSG00000197381 | ADARB1 | -2.422825548 | 5.78E-57 |
| ENSG00000235172 | LINC01366 | -2.423055146 | 1.83E-59 |
| ENSG00000145002 | FAM86B2 | -2.423328577 | 7.12E-08 |
| ENSG00000259020 | gene_sourcehavana | -2.428631543 | 0.046058 |
| ENSG00000135114 | OASL | -2.437470753 | 4.23E-05 |
| ENSG00000130881 | LRP3 | -2.438375999 | 8.19E-06 |
| ENSG00000101977 | MCF2 | -2.438794555 | 1.74E-32 |
| ENSG00000123213 | NLN | -2.443006585 | 1.27E-94 |
| ENSG00000271856 | LINC01215 | -2.449504911 | 4.81E-95 |
| ENSG00000168140 | VASN | -2.451283896 | 3.10E-05 |
| ENSG00000211829 | TRDC | -2.45193332 | 0.014652 |
| ENSG00000133895 | MEN1 | -2.45197387 | 1.50E-144 |
| ENSG00000071575 | TRIB2 | -2.452630465 | 1.12E-12 |
| ENSG00000250899 | gene_sourcehavana | -2.454632525 | 1.17E-10 |

| | | | |
|---|---|---|---|
| ENSG00000167123 | CERCAM | -2.455374955 | 4.29E-16 |
| ENSG00000227268 | KLLN | -2.459943228 | 0.000899 |
| ENSG00000234964 | FABP5P7 | -2.460331733 | 0.003117 |
| ENSG00000198933 | TBKBP1 | -2.465967384 | 7.10E-12 |
| ENSG00000198648 | STK39 | -2.469031091 | 2.86E-07 |
| ENSG00000205336 | ADGRG1 | -2.469253495 | 1.28E-10 |
| ENSG00000215481 | BCRP3 | -2.488124417 | 0.041889 |
| ENSG00000182118 | FAM89A | -2.496755347 | 9.82E-97 |
| ENSG00000112297 | CRYBG1 | -2.502600596 | 1.64E-265 |
| ENSG00000146700 | SSC4D | -2.505139862 | 0.001907 |
| ENSG00000207445 | SNORD15B | -2.50767677 | 0.035925 |
| ENSG00000215086 | NPM1P24 | -2.507819922 | 2.03E-13 |
| ENSG00000157554 | ERG | -2.507854356 | 2.08E-08 |
| ENSG00000272338 | gene_sourcehavana | -2.510963391 | 0.009377 |
| ENSG00000288860 | gene_sourcehavana_ | -2.51127369 | 1.92E-05 |
| ENSG00000185745 | IFIT1 | -2.524714746 | 1.21E-21 |
| ENSG00000124743 | KLHL31 | -2.526933526 | 6.71E-05 |
| ENSG00000175414 | ARL10 | -2.534771068 | 1.89E-37 |
| ENSG00000170667 | RASA4B | -2.534980621 | 5.19E-07 |
| ENSG00000119866 | BCL11A | -2.535909199 | 2.15E-07 |
| ENSG00000289157 | gene_sourcehavana_ | -2.553613116 | 1.58E-06 |
| ENSG00000288573 | gene_sourcehavana | -2.562496954 | 0.000227 |
| ENSG00000288016 | gene_sourcehavana_ | -2.567732225 | 0.010792 |
| ENSG00000076356 | PLXNA2 | -2.570987044 | 9.50E-34 |
| ENSG00000105514 | RAB3D | -2.578919323 | 1.72E-06 |
| ENSG00000178150 | ZNF114 | -2.595337194 | 2.52E-06 |
| ENSG00000168765 | GSTM4 | -2.601820272 | 9.05E-57 |
| ENSG00000215788 | TNFRSF25 | -2.606818476 | 3.62E-09 |
| ENSG00000272913 | LINC03052 | -2.607545107 | 1.64E-06 |
| ENSG00000115705 | TPO | -2.615052033 | 5.40E-32 |
| ENSG00000213366 | GSTM2 | -2.61804205 | 9.63E-19 |
| ENSG00000196502 | SULT1A1 | -2.623651239 | 0.012274 |
| ENSG00000267681 | gene_sourcehavana | -2.660734469 | 0.018645 |
| ENSG00000107554 | DNMBP | -2.671348412 | 9.71E-11 |
| ENSG00000141744 | PNMT | -2.676471736 | 1.07E-30 |
| ENSG00000111186 | WNT5B | -2.67692959 | 0.022041 |
| ENSG00000143333 | RGS16 | -2.683418383 | 1.87E-34 |
| ENSG00000105889 | STEAP1B | -2.687870373 | 2.12E-25 |
| ENSG00000262712 | gene_sourcehavana | -2.695392195 | 1.82E-07 |
| ENSG00000010319 | SEMA3G | -2.698546365 | 0.010446 |
| ENSG00000136111 | TBC1D4 | -2.700386156 | 3.36E-26 |
| ENSG00000011590 | ZBTB32 | -2.700991749 | 3.70E-12 |
| ENSG00000050555 | LAMC3 | -2.708678915 | 0.001843 |
| ENSG00000092067 | CEBPE | -2.710700311 | 2.31E-19 |
| ENSG00000105649 | RAB3A | -2.719719297 | 7.74E-10 |
| ENSG00000198758 | EPS8L3 | -2.720092657 | 1.14E-08 |
| ENSG00000184185 | KCNJ12 | -2.729047817 | 4.68E-41 |
| ENSG00000213104 | NPM1P46 | -2.7314844 | 1.05E-14 |
| ENSG00000127863 | TNFRSF19 | -2.7318523 | 1.83E-16 |

| | | | |
|---|---|---|---|
| ENSG00000274737 | gene_sourcehavana | -2.733612524 | 0.035849 |
| ENSG00000185760 | KCNQ5 | -2.733674923 | 6.41E-10 |
| ENSG00000090932 | DLL3 | -2.741190095 | 0.000154 |
| ENSG00000276527 | gene_sourcehavana | -2.76707649 | 2.81E-07 |
| ENSG00000131773 | KHDRBS3 | -2.782221788 | 7.16E-08 |
| ENSG00000161055 | SCGB3A1 | -2.791101391 | 0.032697 |
| ENSG00000101695 | RNF125 | -2.792928127 | 9.27E-60 |
| ENSG00000232872 | CTAGE3P | -2.817820523 | 0.01522 |
| ENSG00000161243 | FBXO27 | -2.81784646 | 2.04E-09 |
| ENSG00000036672 | USP2 | -2.819128885 | 1.05E-05 |
| ENSG00000231050 | GNB1-DT | -2.821208149 | 0.045567 |
| ENSG00000163013 | FBXO41 | -2.833720297 | 0.000264 |
| ENSG00000131746 | TNS4 | -2.836833522 | 0.001214 |
| ENSG00000129667 | RHBDF2 | -2.84281163 | 4.82E-05 |
| ENSG00000140807 | NKD1 | -2.857097003 | 0.005944 |
| ENSG00000104870 | FCGRT | -2.865594036 | 5.77E-07 |
| ENSG00000100077 | GRK3 | -2.876973494 | 1.86E-90 |
| ENSG00000224728 | IMPDH1P8 | -2.884076213 | 0.007388 |
| ENSG00000245248 | USP2-AS1 | -2.889584451 | 4.83E-11 |
| ENSG00000072110 | ACTN1 | -2.89093673 | 4.23E-12 |
| ENSG00000136425 | CIB2 | -2.893473944 | 0.000949 |
| ENSG00000152990 | ADGRA3 | -2.89374612 | 1.49E-27 |
| ENSG00000154328 | NEIL2 | -2.895050261 | 7.60E-17 |
| ENSG00000136286 | MYO1G | -2.896892481 | 1.23E-74 |
| ENSG00000223573 | TINCR | -2.909345566 | 2.57E-72 |
| ENSG00000138336 | TET1 | -2.910446082 | 3.58E-10 |
| ENSG00000178821 | TMEM52 | -2.915913349 | 5.54E-10 |
| ENSG00000283199 | C13orf46 | -2.936628095 | 0.002446 |
| ENSG00000288771 | gene_sourcehavana_ | -2.957521526 | 2.49E-07 |
| ENSG00000188613 | NANOS1 | -2.975556154 | 1.65E-05 |
| ENSG00000090382 | LYZ | -2.975856372 | 6.08E-06 |
| ENSG00000103811 | CTSH | -2.978183899 | 6.40E-47 |
| ENSG00000243414 | TICAM2 | -2.980163273 | 0.006865 |
| ENSG00000265190 | ANXA8 | -2.983460687 | 0.00694 |
| ENSG00000170891 | CYTL1 | -2.986204312 | 8.04E-22 |
| ENSG00000141753 | IGFBP4 | -3.002193299 | 1.79E-23 |
| ENSG00000248890 | HHIP-AS1 | -3.004010185 | 0.009669 |
| ENSG00000197977 | ELOVL2 | -3.014448562 | 1.49E-29 |
| ENSG00000054219 | LY75 | -3.023045803 | 5.49E-06 |
| ENSG00000157470 | FAM81A | -3.03946673 | 2.33E-17 |
| ENSG00000261218 | gene_sourcehavana | -3.046093079 | 1.91E-14 |
| ENSG00000181418 | DDN | -3.053866676 | 4.73E-15 |
| ENSG00000196189 | SEMA4A | -3.056252138 | 9.99E-07 |
| ENSG00000267058 | gene_sourcehavana | -3.076702248 | 0.014619 |
| ENSG00000145217 | SLC26A1 | -3.081212352 | 1.31E-05 |
| ENSG00000106236 | NPTX2 | -3.090187826 | 1.01E-12 |
| ENSG00000251002 | TRD-AS1 | -3.103138076 | 2.28E-14 |
| ENSG00000155926 | SLA | -3.111242379 | 4.48E-11 |
| ENSG00000160867 | FGFR4 | -3.118707256 | 4.49E-14 |

| | | | |
|---|---|---|---|
| ENSG00000053747 | LAMA3 | -3.135333249 | 0.049114 |
| ENSG00000160185 | UBASH3A | -3.142683997 | 1.52E-21 |
| ENSG00000124507 | PACSIN1 | -3.148497392 | 0.000754 |
| ENSG00000134986 | NREP | -3.157463829 | 2.58E-11 |
| ENSG00000273893 | gene_sourcehavana | -3.161393834 | 0.00545 |
| ENSG00000272006 | gene_sourcehavana | -3.170643224 | 0.016154 |
| ENSG00000144712 | CAND2 | -3.175465505 | 0.023198 |
| ENSG00000180264 | ADGRD2 | -3.180452756 | 1.07E-06 |
| ENSG00000274536 | MIR223HG | -3.180597799 | 2.15E-33 |
| ENSG00000288990 | gene_sourcehavana_ | -3.182166179 | 1.64E-19 |
| ENSG00000163154 | TNFAIP8L2 | -3.19670337 | 0.001318 |
| ENSG00000236809 | SNX25P1 | -3.200579571 | 0.007819 |
| ENSG00000287497 | gene_sourcehavana_ | -3.203928567 | 1.76E-12 |
| ENSG00000104951 | IL4I1 | -3.211878352 | 0.020692 |
| ENSG00000179846 | NKPD1 | -3.228705991 | 3.31E-69 |
| ENSG00000172828 | CES3 | -3.235060333 | 0.001362 |
| ENSG00000198682 | PAPSS2 | -3.249252108 | 8.20E-06 |
| ENSG00000286214 | COPG2IT1 | -3.253212869 | 0.005192 |
| ENSG00000163453 | IGFBP7 | -3.259946772 | 5.17E-08 |
| ENSG00000188095 | MESP2 | -3.27494987 | 1.56E-06 |
| ENSG00000176595 | KBTBD11 | -3.298395597 | 1.14E-12 |
| ENSG00000104432 | IL7 | -3.317821048 | 5.35E-68 |
| ENSG00000179796 | LRRC3B | -3.322007372 | 0.013208 |
| ENSG00000172572 | PDE3A | -3.355170389 | 1.03E-28 |
| ENSG00000081277 | PKP1 | -3.360122026 | 9.56E-14 |
| ENSG00000261618 | LINC02605 | -3.361227001 | 0.00215 |
| ENSG00000101311 | FERMT1 | -3.363541946 | 3.32E-09 |
| ENSG00000285844 | KCNQ5-DT | -3.370810837 | 9.03E-05 |
| ENSG00000164161 | HHIP | -3.373838273 | 0.000467 |
| ENSG00000104833 | TUBB4A | -3.38689731 | 0.00102 |
| ENSG00000113532 | ST8SIA4 | -3.394652526 | 4.53E-43 |
| ENSG00000176387 | HSD11B2 | -3.398606409 | 1.05E-34 |
| ENSG00000183034 | OTOP2 | -3.412832253 | 1.78E-28 |
| ENSG00000132915 | PDE6A | -3.46495412 | 1.76E-05 |
| ENSG00000239911 | PRKAG2-AS1 | -3.472959791 | 0.008249 |
| ENSG00000214787 | MS4A4E | -3.490914696 | 3.44E-07 |
| ENSG00000050628 | PTGER3 | -3.507109718 | 1.24E-45 |
| ENSG00000179399 | GPC5 | -3.559928014 | 1.68E-52 |
| ENSG00000289937 | gene_sourcehavana_ | -3.582504529 | 0.039439 |
| ENSG00000253152 | IGLV3-17 | -3.583132652 | 0.045328 |
| ENSG00000188763 | FZD9 | -3.588885022 | 6.00E-14 |
| ENSG00000188883 | KLRG2 | -3.598155067 | 2.77E-14 |
| ENSG00000235162 | C12orf75 | -3.608074698 | 2.96E-117 |
| ENSG00000121236 | TRIM6 | -3.618247214 | 6.36E-60 |
| ENSG00000180066 | LINC02870 | -3.639557928 | 0.040892 |
| ENSG00000007264 | MATK | -3.659108326 | 1.53E-10 |
| ENSG00000285744 | gene_sourcehavana | -3.662566427 | 0.029224 |
| ENSG00000145832 | SLC25A48 | -3.663556813 | 1.36E-17 |
| ENSG00000260997 | gene_sourcehavana | -3.665101367 | 4.85E-06 |

| | | | |
|---|---|---|---|
| ENSG00000083454 | P2RX5 | -3.675664901 | 9.09E-14 |
| ENSG00000186152 | LILRP1 | -3.684719776 | 5.00E-07 |
| ENSG00000272989 | LINC02012 | -3.688354852 | 0.002169 |
| ENSG00000047662 | FAM184B | -3.7034267 | 1.29E-71 |
| ENSG00000249863 | gene_sourcehavana | -3.705749405 | 4.21E-07 |
| ENSG00000224786 | CETN4P | -3.713553158 | 0.000371 |
| ENSG00000164362 | TERT | -3.734491708 | 1.23E-51 |
| ENSG00000132938 | MTUS2 | -3.762759112 | 2.52E-06 |
| ENSG00000237359 | gene_sourcehavana | -3.767754086 | 0.009932 |
| ENSG00000117643 | MAN1C1 | -3.801573048 | 9.51E-91 |
| ENSG00000172794 | RAB37 | -3.850190377 | 0.022665 |
| ENSG00000180767 | CHST13 | -3.854264894 | 0.023601 |
| ENSG00000185198 | PRSS57 | -3.865821992 | 1.88E-24 |
| ENSG00000262188 | LINC01978 | -3.867814584 | 2.10E-20 |
| ENSG00000205704 | SMIM45 | -3.868318274 | 0.019068 |
| ENSG00000157657 | ZNF618 | -3.86855804 | 1.03E-07 |
| ENSG00000166743 | ACSM1 | -3.870504254 | 0.005065 |
| ENSG00000160678 | S100A1 | -3.88029579 | 0.002741 |
| ENSG00000157890 | MEGF11 | -3.881402921 | 0.024246 |
| ENSG00000157111 | TMEM171 | -3.909401581 | 0.002424 |
| ENSG00000204475 | NCR3 | -3.921843129 | 7.07E-09 |
| ENSG00000250072 | SH3TC2-DT | -3.928459919 | 0.004515 |
| ENSG00000142552 | RCN3 | -3.932719423 | 1.34E-08 |
| ENSG00000248290 | TNXA | -3.936512937 | 0.001957 |
| ENSG00000251287 | ALG1L2 | -3.937954749 | 0.019902 |
| ENSG00000169255 | B3GALNT1 | -3.96357587 | 5.24E-33 |
| ENSG00000132359 | RAP1GAP2 | -3.988257137 | 2.50E-13 |
| ENSG00000292333 | P2RY8 | -4.009138918 | 0.010012 |
| ENSG00000068831 | RASGRP2 | -4.009177067 | 0.01163 |
| ENSG00000170456 | DENND5B | -4.038729297 | 0.001337 |
| ENSG00000250167 | gene_sourcehavana | -4.041382823 | 1.64E-07 |
| ENSG00000244040 | IL12A-AS1 | -4.070473949 | 0.010411 |
| ENSG00000183837 | PNMA3 | -4.085473505 | 0.002189 |
| ENSG00000164647 | STEAP1 | -4.10899539 | 0.009683 |
| ENSG00000215910 | C1orf167 | -4.134680418 | 4.29E-08 |
| ENSG00000163520 | FBLN2 | -4.148897332 | 0.00065 |
| ENSG00000020181 | ADGRA2 | -4.182498834 | 4.92E-05 |
| ENSG00000286856 | gene_sourcehavana_ | -4.188354808 | 0.038287 |
| ENSG00000101638 | ST8SIA5 | -4.206847339 | 0.044877 |
| ENSG00000232682 | gene_sourcehavana | -4.209164583 | 2.84E-09 |
| ENSG00000063127 | SLC6A16 | -4.223785503 | 0.030848 |
| ENSG00000283122 | HYMAI | -4.233175417 | 0.042247 |
| ENSG00000158220 | ESYT3 | -4.254500038 | 0.031349 |
| ENSG00000230314 | ELOVL2-AS1 | -4.281528801 | 0.007043 |
| ENSG00000272456 | gene_sourcehavana | -4.283285833 | 0.006614 |
| ENSG00000198915 | RASGEF1A | -4.311706089 | 2.09E-06 |
| ENSG00000205861 | PCOTH | -4.327908339 | 2.11E-10 |
| ENSG00000009790 | TRAF3IP3 | -4.328389134 | 3.30E-12 |
| ENSG00000172232 | AZU1 | -4.330304209 | 3.44E-09 |

| | | | |
|---|---|---|---|
| ENSG00000214992 | AKAP17BP | -4.337819186 | 0.022948 |
| ENSG00000180044 | C3orf80 | -4.363182959 | 3.70E-28 |
| ENSG00000149922 | TBX6 | -4.364752867 | 1.69E-05 |
| ENSG00000258857 | gene_sourcehavana | -4.370642557 | 0.025605 |
| ENSG00000125845 | BMP2 | -4.373819267 | 0.031781 |
| ENSG00000254352 | C4orf46P3 | -4.382171254 | 0.045567 |
| ENSG00000258365 | gene_sourcehavana | -4.404722154 | 0.019846 |
| ENSG00000184232 | OAF | -4.414443576 | 3.03E-13 |
| ENSG00000278983 | gene_sourcehavana | -4.430426887 | 0.018583 |
| ENSG00000181215 | C4orf50 | -4.436177835 | 0.017807 |
| ENSG00000245522 | LINC02709 | -4.465819216 | 0.017675 |
| ENSG00000131634 | TMEM204 | -4.468810114 | 0.002479 |
| ENSG00000181085 | MAPK15 | -4.5002702 | 0.002499 |
| ENSG00000183134 | PTGDR2 | -4.501254059 | 1.57E-05 |
| ENSG00000203546 | gene_sourcehavana | -4.514157751 | 0.01848 |
| ENSG00000108813 | DLX4 | -4.51599257 | 0.01374 |
| ENSG00000131771 | PPP1R1B | -4.559833626 | 6.88E-46 |
| ENSG00000290074 | gene_sourcehavana_ | -4.615156705 | 0.043716 |
| ENSG00000136378 | ADAMTS7 | -4.659276881 | 0.012965 |
| ENSG00000285571 | gene_sourcehavana | -4.669934998 | 0.038129 |
| ENSG00000283632 | EXOC3L2 | -4.69008961 | 0.040466 |
| ENSG00000266601 | gene_sourcehavana | -4.72368683 | 0.044867 |
| ENSG00000246283 | TTBK2-AS1 | -4.738144326 | 0.035604 |
| ENSG00000141506 | PIK3R5 | -4.752188893 | 0.030825 |
| ENSG00000242767 | ZBTB20-AS4 | -4.7543545 | 0.033054 |
| ENSG00000146151 | HMGCLL1 | -4.778893297 | 0.029691 |
| ENSG00000188822 | CNR2 | -4.779333875 | 0.010327 |
| ENSG00000267272 | LINC01140 | -4.779391802 | 0.006085 |
| ENSG00000161992 | PRR35 | -4.790786575 | 0.028275 |
| ENSG00000258461 | gene_sourceensembl | -4.856385633 | 0.023592 |
| ENSG00000121053 | EPX | -4.858588575 | 0.032263 |
| ENSG00000046653 | GPM6B | -4.868039357 | 0.048067 |
| ENSG00000128011 | LRFN1 | -4.876696316 | 2.00E-08 |
| ENSG00000279667 | gene_sourcehavana | -4.896396573 | 0.028155 |
| ENSG00000082482 | KCNK2 | -4.910894244 | 0.019399 |
| ENSG00000154065 | ANKRD29 | -4.942170913 | 0.022783 |
| ENSG00000260571 | BNIP3P5 | -4.942632352 | 0.026785 |
| ENSG00000288749 | gene_sourcehavana_ | -4.963106223 | 0.016677 |
| ENSG00000256802 | LCIIAR | -4.971673107 | 0.016288 |
| ENSG00000165449 | SLC16A9 | -4.975838368 | 0.000329 |
| ENSG00000211662 | IGLV3-21 | -4.993875177 | 0.034334 |
| ENSG00000132622 | HSPA12B | -5.015733842 | 4.05E-26 |
| ENSG00000183629 | GOLGA8G | -5.062153643 | 0.012491 |
| ENSG00000240065 | PSMB9 | -5.076820575 | 6.90E-15 |
| ENSG00000258603 | gene_sourcehavana | -5.086606863 | 0.011416 |
| ENSG00000183153 | GJD3 | -5.134258216 | 0.013953 |
| ENSG00000270953 | gene_sourcehavana | -5.146413678 | 0.010687 |
| ENSG00000197768 | STPG3 | -5.155616181 | 0.009862 |
| ENSG00000273186 | gene_sourcehavana | -5.164814955 | 0.009126 |

| | | | |
|---|---|---|---|
| ENSG00000065320 | NTN1 | -5.20191386 | 5.78E-08 |
| ENSG00000109846 | CRYAB | -5.210254737 | 0.001363 |
| ENSG00000100385 | IL2RB | -5.367106161 | 5.46E-28 |
| ENSG00000110076 | NRXN2 | -5.375507571 | 1.85E-07 |
| ENSG00000157510 | AFAP1L1 | -5.377306242 | 0.007636 |
| ENSG00000125878 | TCF15 | -5.402727698 | 0.004922 |
| ENSG00000129951 | PLPPR3 | -5.417488515 | 1.12E-38 |
| ENSG00000145192 | AHSG | -5.476265253 | 0.003019 |
| ENSG00000174099 | MSRB3 | -5.526948611 | 0.002368 |
| ENSG00000289881 | gene_sourcehavana_ | -5.595558857 | 0.002339 |
| ENSG00000139537 | CCDC65 | -5.599417677 | 0.001699 |
| ENSG00000164604 | GPR85 | -5.614382942 | 3.03E-33 |
| ENSG00000224729 | PCOLCE-AS1 | -5.64206497 | 0.001431 |
| ENSG00000219607 | PPP1R3G | -5.687913947 | 3.93E-06 |
| ENSG00000167641 | PPP1R14A | -5.688359307 | 5.24E-06 |
| ENSG00000163879 | DNALI1 | -5.877458 | 0.00058 |
| ENSG00000283095 | gene_sourcehavana | -5.907861103 | 0.000824 |
| ENSG00000174059 | CD34 | -6.030248457 | 1.40E-09 |
| ENSG00000115414 | FN1 | -6.100516243 | 0.017432 |
| ENSG00000115919 | KYNU | -6.157433028 | 0.000206 |
| ENSG00000140835 | CHST4 | -6.186344404 | 0.000114 |
| ENSG00000172543 | CTSW | -6.637498047 | 1.87E-05 |
| ENSG00000198944 | SOWAHA | -6.673055282 | 1.40E-05 |
| ENSG00000228401 | gene_sourcehavana | -6.839303241 | 5.23E-06 |
| ENSG00000005844 | ITGAL | -7.527175712 | 1.83E-08 |
| ENSG00000102048 | ASB9 | -7.626481417 | 2.40E-08 |
| ENSG00000143147 | GPR161 | -8.53289955 | 1.23E-10 |
